# Nucleoporin93 (Nup93) Limits Yap Activity to Prevent Endothelial Cell Senescence

**DOI:** 10.1101/2023.11.10.566598

**Authors:** Tung D. Nguyen, Mihir K. Rao, Shaiva P. Dhyani, Justin M. Banks, Michael A. Winek, Julka Michalkiewicz, Monica Y. Lee

## Abstract

Endothelial cells (ECs) form the innermost lining of the vasculature and serve a pivotal role in preventing age-related vascular disease. Endothelial health relies on the proper nucleocytoplasmic shuttling of transcription factors via nuclear pore complexes (NPCs). Emerging studies report NPC degradation with natural aging, suggesting impaired nucleocytoplasmic transport in age-related EC dysfunction. We herein identify nucleoporin93 (Nup93), a crucial structural NPC protein, as an indispensable player for vascular protection. Endothelial Nup93 protein levels are significantly reduced in the vasculature of aged mice, paralleling observations of Nup93 loss when using *in vitro* models of endothelial aging. Mechanistically, we find that loss of Nup93 impairs NPC transport, leading to the nuclear accumulation of Yap and downstream inflammation. Collectively, our findings indicate maintenance of endothelial Nup93 as a key determinant of EC health, where aging targets endothelial Nup93 levels to impair NPC function as a novel mechanism for EC senescence and vascular aging.

## INTRODUCTION

Human aging is defined as a chronic state of low-grade inflammation, wherein proinflammatory factors gradually rise, and anti-inflammatory mechanisms decline over time. The pathophysiological features of dysfunctional endothelium closely mirror the consequences of aging endothelial cells (ECs). As such, age-related endothelial dysfunction is now regarded as a prominent precursor to cardiovascular disease (CVD) development^1,2^. While a causal link between endothelial senescence and vascular disease is well-established, the underlying molecular signaling mechanisms triggered by inflammation and aging are unclear.

Endothelial health is largely dictated by the regulation of specific transcription factors that control the transcriptome and identity of ECs. Vessel inflammation and vascular aging are associated with increased Yes-associated protein (Yap) activity, a transcription co-regulator known to activate EC inflammation^3–7^. In the established vasculature, Yap participates in early inflammatory events by promoting the expression of senescence-associated and pro-inflammatory genes in ECs^3–5^. Hence, nuclear accumulation of Yap is a key event for signal activation and downstream inflammation^8^. While nuclear enrichment and downstream Yap activity have become recognized as a key effector of vascular inflammation^4,5^, the mechanisms of Yap regulation are still not fully understood.

Transcriptional regulators, such as Yap, rely on nucleocytoplasmic shuttling, where subcellular localization determines signal outcome. At the molecular level, nucleocytoplasmic transport occurs through nuclear pore complexes (NPCs), one of the largest components of the nuclear envelope^9,10^. Approximately 30 constituent proteins of the NPC, or nucleoporins, play distinct roles in both NPC assembly and function. NPCs permit passive diffusion of small macromolecules (<40 kDa), whereas larger macromolecules depend on an active, carrier-based transport system that requires signal recognition by transport receptors known as karyopherins (*e.g.* exportins, importins)^9,11^. Previous groups have established nucleoporins as extremely long-lived proteins with limited turnover in post-mitotic cells^13,14^. As such, growing evidence points to the deterioration of nucleoporins as a major consequence of aging in long-lived cells^15–18^. Nucleoporin93 (Nup93) is critical for the NPC structure and one of the most abundant nucleoporins in mammalian cells known to deteriorate with age^13,19,20^. As arguably one of the largest organ systems in the human body, the adult endothelium is constantly subjected to various stressors and particularly vulnerable to aging. Categorized as an inner scaffold nucleoporin, Nup93 plays a vital role in the recruitment of other nucleoporins required for NPC formation and their cargo transport function^19^. It is therefore possible that endothelial NPCs, namely Nup93, accumulate damage over time, thereby impairing NPC transport properties that lead to premature vascular aging and accelerated onset of CVD.

The present study sought to determine the relationship between long-term inflammation and endothelial Nup93 protein expression. We find that chronic inflammation decreases endothelial Nup93 levels to permit Yap signaling for EC senescence. These consequential effects reflect abnormal NPC transport properties, as both long-term inflammation and targeted loss of endothelial Nup93 leads to aberrant nuclear accumulation of Yap. Experiments revealed that restoring expression of Nup93 in senescent ECs reverses aging and promotes endothelial reprogramming toward a healthy EC phenotype. We demonstrate endothelial Nup93 as an indispensable player in vascular health and suggest strategies to maintain baseline levels of Nup93 may prevent age-related vascular disease.

## MATERIALS AND METHODS

### Animal Procedures

Aging studies reflect young (1-month) and aged (17-month) wild-type (WT) C57BL/6 mice maintained on a standard laboratory diet (Teklad 7912, Envigo). Atherosclerosis studies were performed in ApoE^‒/‒^ knockout mice (JAX, C57BL/6) as previously described^22–24^. At ∼10 weeks of age, ApoE^‒/‒^ mice were fed *ad libitum* with a high-fat (40% kcal), high-cholesterol (1.25%) Western diet (Research Diets, D12108) or a standard laboratory diet for 12 weeks. Littermates were randomly assigned to the respective experimental groups, and samples were numerically coded to facilitate blinded analyses. Both male and female mice were used for the present studies. At harvest, mice were euthanized via prolonged isoflurane inhalation followed by exsanguination under anesthesia. All animal experiments were approved by the Institutional Animal Care and Use Committee (IACUC) at the University of Illinois at Chicago.

### Isolation of Mouse Lung Endothelial Cells (MLECs)

MLECs were isolated as previously described^25–27^. In brief, mice were anesthetized as described above and perfused with PBS (pH 7.4) via the left ventricle prior to removal of the lung. Perfused lungs were finely minced and digested in 300U/mL Type I Collagenase/PBS solution (37°C, 1 hour), Digested tissues were passed through a 70µm cell strainer, centrifuged and then resuspended in sterile 0.1% BSA/PBS solution. Magnetic Dynabeads (Invitrogen, 11035) were pre-conjugated with an anti-mouse PECAM1 (BD553370) antibody and added to the cell solution for EC enrichment (15min, room temp.). Bound ECs were washed in PBS and harvested for protein analysis using a RIPA buffer-based lysis solution. Protein concentrations were determined using a standard Bradford Colorimetric Assay kit (Bio-Rad, 5000116).

### Immunostaining of Mouse Tissues

Mice were anesthetized as described above and perfused with PBS (pH 7.4) prior to removal of the target organs (*e.g.* heart, aortic arch, brachiocephalic artery). Samples were fixed with 4% PFA/PBS solution (4°C, overnight) followed by paraffin embedding or dehydration in 30% Sucrose/PBS solution for OCT embedding. Tissues were sectioned (5µm), permeabilized with 0.2% Triton-X 100 /PBS (10 minutes, room temperature), and blocked with 5% BSA/PBS (1 hour, room temp). Adjacent tissue sections were incubated with the following primary antibodies (4°C, overnight): anti-Nup93 (1:500, Lusk Lab, Yale University); anti-γH2AX (1:200, Sigma-Aldrich, 05-636); anti-Mac2 (1:100, Cedarlane, CL8942AP); anti-Yap (1:200, Santa Cruz, sc101199); anti-VWF (1:200, Abcam, ab11713). Tissue sections were subsequently incubated with corresponding AlexaFluor secondary antibodies (2 hours, room temp), and nuclei were stained with DAPI. Images were obtained using an inverted fluorescent microscope (Leica DMi8) and quantified using the ImageJ software.

### Cell Culture

Human retinal endothelial cells (HRECs) were purchased from (Cell Systems, ACBRI-181-V) and cultured in endothelial cell growth medium (Lonza, CC-3162) supplemented with 5% FBS (Corning, 35-011-CV), 1% L-glutamine, and 1% penicillin-streptomycin (37°C in 5% CO_2_). Confluent HRECs were subcultured every 3 days, and cells before passage 10 were used for all experiments. Human monocytes (THP-1) were purchased (ATCC, TIB-202) and cultured in RPMI-1640 medium (Corning, 15-040-CV) supplemented with 10% FBS, 1% L-glutamine, and 1% penicillin-streptomycin. Human embryonic kidney 293T (HEK293T) cells used for lentivirus packaging were generously provided by Dr. Jonathan Coloff (UIC) and cultured in high glucose DMEM (Lonza, BE12-709F) supplemented with 10% FBS, 1% L-glutamine, and 1% penicillin-streptomycin.

### Lentiviral Transduction

Glycerol stocks of pLKO.1-puro lentiviral plasmid vectors containing shRNA targeting human Nup93 (NM_014669) and an empty vector insert (shEmpty) were purchased from Sigma-Aldrich. Lentivirus expressing shNup93 or shEmpty were generated by PEI-mediated co-transfection of pLKO.1-puro, psPAX2, and pMD2.G in HEK93T cells (72 hours). Lentiviral-containing media was collected, sterile-filtered through a 0.45µm PES membrane, and stored at −80°C. Human Nup93 (NM_014669) and an empty insert (Lenti–Empty) were subcloned in-house into a pCCL-PGK-MCS lentiviral plasmid vector (gift from Dr. Jan Kitajewski [UIC]). The human Nup93 lentiviral construct was further designed to contain a FLAG tag at the C-terminus of Nup93. Lentivirus expressing Nup93 (Lenti–Nup93-FLAG) or Lenti–Empty were generated using a similar method as above. The Rev-Glucocorticoid Receptor-GFP (RGG) expression vector (gift from Dr. John Hanover^28^ [NIH]) was subcloned into a pLenti-puro lentiviral plasmid vector. Lentiviral particles for the RGG construct were generated using a similar method as described above. Primary sub-confluent HRECs were transduced with the respective lentivirus (1:10 dilution in EC growth medium) for 20 hours (37°C, 5% CO_2_) followed by incubation of lentivirus-infected HRECs in virus-free EC growth medium for 48 hours. Confluent HRECs infected by lentiviral vectors carrying puromycin-resistance were treated with 2.5µg/mL puromycin (Gibco, A11138-03) for 24 hours to eliminate non-transduced cells.

### β-Galactosidase Senescence Staining

Senescence-associated β-Galactosidase (SA-βGal) was measured according to the manufacturer’s instructions (Cell Signaling, 9860S). In brief, confluent HRECs were fixed with the provided 1X Fixative solution, washed in PBS, and incubated in β-Gal Staining solution (pH 6) overnight at 37°C to develop the cyan color in senescent cells. Images were visualized using an inverted fluorescent microscope (Leica DMi8) and quantified using the ImageJ software.

### Cell Cytotoxicity

*In vitro* cytotoxicity was determined by measuring lactate dehydrogenase (LDH) release upon cell lysis according to the manufacturer’s instructions (Promega, G1780). In brief, culture media from confluent HRECs were transferred to sterile 96-well plates and treated with CytoTox96 (30 minutes, room temp) followed by equal volume of Stop Solution (1M acetic acid). LDH release from HRECs was assessed by measuring absorbance at 490nm and comparing values to a Maximum LDH Release sample. Cell cytotoxicity values are presented relative to control conditions.

### Western Blotting

Cells were harvested for protein analysis using a RIPA-based lysis solution. Protein concentrations were determined using a standard Bradford Colorimetric Assay kit (Bio-Rad, 5000116). Proteins were separated by SDS-PAGE and transferred to 0.45µm nitrocellulose membranes. Membranes were blocked in 1% Casein/PBS (1 hour, room temp) and incubated with the following primary antibodies (4°C, overnight): anti-Nup93 (1:1000, Sigma-Aldrich, HPA017937); anti-Nup107 (1:200, Invitrogen, MA1-10031); anti-LaminB1 (1:2000, Proteintech, 12987-1-AP); anti-γH2AX (1:1000, Sigma-Aldrich, 05-636); anti-H2AX (1:1000, Cell Signaling, 2595S); anti-GAPDH (1:1000, Cell Signaling, 2118S); anti-ICAM-1 (1:1000, Abcam, ab109361); anti-VCAM-1 (1:5000, Abcam, ab134047); anti-β-Actin (1:5000, Sigma-Aldrich, A5441); anti-Yap (1:1000, Santa Cruz, sc101199); anti-RelA (1:1000, Proteintech, 10745-1-AP); anti-Stat3 (1:1000, Cell Signaling, 12640S); anti-FoxO1 (1:1000, Cell Signaling, 2880S); anti-LaminA/C (1:1000, Cell Signaling, 4777S); anti-α/β-Tubulin (1:1000, Cell Signaling, 2148S). Corresponding DyLight secondary antibodies (Rockland) were introduced (1 hour, room temp) and visualized using the LI-COR Odyssey Imaging System. Using ImageJ software, protein levels were normalized to housekeeping protein. For fractionation experiments, nuclear fractions were normalized to LaminA/C whereas cytoplasmic fractions were normalized to α/β-Tubulin. Normalized protein values are presented relative to control conditions.

### Subcellular Fractionation Assay

Nuclear and cytoplasmic protein extracts were isolated from confluent HRECs using a Subcellular Fractionation (SF) buffer (250mM Sucrose, 20mM HEPES pH 7.4, 10mM KCl, 2mM MgCl_2_, 1mM EDTA, 1mM EGTA) supplemented with protease inhibitor cocktail. In brief, HRECs were harvested with SF buffer, agitated on a rotator (4°C, 30 minutes), and centrifuged to separate cytoplasmic (supernatant) and nuclear (pellet) fractions. Cytoplasmic fractions were transferred into a separate microcentrifuge tube, while nuclear fractions were lysed using RIPA buffer-based lysis solution supplemented with protease inhibitor cocktail. Both nuclear and cytoplasmic extracts were analyzed via immunoblotting, as detailed above.

### Immunofluorescence Staining

Confluent HRECs were fixed in 4% PFA/PBS (10 minutes, room temp) and permeabilized in 0.2% Triton-X 100/PBS (10 minutes, room temp). Cells were subsequently blocked in 5% BSA/PBS (1 hour, room temp) and incubated with the following primary antibodies (4°C, overnight): anti-Nup93 (1:750, Lusk Lab, Yale University); anti-LaminB1 (1:250, Proteintech, 12987-1-AP); anti-γH2AX (1:250, Sigma-Aldrich, 05-636); anti-Yap (1:250, Santa Cruz, sc101199); anti-GFP (1:250, Santa Cruz, sc9996). Cells were then incubated with corresponding AlexaFluor secondary antibodies (2 hours, room temp) and nuclei were stained with DAPI. Images were visualized using an inverted fluorescent microscope (Leica DMi8) and quantified using ImageJ software. To quantify nuclear and cytoplasmic localization, signal intensities in cell nuclei were calculated relative to DAPI staining. Cytoplasmic localization was then quantified by subtracting nuclear signal from the whole image intensity. Nuclear signals were divided by cytoplasmic intensity in each condition for ratiometric data presentation.

### Monocyte Adhesion Assay

THP-1 monocytes (2×10^5^ cells/mL) were labeled with 5µM cell-permeant Calcein-AM in DMSO (1:200, BioLegend, 425201) for 20 minutes at 37°C, centrifuged and then resuspended in endothelial cell growth medium. Labeled THP-1 cells were incubated with confluent HRECs for 15 minutes at 37°C, and non-adherent cells were removed by gentle PBS washing. Adherent THP-1 cells and HRECs were fixed with 4% PFA in PBS solution for 10 minutes at room temperature, washed in PBS, and visualized using an inverted fluorescent microscope (Leica DMi8). EC-monocyte attachment was quantified by counting the number of adhered fluorescent THP-1 using ImageJ software.

### RNA Extraction and RT-qPCR

Confluent HRECs were harvested for total RNA using the RNeasy Mini Kit (Qiagen, 74104). RNA purity and concentration were assessed with a NanoDrop One instrument (Invitrogen). Total RNA (0.5µg) was converted into cDNA using the High-Capacity cDNA Reverse Transcription Kit (Applied Biosystems, 4368814). RT-qPCR was performed using the Fast SYBR Green Master Mix (Applied Biosystems, 4385612) with the ViiA 7 Real-Time PCR System (Applied Biosystems). Relative mRNA levels were normalized to GAPDH, and normalized levels were further expressed relative to control. The primer sequences used in this study are listed in Supplemental Table 1.

### Bulk RNA Sequencing

Bulk RNA sequencing was performed in isolated RNA samples (1.5µg) and analyzed with a reference genome (NovoGene Co Inc). Differentially expressed genes were identified (Padj<0.01 and fold change cut-off>2) for subsequent pathway analyses using Ingenuity Pathway Analysis (IPA) software and the Reactome Pathway database. IPA was used for enrichment analyses of pathways and upstream transcriptional regulators, whereas Reactome was used solely for enrichment analyses of pathways. The analyzed RNA sequencing data, including upregulated and downregulated DEGs, are available in Supplemental Table 2.

### Flow Cytometry

Confluent HRECs were detached using 0.05% Trypsin (Corning, MT25052CI) and collected in growth medium. Cells were centrifuged and resuspended in sterile FACS buffer (3% FBS, 2mM EDTA, 0.1% NaN_3_). Cell suspensions were incubated with the following antibodies for (20min, on ice): anti-ICAM1-PB (1:25, BioLegend, 322715); anti-VCAM1-PE (1:25, BioLegend, 305805); anti-E-Selectin-PE (1:25, BioLegend, 322605); AnnexinV-FITC (1:25, BioLegend, 640905); Propidium Iodide (1:50, BioLegend, 421301). Cells were then washed and resuspended in ice-cold FACS buffer. Cell suspensions were additionally incubated with 7-AAD Solution (1:100, BioLegend, 420404) for 5 minutes to detect dead cells and passed through cell strainer tubes (Falcon, 352235). Flow cytometric analysis was performed using the CytoFLEX S Flow Cytometer (Beckman Coulter) and quantified with the FlowJo v10 software. Surface levels of ICAM1, VCAM1, and E-Selectin were calculated from mean fluorescence intensities and further normalized relative to control. Cell proliferation was measured using the Click-iT EdU Flow Cytometry Assay Kit (Invitrogen, C10425) following manufacturer’s instructions.

### Chemicals and Reagents

Tumor necrosis factor alpha (TNFα) was purchased (Sigma, H8916) and dissolved in sterile Milli-Q water to a stock concentration of 10µg/mL for *in vitro* experiments. Verteporfin (VP) was purchased (ApexBio, A8327) and dissolved in sterile DMSO to a stock concentration of 10mM for *in vitro* experiments. Dexamethasone (DEX) (10mM, in DMSO) was purchased (Gibco, A13449) for *in vitro* experiments.

### Statistical Analysis

All data are presented as mean ± SEM and analyzed using GraphPad Prism v8.0 software. Statistical analyses reflect an unpaired two-tailed Student t-test for two-group comparisons of normally distributed data with equal variance. Multi-group comparisons with equal variances were performed using a 1-way ANOVA followed by Dunnett’s posthoc test or 2-way ANOVA followed by Sidak test. A p-value<0.05 was considered statistically significant.

## RESULTS

### Endothelial Nup93 expression declines with age and chronic inflammation

To determine how aging impacts endothelial Nup93 protein expression, highly vascularized tissues (*i.e.* heart, lung) were isolated from both young (1-month) and aged (17-month) mice. Cross-sectional analysis of the coronary vasculature was limited to the endothelial layer, as determined by VWF expression. Endothelial expression of γH2AX (or H2AX phosphorylated at serine 139), a marker for DNA damage and aging, was significantly upregulated in aged mice when compared to younger cohorts (**Figure 1A&1B**), as expected. Endothelial expression of Nup93 was, however, substantially reduced in the coronary vasculature of aged mice (**Figure 1A&1C**). Primary mouse lung ECs (MLECs) were also isolated from young and aged mouse lung tissue for subsequent immunoblot analyses. Indicative of aging, MLECs isolated from aged mice displayed significantly higher levels of γH2AX and reduced expression of LaminB1, both well-established molecular markers of cellular senescence (**Figure 1D-F**). More importantly, aged mice showed a striking reduction in Nup93 protein levels to corroborate the immunofluorescence data (**Figure 1G**), whereas other scaffold nucleoporins, such as Nup107, remained largely unaffected (**Figure 1H**).

**Figure 1.**
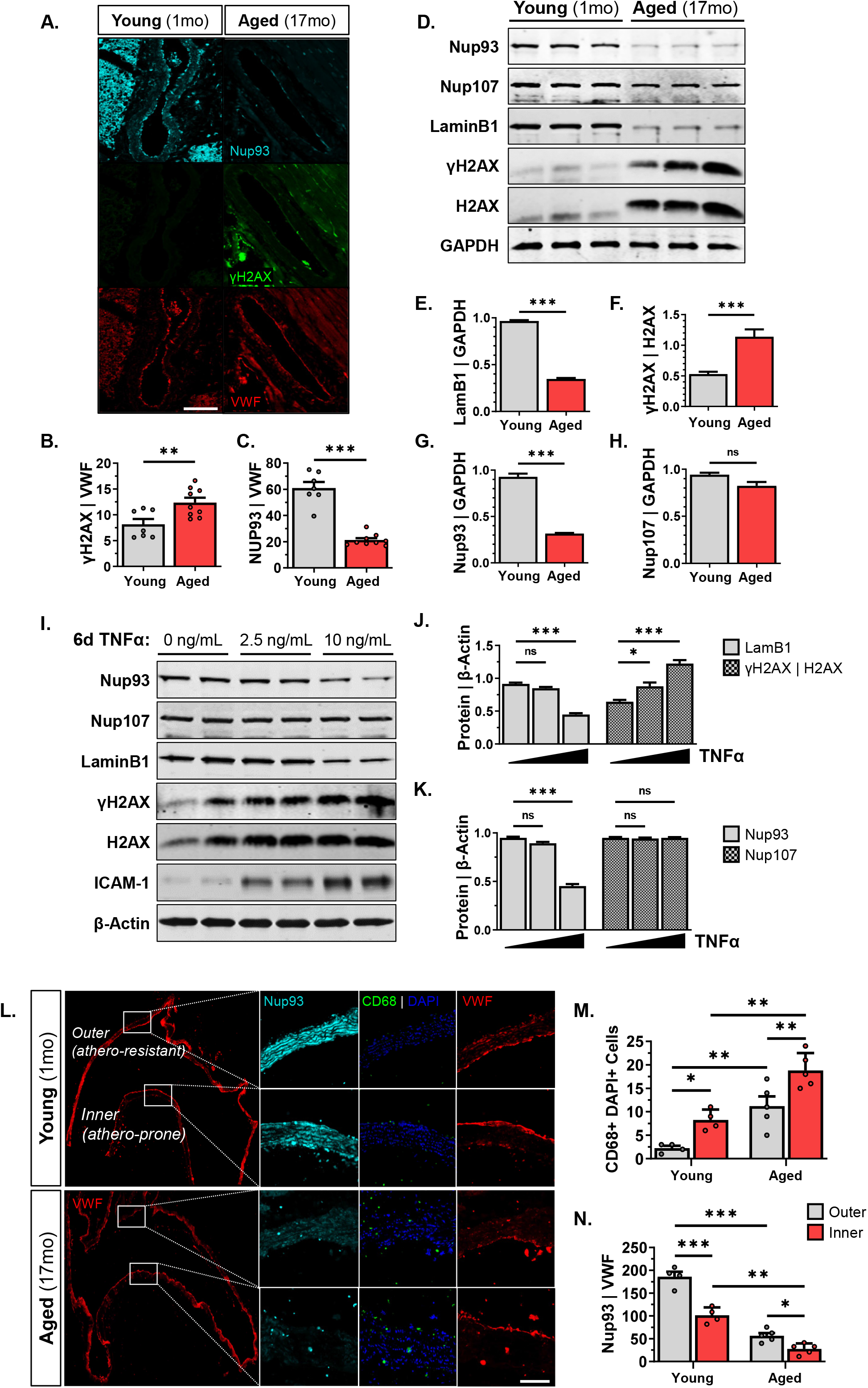
Endothelial Nup93 expression declines with vascular aging and chronic inflammation. **(A)** Immunofluorescence staining of young (1-month) and aged (17-month) mice indicates an age-associated increase in γH2AX expression and corresponding decrease in Nup93 in the coronary vasculature. (n=7 young, n=9 aged). Scale bars=100µm. Quantified in **(B)** and **(C)**. **(D)** Immunoblotting of primary MLECs isolated from aged mice express established senescence markers and a similar decrease in Nup93 protein levels when compared to young mice. (n=3-4 mice per lane). Nup107 remains unaffected. Quantified in **(E-H)**. **(I)** Chronic TNFα treatment (10ng/mL, 6days) in primary HRECs leads to a similar acquisition of senescence markers and decreased Nup93 protein expression, whereas Nup107 remains unchanged. Quantified in **(J)** and **(K)**. **(L)** Representative image of the cross-sectional analysis on the aortic arch showing the inner (athero-resistant) and outer (athero-prone) curvature. Immunofluorescence staining in the aortic arch indicates an age associated increase in CD68+ positive cells and decreased endothelial Nup93 expression. (n=4 young, n=5 aged). Scale bars=100µm. Quantified in **(M)** and **(N)**. *\*\*\* p<0.001, ** p<0.01, * p<0.05*

To model endothelial aging *in vitro*, primary human retinal ECs (HRECs) were exposed to the inflammatory cytokine tumor necrosis factor alpha (TNFα) for 6 consecutive days^29^. Similar to previous reports, chronic exposure to TNFα leads to increased expression of adhesion molecules (*i.e.* ICAM1), a feature of EC inflammation and senescence (**Figure 1I**). Moreover, *in vitro* modeling of endothelial aging revealed increased γH2AX levels and decreased LaminB1, confirming the established link between chronic inflammation and endothelial senescence (**Figure 1I&1J**). Intriguingly, endothelial Nup93 protein levels declined with long-term exposure to TNFα, whereas Nup107 protein expression was not affected, similar to that observed in isolated primary MLECs (**Figure 1I&1K**). To compare the effects of chronic versus acute inflammation, HRECs were treated with TNFα for 24 hours at concentrations up to 40ng/mL. ICAM1 protein levels were significantly upregulated across the range of TNFα concentrations used (**Figure S1A&S1B**). Protein levels of Nup93 and LaminB1, however, were not significantly altered with acute EC inflammation, even when using higher TNFα concentrations (**Figure S1A-D**). Taken together, these findings indicate that endothelial Nup93 is selectively reduced with aging and chronic low-grade inflammation.

### Vascular inflammation is associated with suboptimal levels of endothelial Nup93

Our initial findings established an inverse correlation between inflammation and endothelial Nup93 expression to suggest Nup93 protein may be targeted in other conditions of vascular inflammation. We therefore took advantage of the hemodynamic influences on the murine aortic arch to assess the *in vivo* relationship between endothelial Nup93 and flow-driven vascular inflammation. It is well-established that laminar and disturbed blood flow patterns activate different signaling cascades in ECs, resulting in an atheroprotective and atheroprone phenotype, respectively^30^. Both flow patterns are known to occur in the aortic arch; the outer curvature exhibits an anti-inflammatory profile whereas the inner curvature exhibits a pro-inflammatory profile^31^ (**Figure 1L**). Immunohistological staining of the aortic arch in WT mice revealed an increase in CD68-positive cells in the inner curvature, a macrophage cell-marker and common readout of vessel inflammation, as found in previous reports^32,33^ (**Figure 1L&1M**). These indices of inflammation were accompanied by a significant reduction in endothelial Nup93, as determined by quantification of Nup93 signal intensity in the VWF-positive layer (**Figure 1L&1N**). We find that Nup93 expression is significantly decreased in the inner curvature, a pro-inflammatory region of the aortic arch, thereby corroborating our *in vitro* data.

We next sought to investigate the impact of Western Diet consumption, a trigger for chronic metabolic inflammation, on endothelial Nup93 expression. To test this, 10-week-old atherosclerotic-prone ApoE^-/-^ mice were challenged with a 12-week Western Diet. Immunohistological staining of isolated brachiocephalic arteries (BCA) indicates a significant increase in atherosclerotic lesion deposition, as shown by the visible plaque formation and enhanced Mac2 macrophage expression when compared to chow-fed mice (**Figure S1E&S1F**). More importantly, we find that endothelial expression of Nup93 is significantly reduced in Western Diet-fed mice (**Figure S1E&S1G**). These results mirror our previous observations of decreased endothelial Nup93 expression in the coronary vasculature of aged mice (**Figure 1A**) and suggest suboptimal expression of Nup93 in ECs may precede cardiovascular disease progression.

### Endothelial Nup93 knockdown sensitizes ECs to inflammation

To identify transcriptional programs that may be dysregulated with endothelial loss of Nup93, we performed bulk RNA sequencing (RNAseq) on shEmpty- and shNup93-transduced HRECs. RNAseq analysis revealed 794 differentially expressed genes (DEGs), of which 459 were upregulated and 335 were downregulated with endothelial Nup93 loss (Padj<0.01; fold change cut-off>2) (**Figure 2A, Supp. Table 2**). DEGs were further analyzed using Ingenuity Pathway Analysis (IPA) software to identify canonical pathways associated with the transcriptomic impact of endothelial Nup93 loss. Intriguingly, pathway analyses of the upregulated DEGs identified ‘Granulocyte Adhesion and Diapedesis’, ‘Atherosclerosis’, and other pro-inflammatory associated signaling pathways (**Figure 2B**). Upregulated DEGs were also analyzed using the Reactome Pathway database which indicated the ‘Senescence-Associated Secretory Phenotype’ as a significantly upregulated pathway in Nup93-deficient ECs (**Figure S2A**). Conversely, the downregulated DEGs yielded several biological processes related to cell cycle regulation and DNA damage response through both IPA and Reactome Pathway databases (**Figure S2B&S2C**).

**Figure 2.**
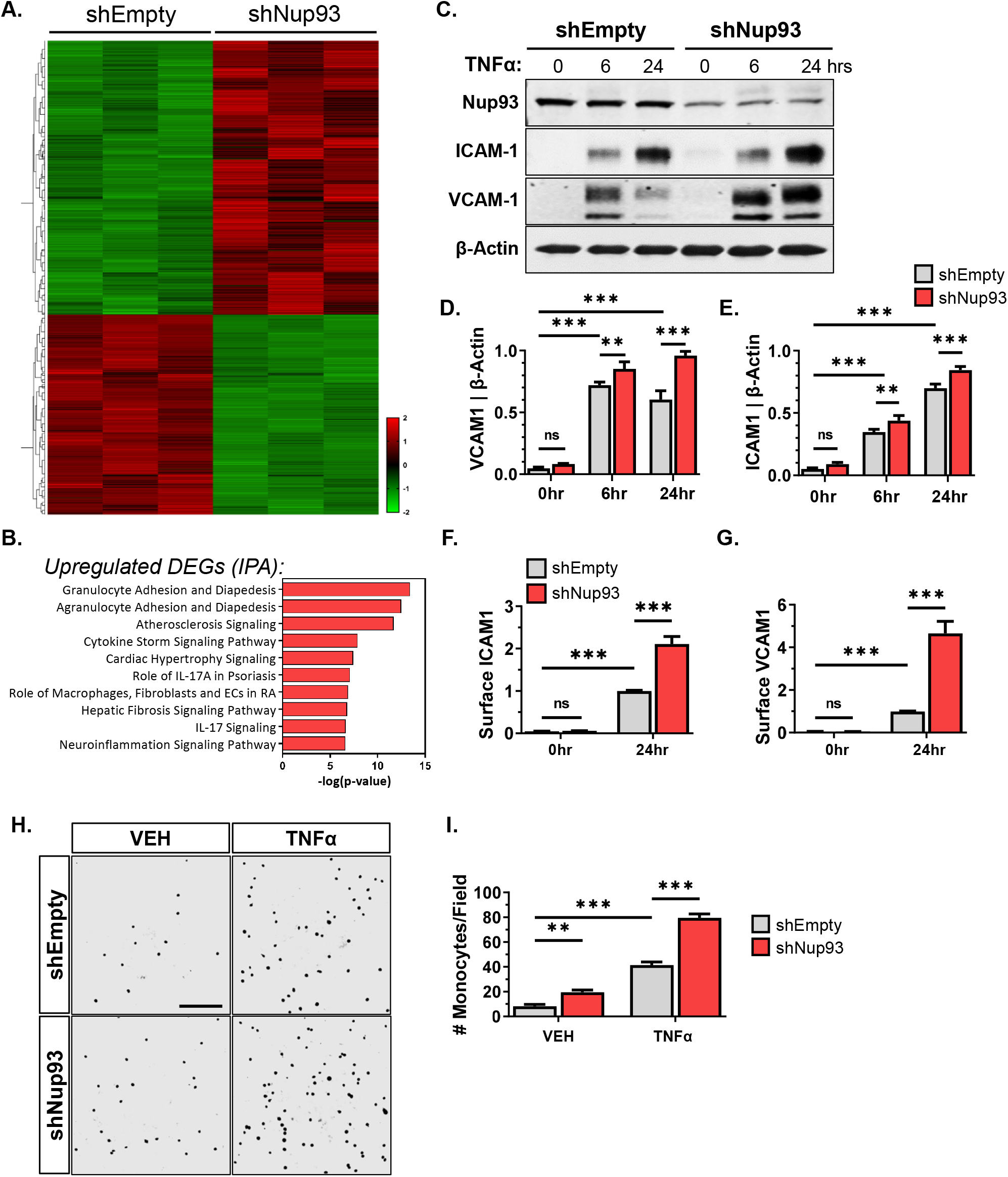
Endothelial loss of Nup93 sensitizes endothelial cells to inflammation. Primary HRECs were transduced with lentiviral constructs (shEmpty & shNup93) for Nup93 knockdown and subsequent bulkRNA sequencing. **(A)** Heatmap of the differentially expressed genes (459 upregulated, 335 downregulated; Padj<0.01; fold change cut-off>2). Horizontal columns: individual genes, Vertical Columns: individual HREC sample. **(B)** Ingenuity Pathway Analysis (IPA) of the upregulated DEGs identifies several inflammation-associated pathways. **(C)** Loss of Nup93 in HRECs together with TNFα [2.5ng/mL] treatment leads to a heightened and sustained expression of both ICAM1 and VCAM1. Quantified in **(D)** and **(E)**. FACS-based analyses of HRECs 24hrs post-TNFα treatment show an increase in surface levels of both **(F)** ICAM1 and **(G)** VCAM1 in Nup93 knockdown ECs. **(H)** Endothelial-monocyte adhesion assays indicate increased monocyte adhesion in TNFα-treated [2.5ng/mL, 24hrs] ECs that is further heightened with loss of Nup93. Quantified in **(I)**. Scale bars=200µm. *n=3, *** p<0.001, ** p<0.01*

Endothelial senescence is associated with a decline in EC health and increased inflammation. To test the consequence of Nup93 loss on EC inflammation, we treated control and Nup93-depleted HRECs with TNFα at various time points (0, 6, and 24 hours; 2.5ng/mL). The loss of Nup93 together with TNFα treatment leads to increased protein expression of adhesion molecules VCAM1 and ICAM1, both of which are expressed in inflamed endothelium (**Figure 2C-E**). The observed increase in inflammatory adhesion molecules occurs within 6 hours of TNFα treatment and is sustained after 24 hours of exposure. Furthermore, analysis using flow cytometry methods corroborates an increase in surface expression of both ICAM1 and VCAM1 (**Figure 2F&2G**). Elevated surface expression of adhesion molecules promotes the attachment of leukocytes to ECs, thereby driving vascular inflammation and disease pathogenesis. We therefore performed endothelial-monocyte adhesion assays using fluorescently-labeled THP-1 monocytes to determine the functional consequence of endothelial Nup93 depletion. Endothelial treatment with TNFα significantly increased the number of adherent monocytes in shEmpty conditions, as expected (**Figure 2H&2I**). Suboptimal levels of endothelial Nup93 together with TNFα treatment led to even greater monocyte attachment, in line with the observed increase in adhesion molecules. Interestingly, reduced endothelial Nup93 expression promoted monocyte adhesion independent of cytokine treatment, as seen in vehicle conditions (**Figure 2H&2I)**. This baseline increase in monocyte attachment is likely attributed to elevated surface levels of E-selectin (**Figure S2D&S2E**) in Nup93 deficient ECs, another well-known adhesion molecule induced by inflammation selectively in ECs. Taken together, these findings demonstrate that loss of Nup93 promotes EC inflammation.

### Endothelial Nup93 depletion induces features of EC senescence

To determine whether decreased Nup93 results in endothelial senescence, shRNA-based methods were used to selectively knockdown Nup93 expression in primary HRECs. Western Blot analyses validated our Nup93-targeted approach, as other structural nucleoporins, such as Nup107, remain unaffected (**Figure 3A-C**). We subsequently examined the expression of well-accepted cell senescence markers using both immunoblotting and immunofluorescence techniques. Endothelial loss of Nup93 resulted in increased levels of γH2AX and reduced LaminB1 protein expression (**Figure 3D&3E**, **Figure S3A-C**) in agreement with our earlier *in vivo* findings (**Figure 1**). Furthermore, expression of senescence-associated β-Galactosidase (SA-βGal) was significantly enhanced in Nup93 depleted ECs, corroborating EC senescence (**Figure 3F&3G**). We next examined the impact of Nup93 depletion on EC proliferation by measuring EdU incorporation using FACs-based methods. Nup93 knockdown in HRECs led to a significant decrease in the percentage of S-phase cells and a corresponding increase in G1-phase cells suggesting G1 arrest (**Figure 3H**). Moreover, Nup93 knockdown approach did not have a significant impact on cellular apoptosis, necrosis, or cytotoxicity (**Figure S3D&S3E**). Therefore, loss of endothelial Nup93 induces features of endothelial senescence while maintaining viability.

**Figure 3.**
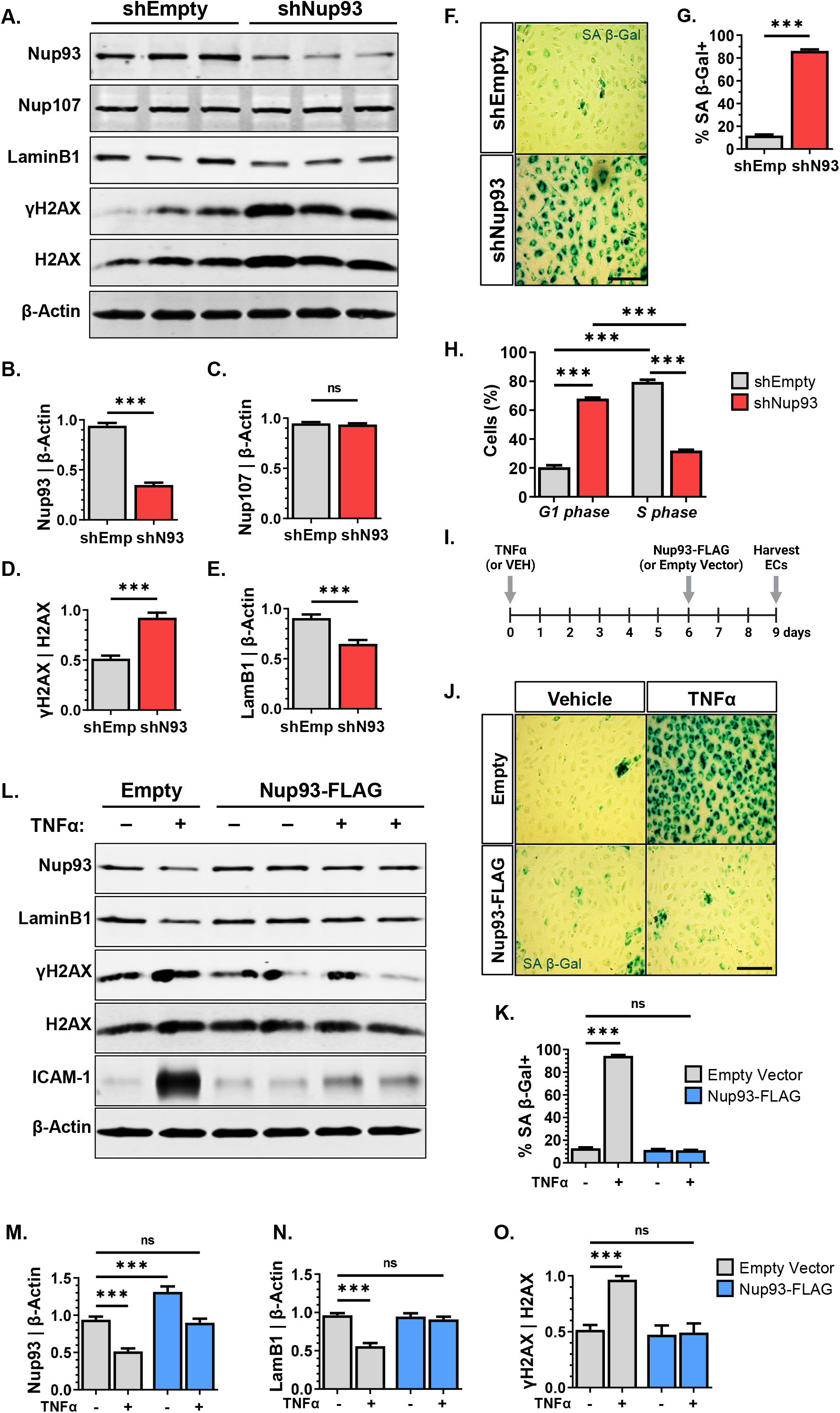
Loss of Nup93 induces features of endothelial cell senescence. **(A)** Immunoblotting analysis indicates a significant decrease in Nup93 protein levels when using shRNA-mediated Nup93 knockdown methods. Nup107 expression remains unaffected. Quantified in **(B)** and **(C)**. Loss of Nup93 leads to increased expression of γH2AX (H2AX phosphorylated at S139) and decreased LaminB1 levels, both well-established readouts for cellular senescence. Quantified in **(D)** and **(E). (F)** Increased SA-βgal staining in Nup93 knockdown ECs. Quantified in **(G)**. **(H)** FACS analyses of EdU-labeled HRECs indicate a significant decrease in S-phase cells and a corresponding increase in G1-phase cells to indicate G1 arrest. **(I)** *Schematic*: HRECs were first exposed to the chronic inflammation model (TNFα [10ng/mL]; 6 days) followed by lentiviral transduction for exogenous expression and restoration of Nup93 protein. **(J)** Chronic inflammation leads to increased SA-βgal expression where exogenous expression of Nup93 in already senescence ECs leads to a significant reduction in SA-βgal, similar to healthy ECs. Quantified in **(K)**. **(L)** Endothelial exposure to chronic inflammation followed by exogenous Nup93 expression restores LaminB1 and γH2AX protein levels comparable to healthy ECs. Quantified in **(M-O)**. Scale bars=200µm. n=3, *\*\*\* p<0.001*

### Restoring Nup93 protein reduces hallmarks of endothelial senescence

Based on the observation that loss of Nup93 triggers features of endothelial senescence, we then hypothesized that Nup93 overexpression would prevent inflammation-induced EC senescence. To test this, lentiviral-based methods were used to initially achieve a moderate overexpression (OE) of Nup93 in primary HRECs, as shown in **Figure S4A&S4D** (*lane1 vs. lanes2&3)*. HRECs (Empty & Nup93-OE) were then exposed to TNFα (10ng/mL) over a course of 6 days. Overexpression of Nup93 delayed the acquisition of cellular senescence markers, as both γH2AX and LaminB1 protein levels were comparable to control (Empty+vehicle, *lane1*) conditions (**Figure S4A-C**). Total Nup93 levels at the end of the 6^th^ day of the inflammation period remained comparable to control conditions (**Figure S4D**). We next sought to determine whether restoring Nup93 would reverse features of aging in already senescent ECs. To test this, HRECs were initially exposed to chronic inflammation (TNFα [10ng/mL]; 6 days) followed by lentiviral transduction for exogenous expression and restoration of Nup93 protein (schematic in **Figure 3I**). Similar to previous results, chronic TNFα treatment in Empty vector controls led to a significant increase in SA-βGal activity indicative of senescence (**Figure 3J&3K**). Exogenous expression of Nup93, however, drastically reduced SA-βGal levels, similar to that of vehicle-treated Empty vector controls (**Figure 3J&3K**). Furthermore, restoring Nup93 protein to baseline levels in already senescent ECs promoted the recovery of LaminB1 expression and decreased γH2AX to levels comparable to that observed in healthy and non-senescent ECs (**Figure 3L-O**). Taken together, these findings indicate that a baseline level of Nup93 is necessary to maintain EC health, whereas age-related chronic inflammation leads to suboptimal Nup93 expression, EC senescence, and dysfunction.

### Endothelial Nup93 depletion enhances Yap nuclear localization and activity

RNAseq analyses of Nup93-deficient ECs identified several pro-inflammatory and senescence-related pathways (**Figure 2B & Figure S2A**). Given the importance of Nup93 in NPC function, we reasoned that the Nup93 loss may lead to the mislocalization of transcription factors (TFs) that provoke aberrant gene expression. We therefore utilized the IPA Upstream Regulator analysis feature to identify upstream mediators that could explain the observed gene expression changes following loss of Nup93. Enrichment analyses of the differentially upregulated genes in Nup93-deficient ECs identified several TFs known to drive endothelial inflammation (**Figure 4A**). Intriguingly, these analyses also identified the transcriptional cofactor Yap and several of its canonical partners (*i.e.* RelA, Stat3, FoxO1)^34–36^ as upstream regulators predicted to drive transcriptional changes associated with Nup93 loss (**Figure 4A**). Previous studies reported a significant correlation between vascular aging and nuclear accumulation of Yap^3^, where Yap activity in the adult vasculature is known to trigger EC inflammation and disease^4,5,7^. Here, we found that chronic exposure to TNFα (10ng/mL, 6 days) significantly increased nuclear accumulation of Yap, as assessed through immunofluorescent staining (**Figure S4E&S4F**). Moreover, endothelial exposure to long-term inflammation increases downstream Yap target gene expression (*i.e.* CYR61, CTGF) in support of previous reports of Yap hyperactivation associated with inflammation (**Figure S4G&S4H**).

**Figure 4.**
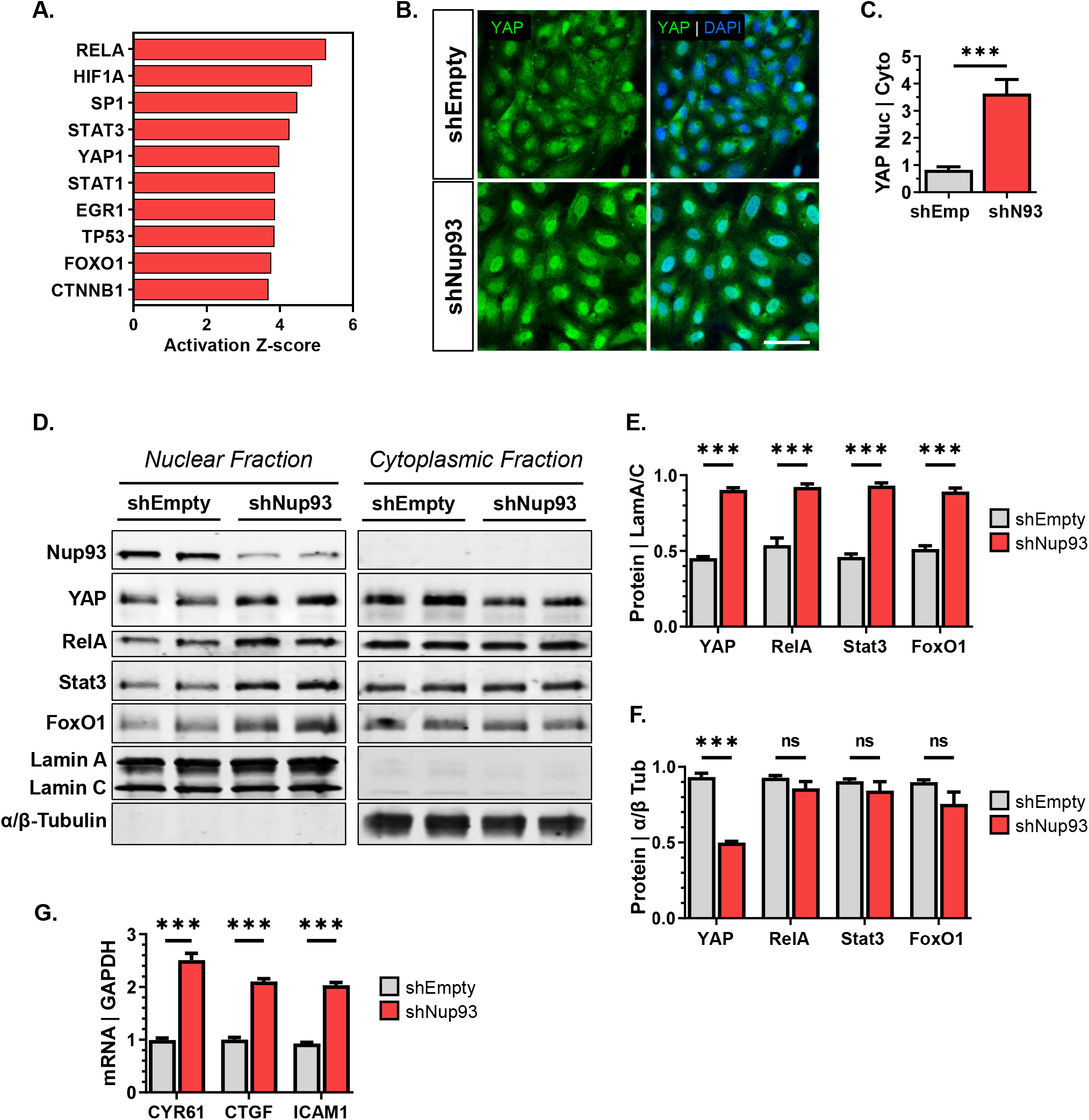
Endothelial loss of Nup93 increases nuclear Yap levels for downstream signaling. **(A)** The differentially upregulated genes associated with Nup93 knockdown were subject to pathways analyses (IPA) to identify several upstream transcriptional regulators, including Yap. **(B)** Immunofluorescence staining of Yap in shEmpty and shNup93-transduced HRECs. Quantified in **(C)**. **(D)** Immunoblotting of nuclear and cytoplasmic fractions in shNup93-transduced HRECs and shEmpty vector controls. Quantification of select proteins in the nucleus **(E)** and **(F)** cytoplasm. **(G)** RT-qPCR analysis indicates a significant increase in Yap target gene expression in Nup93 depleted ECs. Scale bars=50µm. n=3, *\*\*\* p<0.001*

We next sought to determine the putative regulatory role of Nup93 in endothelial Yap signaling. We observed increased nuclear Yap levels with endothelial loss of Nup93 when using immunofluorescence methods (**Figure 4B&4C**). Moreover, subcellular fractionation indicated a significant increase in nuclear Yap levels along with its canonical partners (*e.g.* RelA, Stat3, FoxO1) (**Figure 4D&4E**). Cytoplasmic levels of RelA, Stat3, and FoxO1 remained largely unaffected whereas Yap cytoplasmic levels were significantly reduced upon endothelial Nup93 knockdown (**Figure 4D&4F**). Moreover, RT-qPCR indicated a significant increase in Yap target gene expression (**Figure 4G**). These findings are similar to results obtained when using the chronic inflammation model (**Figure S4E-S4H**) in support of a regulatory role of Nup93 in endothelial Yap signaling.

### Yap inhibition attenuates EC senescence and inflammation induced by Nup93 deficiency

To definitively link enhanced Yap activity as the primary mechanism of endothelial dysfunction in Nup93-deficient human ECs, we introduced verteporfin (VP), a potent pharmacological inhibitor of Yap. In validating the pharmacological effects of VP, we found that VP treatment (0.5μM, 24hrs) in shEmpty vector transduced control cells significantly decreased Yap nuclear-to-cytoplasmic levels, as shown by immunofluorescent staining (**Figure 5A&5B**). These effects of VP were further corroborated when using subcellular fractionation methods (**Figure S5A-C**). Our previous observations indicated increased baseline levels of nuclear Yap in Nup93-deficient HRECs. Pharmacological treatment with VP resulted in an even more pronounced effect in Nup93-depleted ECs and the nuclear-to-cytoplasmic ratio of Yap post-VP treatment was restored to levels similar to that observed in shEmpty VP-treated controls (**Figure 5A&5B**). Furthermore, we found that VP treatment in Nup93-deficient ECs mitigates Yap transcriptional activity to levels comparable to control conditions, as indicated by the significant reduction in Yap target gene expression, including inflammatory genes (*i.e.* ICAM1) (**Figure S5D-F**). These results validate the inhibitory effect of VP on Yap signaling and confirm Yap hyperactivity as the major mechanism of inflammatory gene expression in Nup93-deficient ECs.

**Figure 5.**
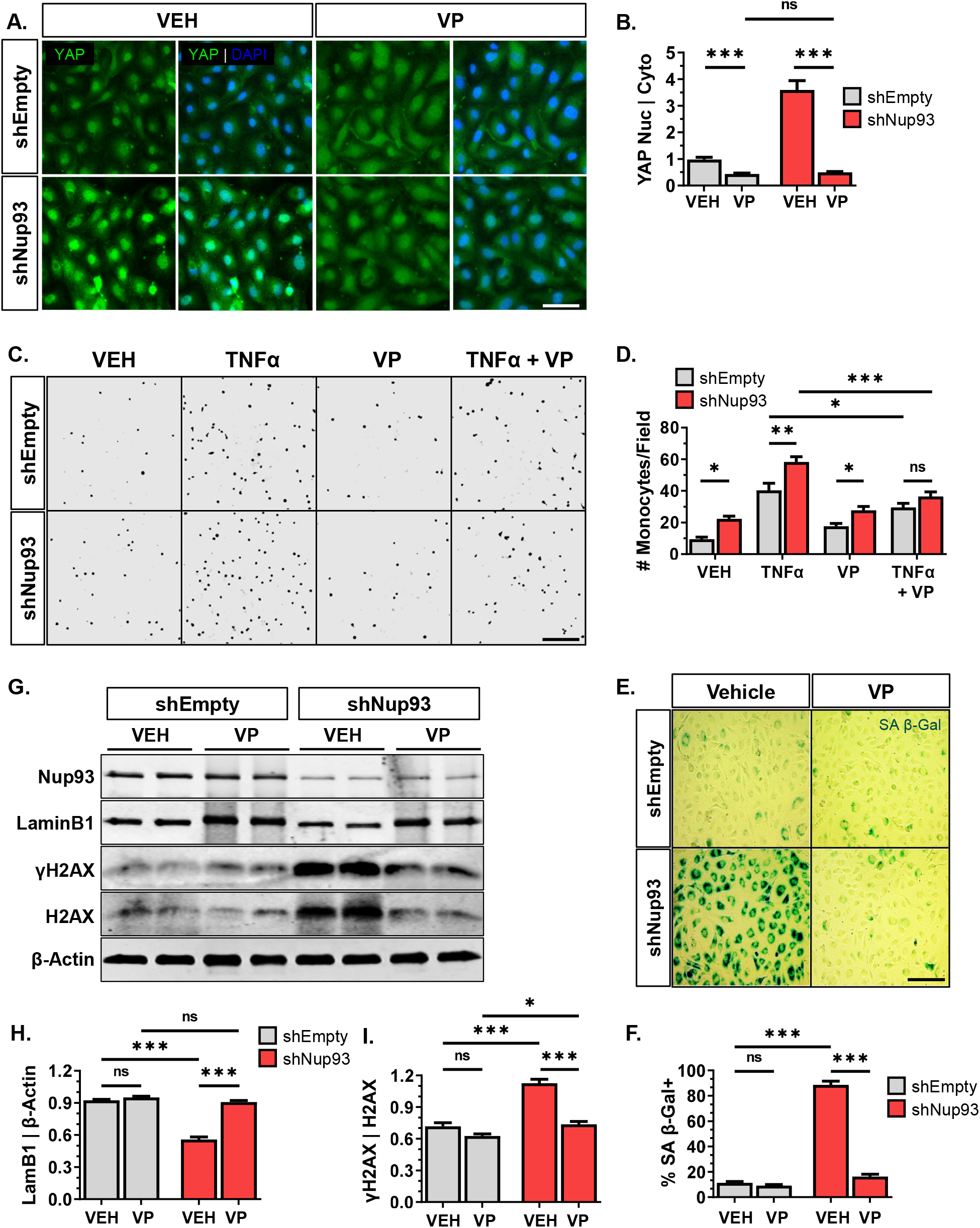
Pharmacological inhibition of Yap attenuates endothelial senescence and inflammation induced by Nup93 deficiency. **(A)** Immunofluorescence staining of shEmpty- and shNup93-transduced HRECs treated with verteporfin (VP, [0.5µM]) or vehicle for 24 hours. Scale bar=50µm. Quantified in **(B)**. **(C)** Representative images of adherent THP-1 monocytes across indicated conditions. HRECs (shEmpty & shNup93) were treated with verteporfin [0.5µM], TNFα [2.5ng/mL], or both verteporfin and TNFα prior to incubation with fluorescently labeled monocytes (pseudo colored to grayscale). Scale bar=200µm. Quantified in **(D)**. **(E)** Representative SA β-Gal staining of shEmpty- and shNup93-transduced HRECs treated with verteporfin ([0.5µM], 24hrs) shows a significant reduction in SA β-Gal signal, similar to healthy ECs. Scale bar=200µm. Quantified in **(F)**. **(G)** Verteportin treatment in shNup93-transduced cells restores LaminB1 and γH2AX protein levels comparable to healthy ECs. Quantified in **(H)** and **(I)**. n=3, *\*\*\* p<0.001, ** p<0.01, * p<0.05*

To examine the functional effects of VP treatment in Nup93-deficient ECs, endothelial-monocyte adhesion assays were again performed. Consistent with our previous observations, loss of endothelial Nup93 significantly increased the number of adherent THP-1 monocytes in both vehicle and TNFα-treated ECs (**Figure 5C&5D**). Treatment with VP, however, significantly reduced monocyte attachment in TNFα-exposed ECs, thereby suggesting that Yap inhibition is sufficient to attenuate the augmented inflammation seen with the combination of both TNFα exposure and Nup93 loss (**Figure 5C&5D**). However, the baseline increase in monocyte attachment as a result of Nup93 loss was unaffected by VP treatment to insinuate that Nup93-depletion promotes the expression of adhesion molecules independent of Yap activity.

Given the observed benefits of VP treatment in Nup93-deficient ECs, we next evaluated whether VP treatment could attenuate features of EC senescence in Nup93-depleted ECs. To test this, primary HRECs were infected with shEmpty or shNup93 lentiviral constructs. Once fully confluent, transduced ECs were treated with VP (0.5μM) or vehicle for 24hrs. As shown in previous results, loss of Nup93 significantly increased the percentage of SA-βGal positive ECs in vehicle-treated conditions (**Figure 5E&5F**). Inhibition of Yap, however, resulted in a drastic attenuation of SA-βGal activity to levels comparable to that observed in healthy ECs (**Figure 5E&5F**). VP treatment in Nup93-deficient ECs also restored LaminB1 protein expression and decreased γH2AX levels, thereby indicating a reprogramming toward a healthy EC profile (**Figure 5G-I**). Taken together, these findings demonstrate that EC senescence is largely reliant on Yap activation. Moreover, inhibiting Yap signaling reverses features of EC senescence to parallel observations when Nup93 protein levels are restored.

### NPC stability is adversely impacted by Nup93 deficiency in ECs

Nup93 is one of the most critical structural nucleoporins involved in the regulation of NPC function and transport properties^19,37^. Small molecules (<40kDa) passively shuttle through the NPC, whereas larger cargos are actively transported with high selectivity^10^. Our previous results indicate Yap hyperactivation as a primary mechanism of EC senescence, where loss of Nup93 may consequently promote nuclear Yap accumulation. Using our previously introduced Nup93 restoration model (**Figure 3I)**, lentiviral-mediated introduction of Nup93 did not affect baseline cellular Yap distribution in vehicle-treated ECs (**Figure 6A&6B**). Exogenous Nup93 expression in TNFα-treated ECs, however, greatly reduced the nuclear Yap signal comparable to that of healthy and non-senescent ECs (**Figure 6A&6B**). Based on these observations, we reasoned that endothelial loss of Nup93 may destabilize NPCs and consequently enhance baseline nuclear permeability. To investigate the impact of Nup93 loss on nuclear transport, HRECs were transduced with a dexamethasone (DEX) responsive GFP-tagged glucocorticoid receptor (GR)^28^. This 64kDa engineered construct contains the full-length Rev sequence and the hormone-responsive element from the GR to provide a steroid-specific nuclear localization sequence (‘RGG’, **Figure 6C**). Similar to the molecular weight of Yap, this artificial reporter system is therefore hormone-inducible and can precisely control nuclear protein import of GFP. In line with previous groups using this established reporter system^38–40^, the RGG construct localizes to the cytoplasm in the absence of steroid, where GFP signal is predominantly cytoplasmic in shEmpty control HRECs (**Figure 6D&6E**). Validating the utility of the RGG construct, we found that DEX treatment (1μM, 30min) in shEmpty transduced ECs induced the nuclear import of the GFP signal, as expected (**Figure 6D&6E**). shRNA-mediated loss of Nup93, however, significantly increased nuclear GFP intensity in the absence of DEX. Treatment of Nup93-deficient HRECs with DEX led to a further increase in nuclear GFP signal, similar to DEX-treated shEmpty control HRECs (**Figure 6D&6E**).

**Figure 6.**
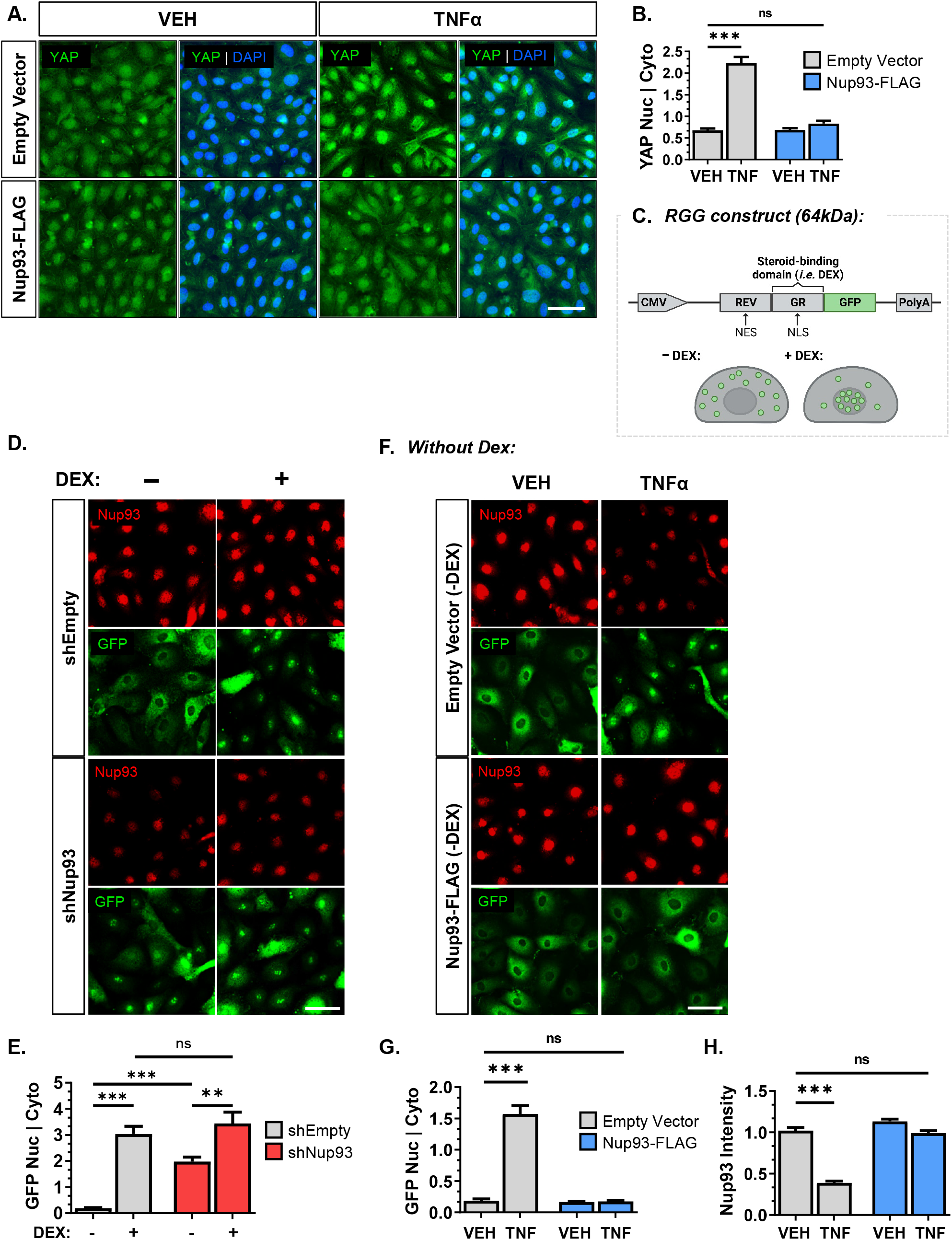
Endothelial loss of Nup93 leads to the nuclear accumulation of cargo protein. **(A)** HRECs were first exposed to the chronic inflammation model followed by lentiviral-mediated expression of exogenous Nup93, as detailed in Figure 3I. Restoring Nup93 protein levels redistributes Yap to the cytoplasm as shown via immunofluorescence, similar to healthy ECs. Quantified in **(B)**. **(C)** Schematic of the RGG expression construct. The RGG plasmid contains the full-length Rev sequence and the hormone-responsive element from the glucocorticoid receptor (GR). Dexamethasone (DEX) treatment leads to nuclear localization of the construct, as visualized by the GFP tag. Adapted from Love & Sweitzer, PNAS, 1998. **(D)** HRECS (shEmpty & shNup93) were further transduced with the RGG construct using lentiviral methods. HRECs were treated with DEX (1µM, 30min) where indicated. Immunofluorescence imaging of GFP signal indicates a baseline increase in nuclear GFP in shNup93 HRECs without DEX exposure. GFP nuclear-to-cytoplasmic intensities quantified in **(E)**. **(F)** HRECs were transduced with the RGG construct using lentiviral methods. Transduced cells were then exposed to the chronic inflammation model followed by lentiviral-mediated expression of exogenous Nup93. GFP nuclear-to-cytoplasmic intensities quantified in **(G)**. Nup93 immunofluorescence intensity quantified in **(H)**. Scale bar=50µm. n=3, *\*\*\* p<0.001, ** p<0.01*

To examine the functional consequence of chronic inflammation on endothelial NPC transport properties, ECs expressing the RGG construct were subject to chronic inflammation (TNFα [10ng/mL]; 6 days). In line with immunoblot results (**Figure 1I**), long-term exposure to TNFα significantly reduced endogenous Nup93 signal intensity (**Figure 6F&6H**). While healthy VEH treated ECs again showed a predominant cytoplasmic distribution of the GFP signal, long-term endothelial exposure to TNFα significantly increased nuclear GFP intensity in the absence of DEX (**Figure 6F&6G**). Treatment with DEX did not noticeably alter GFP distribution in chronically inflamed ECs (**Figure S6A&S6B**). We found that restoring Nup93 levels in already senescent ECs reversed the molecular hallmarks of aging (**Figure 3J-3O**). We therefore reasoned that restoration of Nup93 could recover NPC function in TNF-treated ECs. To test this, HRECs were first exposed to the chronic inflammation model followed by lentiviral infection of Nup93 (schematic in **Figure 3I**). Moderate overexpression of Nup93 using lentiviral methods did not affect the baseline distribution of the RGG construct; cytoplasmic GFP signal was indistinguishable when comparing Empty vector and Nup93-transduced vehicle-treated ECs (**Figure 6F&6G**). Overexpression of Nup93 protein in TNF-exposed ECs, however, led to a remarkable restoration of the cytoplasmic GFP signal, and the nuclear to cytoplasmic ratio was comparable to healthy ECs (**Figure 6F&6G**). Moreover, NPC transport properties remain intact, as treatment with DEX led to increased nuclear GFP signal, similar to healthy ECs (**Figure S6A&S6B**). These observations suggest that endothelial senescence and associated loss of Nup93 alters NPC transport properties for enhanced import and/or impaired export.

## DISCUSSION

Beyond its traditional role in cellular compartmentalization, the nuclear envelope has become a subject of interest as a novel regulator of EC function^41–43^. Our *in vitro* and *in vivo* results collectively demonstrate loss of NPC components as a major contributor to endothelial senescence. While endothelial senescence has been long-recognized in the development of CVD, the underlying mechanisms remain incompletely understood. Highlighting the novelty of our findings, our research stands as the first documentation of NPC dysregulation in EC activation and vascular aging. Here, we demonstrate an age-associated decline in endothelial Nup93 protein level, where inflammation-induced loss of Nup93 triggers aberrant signaling involved in EC senescence. Human aging and chronic disease are associated with inflammation driven by systemic proinflammatory cytokines. As the innermost lining of the vasculature, ECs are the first line of exposure to circulating proinflammatory agents and other insults. While we currently do not know the mechanism(s) by which inflammation leads to endothelial Nup93 protein depletion, our findings align with prior reports that highlight the vulnerability of Nup93 to oxidative protein damage^13^, a characteristic associated with inflammation and aging. This raises the possibility that proinflammatory factors degrade distinct endothelial NPC components as a means of vascular damage. Further studies, however, are necessary to fully understand the process by which inflammation targets endothelial NPC components.

ECs are also particularly vulnerable to the physiological changes caused by hemodynamic flow patterns. Physiological shear stress is well-known to induce profound changes in endothelial cellular shape that is coupled with drastic changes in the EC transcriptome^30^. These endothelial adaptations rely on both actin remodeling and the proper localization of mechanosensitive TFs to mediate a coordinated biological response to flow^31,44^. In the established adult vasculature, Yap is one of the most robust mechanosensitive transcriptional cofactors driving the detrimental effects of atheroprone flow^4,5^. Increases in F-actin formation also promote nuclear Yap localization, demonstrating the mechanical susceptibility of Yap signaling^45,46^. Moreover, mechanical forces transmitted to the nuclear envelope can affect the NPC diameter to regulate Yap nucleocytoplasmic transport^47^. Multiple studies have reported increased Yap activity in atheroprone flow conditions using both *in vitro* and *in vivo* models. Both diet-induced dyslipidemia and aging models have also been reported to increase endothelial and vascular stiffening^48,49^, further supporting the mechanical regulation of pathological Yap signaling. In light of recent studies implicating Nup93 in actin cytoskeletal remodeling^50^, Nup93 loss in ECs may also trigger F-actin formation for endothelial stiffening. Hence, the cytoskeletal and transcriptomic changes associated with endothelial dysfunction may reflect aberrations at the NPC, a topic to be explored in future work.

The loss of Nup93 likely impacts the cellular localization of other TFs, as demonstrated in our own data as well as others^51–53^. Our collection of studies, however, identifies Yap signaling as the major consequence of endothelial Nup93 loss. As a transcriptional cofactor, Yap can co-activate multiple inflammation-associated TFs to mediate a synchronized response and may account for the prominent Yap effect seen herein^34–36,54^. The role of Yap signaling in the natural aging process, however, remains controversial. Several *in vitro* studies using various cell types show Yap to either inhibit or promote the expression of established senescence markers^55^. While aging is a general process that impacts all cells, the role of Yap may be entirely cell-type dependent. Our data indicates a causal relationship between increased Yap and endothelial aging, consistent with previous reports in primary ECs^3^. Moreover, our studies further identify nucleoporins in the regulation of Yap signaling to provide incremental evidence that reveal a connection between Yap signaling and nuclear envelope components^55,56^.

As components of the NPC, nucleoporins are critical for proper nucleocytoplasmic transport in all cells. The critical nature of Nup93 in NPC transport function is not surprising, as several groups have previously reported NPC transport defects^13,57^. Our studies using the hormone-inducible GFP reporter construct not only substantiates the importance of Nup93 in establishing the NPC diffusion barrier, but are the first to detail the consequences of nucleoporin loss in primary ECs. We show that unlike the cytoplasmic GFP distribution observed in healthy ECs, endothelial loss of Nup93 results in a prominent nuclear GFP reporter signal. Given the cytoplasmic preference for the GFP reporter in steroid-deficient conditions, an equilibrium of GFP signal between the cytoplasm and nucleus would be expected if loss of Nup93 produced more passively permissive NPCs. The distinct shift toward a nuclear GFP signal, however, suggests that Nup93 loss may impair regulatory features of active transport to collectively reduce cargo export and/or heighten import. Our observations support recent findings implicating age-associated nuclear compartmentalization^15,16^. While the underlying mechanisms warrant future investigation, our results establish suboptimal levels of Nup93, and likely NPC transport issues, as a key feature of EC senescence. Moreover, our results establish Nup93 restoration as a feasible approach for reprogramming senescent ECs toward favorable health.

The clinical implications of senescence in chronic human diseases have launched research into senolytics, or therapies that selectively clear and remove senescent cells. It remains unclear, however, whether removal of senescent cells is truly beneficial. Prior studies using senolytics indicate that failure to replace ECs instead leads to debilitating consequences^58^. While still an area of controversy, the idea of reprogramming senescent cells would instead be favorable. We find that restoring Nup93 protein expression in senescent ECs reverses features of cellular aging. It is therefore intriguing to propose that endothelial dysfunction may be initiated with NPC degradation. Endothelial NPC dysregulation may activate an initial inflammatory defense mechanism, where the inability to repair NPCs leads to a misguided response resulting in perpetual inflammation and consequent EC dysfunction. As a last resort, cellular senescence programs may be prompted to protect the endothelium against irreparable damage. Our findings suggest that reestablishing NPC integrity may serve as a potential senotherapeutic to restore endothelial health and mitigate vascular disease.

In summary, our study is the first to implicate NPC components in endothelial and vascular health. Using established *in vitro* models of endothelial aging, the present study reveals the vulnerability of endothelial Nup93 to chronic inflammation, thereby contributing to the newly emerging concept that nuclear envelope proteins function as novel regulators of EC behavior. We find Yap activation as a key driver of inflammation-induced EC senescence and substantiate the pathogenic role of Yap signaling in the established vasculature. Lastly, our studies indicate that restoring Nup93 protein levels is sufficient for endothelial reprogramming toward healthy ECs. These results collectively reveal NPC dysregulation as a biomarker of endothelial senescence, where restoring Nup93 to baseline levels could serve as a novel and promising senotherapeutic approach.

## Supporting information

SuppFig

## ACKNOWLEDGEMENTS

We thank Patrick Lusk, Ph.D. (Yale University) for his invaluable advice on NPC biology and Yuwei Jiang, Ph.D. (UIC) for providing aged mouse tissue. This work was supported in part by the National Institutes of Health (R00 HL130581 to M.Y.L.) and the American Heart Association (AHA 941057 to M.Y.L.).

## AUTHOR CONTRIBUTIONS

T.D.N performed the gene expression/pathway analyses, and performed most of the curation, visualization, and formal analyses of the data presented herein. S.P.D performed the RT-qPCR validation experiments. M.K.R performed the monocyte adhesion assays. J.M.B. performed the SA-βGal assays. J.M. harvested and sectioned mouse tissues for immunostaining. M.A.W generated constructs and reagents. M.Y.L conceived and directed the project. T.D.N and M.Y.L wrote the manuscript, which was reviewed and edited by all authors.

## Notes

### Competing Interest Statement

The authors have declared no competing interest.

