## Supplementary material for "Nucleoporin93 (Nup93) Limits Yap Activity to Prevent Endothelial Cell Senescence": SuppFig

Figure S1

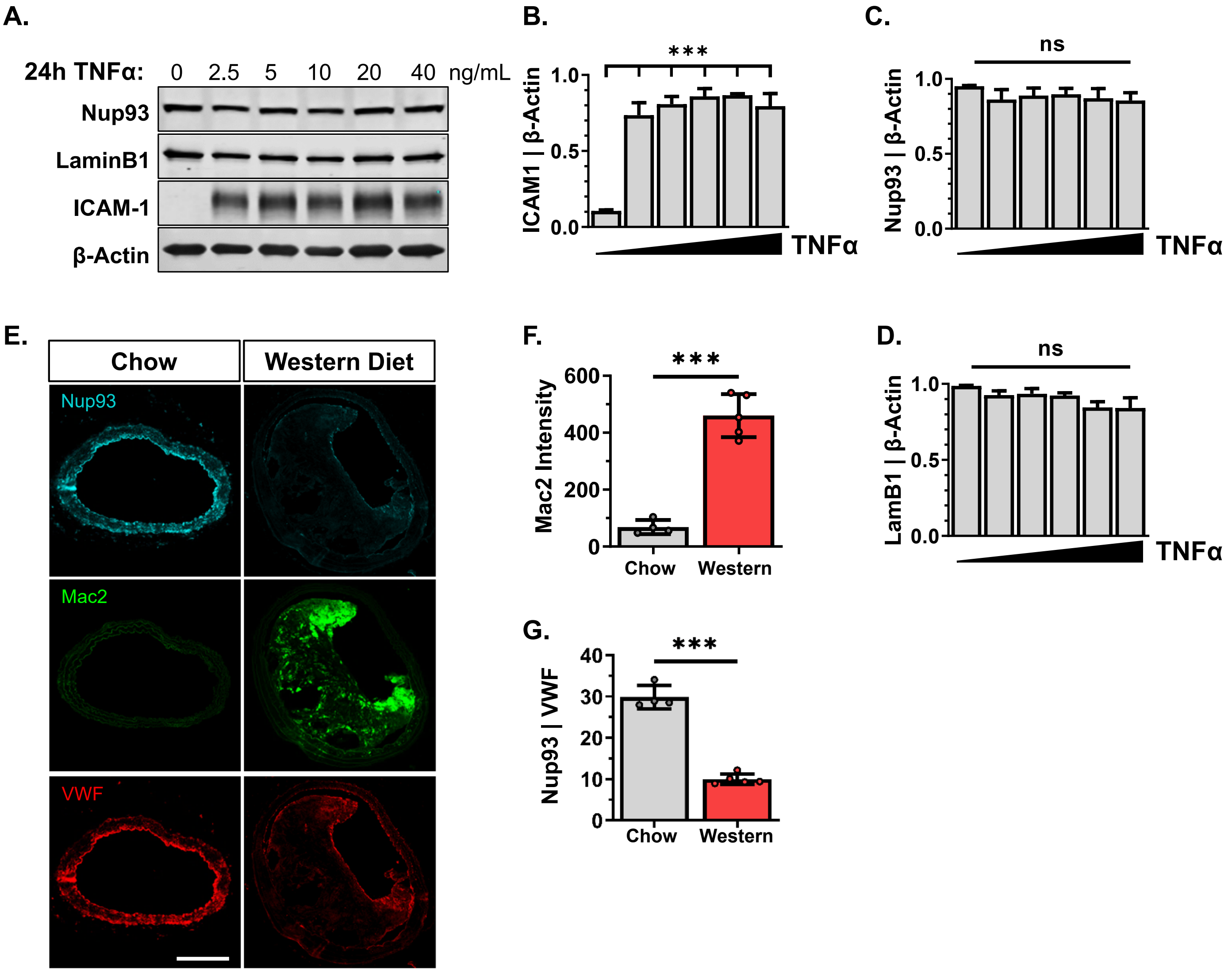

A.
24h TNFα: 0 2.5 5 10 20 40 ng/mL
Nup93
LaminB1
ICAM-1
β-Actin
B.
***
ICAM1 | β-Actin
1.0
0.5
0.0
TNFα
C.
ns
Nup93 | β-Actin
1.0
0.5
0.0
TNFα
E.
Chow
Western Diet
Nup93
Mac2
VWF
F.
***
Mac2 Intensity
600
400
200
0
Chow
Western
D.
ns
LamB1 | β-Actin
1.0
0.5
0.0
TNFα
G.
***
Nup93 | VWF
40
30
20
10
0
Chow
Western

**Figure S2**

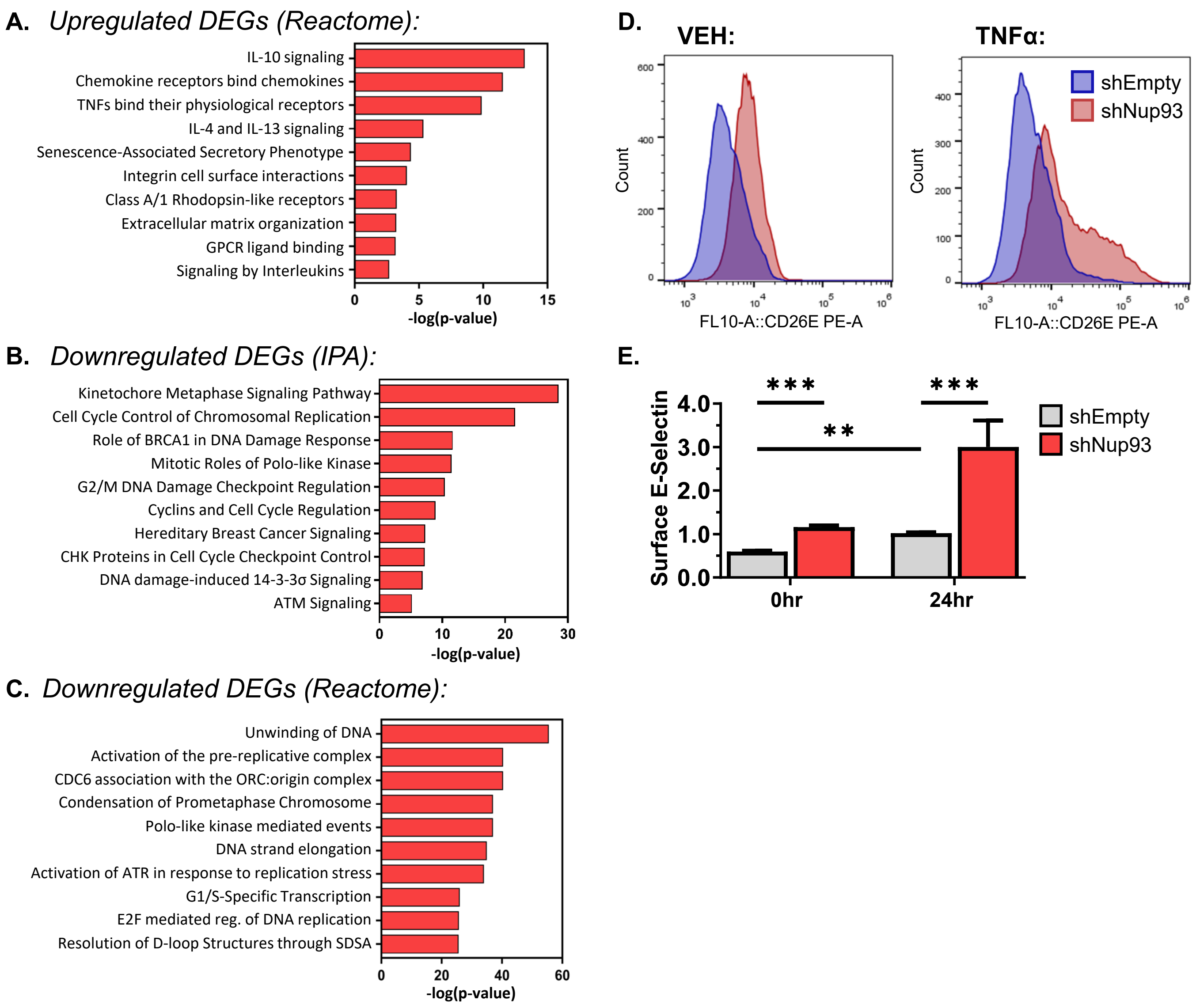
A. Upregulated DEGs (Reactome):
IL-10 signaling
Chemokine receptors bind chemokines
TNFs bind their physiological receptors
IL-4 and IL-13 signaling
Senescence-Associated Secretory Phenotype
Integrin cell surface interactions
Class A/1 Rhodopsin-like receptors
Extracellular matrix organization
GPCR ligand binding
Signaling by Interleukins
0 5 10 15
-log(p-value)
B. Downregulated DEGs (IPA):
Kinetochore Metaphase Signaling Pathway
Cell Cycle Control of Chromosomal Replication
Role of BRCA1 in DNA Damage Response
Mitotic Roles of Polo-like Kinase
G2/M DNA Damage Checkpoint Regulation
Cyclins and Cell Cycle Regulation
Hereditary Breast Cancer Signaling
CHK Proteins in Cell Cycle Checkpoint Control
DNA damage-induced 14-3-3σ Signaling
ATM Signaling
0 10 20 30
-log(p-value)
C. Downregulated DEGs (Reactome):
Unwinding of DNA
Activation of the pre-replicative complex
CDC6 association with the ORC:origin complex
Condensation of Prometaphase Chromosome
Polo-like kinase mediated events
DNA strand elongation
Activation of ATR in response to replication stress
G1/S-Specific Transcription
E2F mediated reg. of DNA replication
Resolution of D-loop Structures through SDSA
0 20 40 60
-log(p-value)
D. VEH:
TNFα:
shEmpty
shNup93
Count
FL10-A::CD26E PE-A
E.
Surface E-Selectin
***
**
***
4.0
3.0
2.0
1.0
0.0
0hr
24hr
shEmpty
shNup93

**Figure S3**

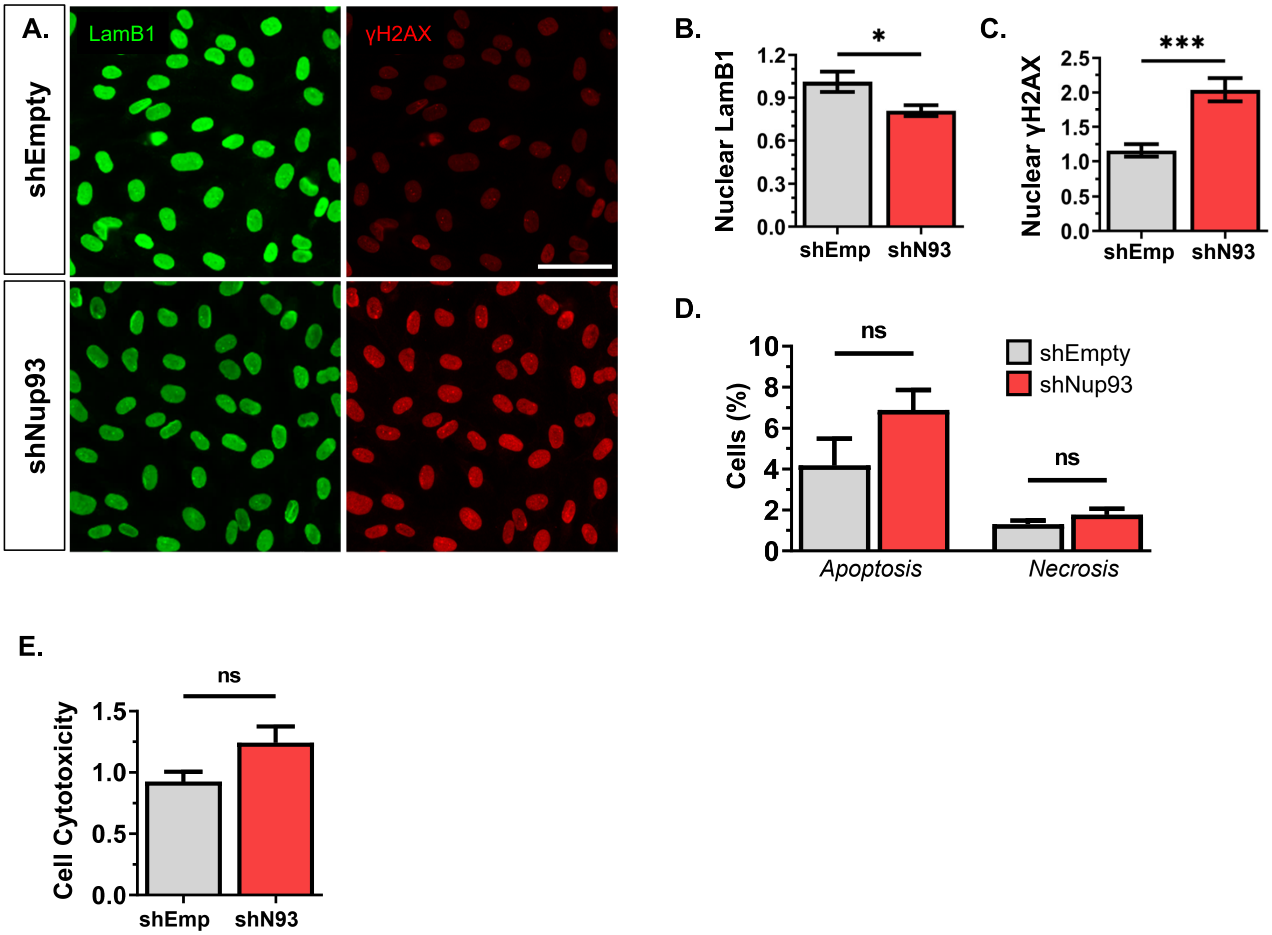

A.
LamB1
γH2AX
shEmpty
shNup93
B.
Nuclear LamB1
*
1.2
0.9
0.6
0.3
0.0
shEmp
shN93
C.
Nuclear γH2AX
***
2.5
2.0
1.5
1.0
0.5
0.0
shEmp
shN93
D.
Cells (%)
ns
ns
10
8
6
4
2
0
shEmpty
shNup93
Apoptosis
Necrosis
E.
Cell Cytotoxicity
ns
1.5
1.0
0.5
0.0
shEmp
shN93

### Figure S4

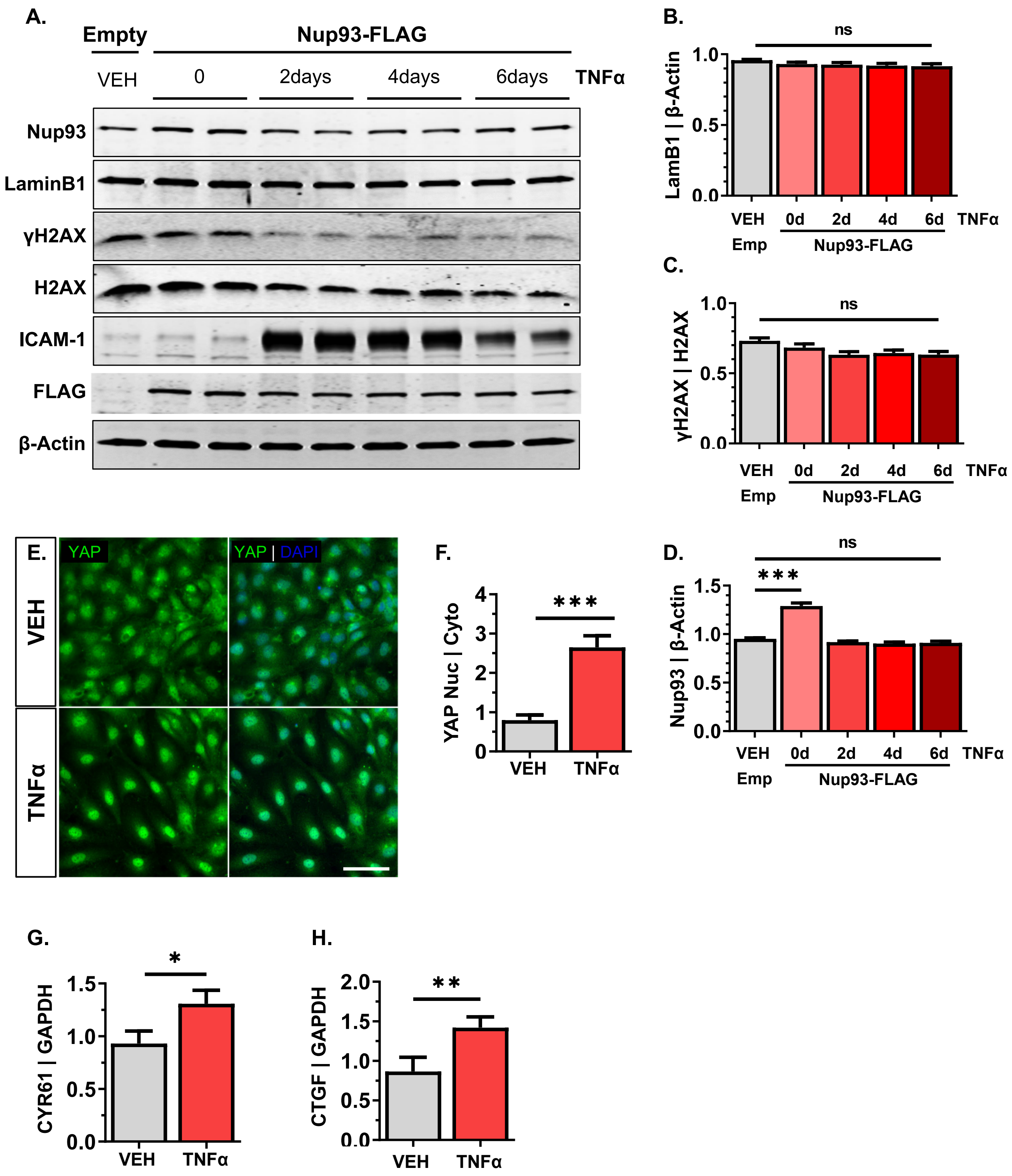
A.
Empty
Nup93-FLAG
VEH
0
2days
4days
6days
TNFα
Nup93
LaminB1
γH2AX
H2AX
ICAM-1
FLAG
β-Actin
B.
ns
LamB1 | β-Actin
VEH
0d
2d
4d
6d
TNFα
Emp
Nup93-FLAG
C.
ns
γH2AX | H2AX
D.
ns
***
Nup93 | β-Actin
E.
YAP
YAP | DAPI
VEH
TNFα
F.
***
YAP Nuc | Cyto
G.
*
CYR61 | GAPDH
H.
**
CTGF | GAPDH

### Figure S5

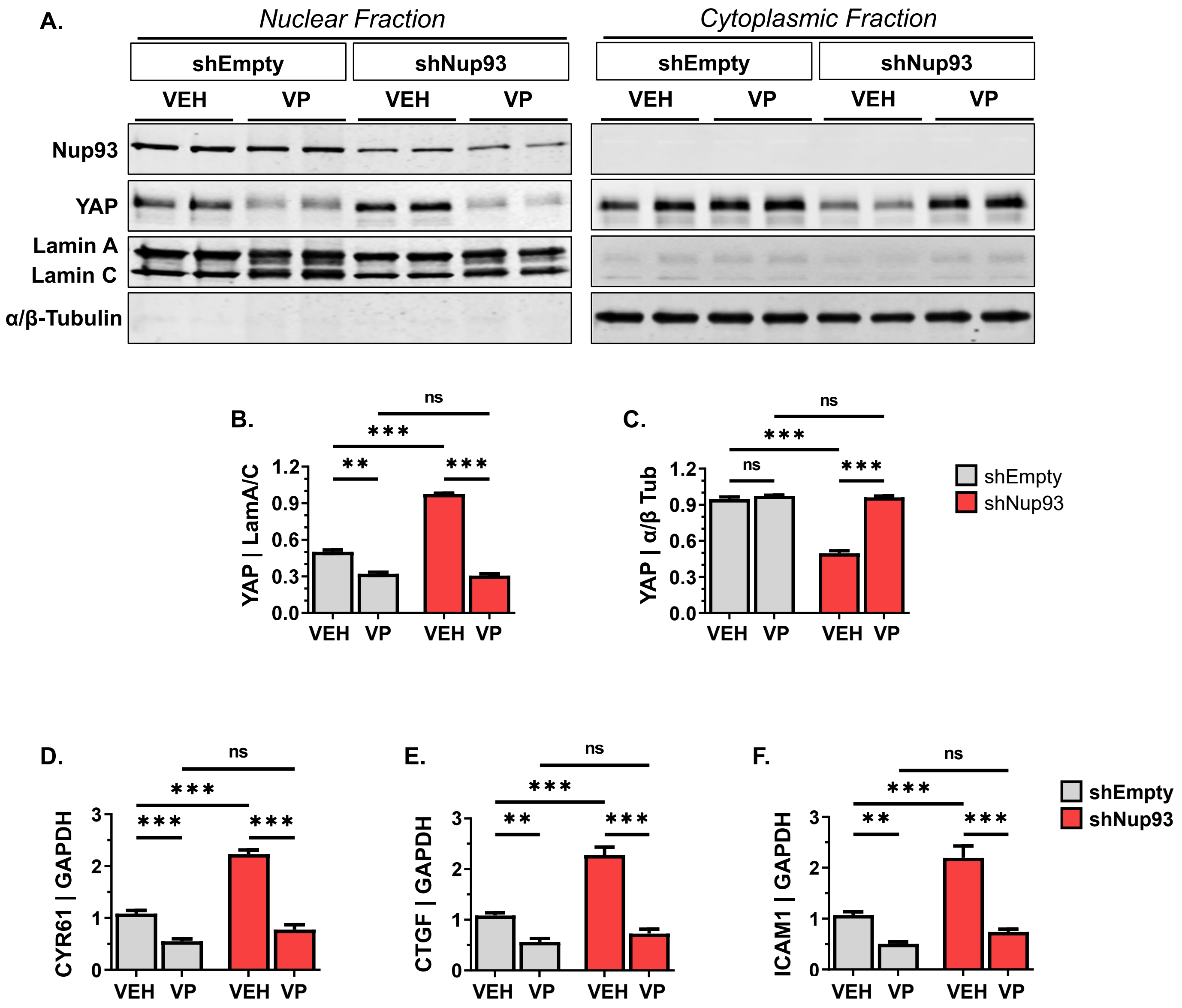

A.
Nuclear Fraction
Cytoplasmic Fraction
shEmpty
shNup93
VEH
VP
Nup93
YAP
Lamin A
Lamin C
α/β-Tubulin
B.
YAP | LamA/C
C.
YAP | α/β Tub
shEmpty
shNup93
ns
D.
CYR61 | GAPDH
E.
CTGF | GAPDH
F.
ICAM1 | GAPDH

### Figure S6

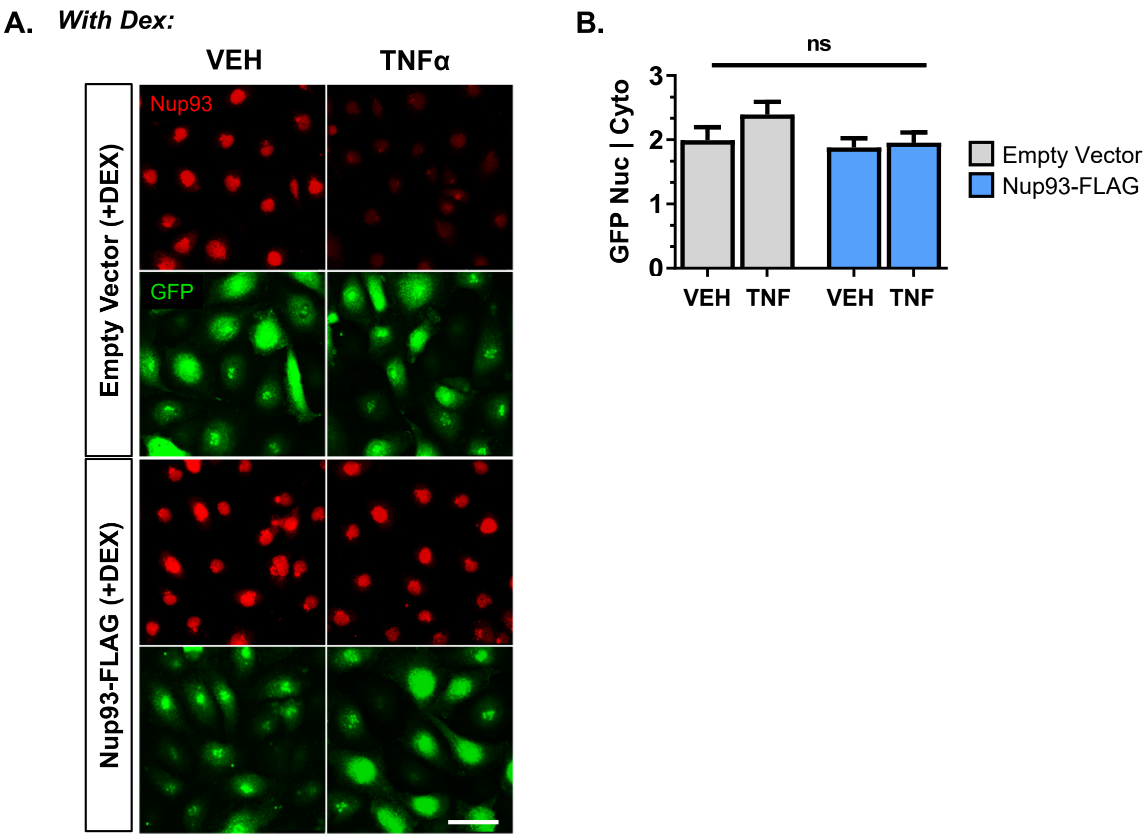

A.
With Dex:
VEH
TNFα
Empty Vector (+DEX)
Nup93-FLAG (+DEX)
Nup93
GFP
B.
ns
GFP Nuc | Cyto
0
1
2
3
VEH
TNF
VEH
TNF
Empty Vector
Nup93-FLAG

**Table S1. RT-qPCR primers for detection of human transcripts**

| Target gene | Forward sequence (5'→3') | Reverse sequence (5'→3') | Product Size (bp) |
|---|---|---|---|
| *CYR61* | TGAAGCGGCTCCCTGTTTT | CGGGTTTCTTTCACAAGGCG | 175 |
| *CTGF* | ACCGACTGGAAGACACGTTTG | CCAGGTCAGCTTCGCAAGG | 192 |
| *ICAM1* | TTGGGCATAGAGACCCCGTT | GCACATTGCTCAGTTCATACACC | 93 |
| *VCAM1* | CAGTAAGGCAGGCTGTAAAAGA | TGGAGCTGGTAGACCCTCG | 144 |
| *GAPDH* | CTCTCTGCTCCTCCTGTTCGAC | TGAGCGATGTGGCTCGGCT | 71 |

**Table S2. Differentially expressed genes in shNup93-transduced primary HRECs (compared to shEmpty control).**

| Gene_ID | shNup93 A | shNup93 B | shNup93 C | shEmpty A | shEmpty B | shEmpty C | log2FoldChange | padj | GeneName |
|---|---|---|---|---|---|---|---|---|---|
| ENSG00000114812 | 22.32649827 | 13.55212335 | 28.12189076 | 0 | 0 | 0 | 6.936330717 | 1.91E-06 | VIPR1 |
| ENSG00000137203 | 23.3413391 | 9.382239244 | 16.22416775 | 0 | 0 | 0 | 6.551147929 | 1.77E-05 | TFAP2A |
| ENSG00000183160 | 19.28197578 | 8.339768217 | 19.4690013 | 0 | 0 | 0 | 6.494275047 | 2.39E-05 | TMEM119 |
| ENSG00000179399 | 56.83108651 | 40.65637006 | 40.01961377 | 0 | 0 | 2.000122788 | 6.179650147 | 2.12E-07 | GPC5 |
| ENSG00000183090 | 15.22261246 | 5.212355136 | 11.89772301 | 0 | 0 | 0 | 5.952561525 | 0.00035493 | FREM3 |
| ENSG00000015520 | 14.20777163 | 23.97683362 | 15.14255656 | 0.879212552 | 0 | 0 | 5.712135973 | 0.000160985 | NPC1L1 |
| ENSG00000006210 | 155.2706471 | 115.714284 | 122.2220637 | 1.758425104 | 2.892657035 | 3.000184182 | 5.699468603 | 3.60E-22 | CX3CL1 |
| ENSG00000163606 | 2.029681661 | 8.339768217 | 16.22416775 | 0 | 0 | 0 | 5.666727829 | 0.001742966 | CD200R1 |
| ENSG00000215914 | 3.044522491 | 9.382239244 | 11.89772301 | 0 | 0 | 0 | 5.538978806 | 0.002212355 | MMP23A |
| ENSG00000130294 | 10.1484083 | 8.339768217 | 5.408055916 | 0 | 0 | 0 | 5.517327128 | 0.001966916 | KIF1A |
| ENSG00000233930 | 7.103885813 | 13.55212335 | 2.163222366 | 0 | 0 | 0 | 5.451012743 | 0.003601496 | KRTAP5-AS1 |
| ENSG00000043462 | 9.133567474 | 10.42471027 | 23.79544603 | 0.879212552 | 0 | 0 | 5.410841986 | 0.000674412 | LCP2 |
| ENSG00000198771 | 5.074204152 | 9.382239244 | 7.571278282 | 0 | 0 | 0 | 5.397630586 | 0.003130131 | RCSD1 |
| ENSG00000163121 | 8.118726644 | 3.127413081 | 9.734500648 | 0 | 0 | 0 | 5.327327452 | 0.004581153 | NEURL3 |
| ENSG00000224294 | 9.133567474 | 7.29729719 | 4.326444732 | 0 | 0 | 0 | 5.314402137 | 0.004395954 | PINCR |
| ENSG00000104490 | 10.1484083 | 9.382239244 | 1.081611183 | 0 | 0 | 0 | 5.305471582 | 0.006816555 | NCALD |
| ENSG00000070985 | 11.16324914 | 8.339768217 | 1.081611183 | 0 | 0 | 0 | 5.303788972 | 0.006918151 | TRPM5 |
| ENSG00000183114 | 9.133567474 | 1.042471027 | 9.734500648 | 0 | 0 | 0 | 5.252078773 | 0.008075344 | FAM43B |
| ENSG00000038945 | 8.118726644 | 7.29729719 | 4.326444732 | 0 | 0 | 0 | 5.241824996 | 0.005619906 | MSR1 |
| ENSG00000132517 | 2.029681661 | 10.42471027 | 6.489667099 | 0 | 0 | 0 | 5.179416518 | 0.008566015 | SLC52A1 |
| ENSG00000007314 | 15.22261246 | 5.212355136 | 16.22416775 | 0.879212552 | 0 | 0 | 5.170990293 | 0.001717142 | SCN4A |
| ENSG00000137491 | 45.66783737 | 51.08108033 | 59.48861507 | 0.879212552 | 3.856876047 | 0 | 5.045747142 | 3.34E-10 | SLCO2B1 |
| ENSG00000249464 | 20.29681661 | 16.67953643 | 22.71383485 | 0.879212552 | 0 | 1.000061394 | 4.981906613 | 0.000183171 | LINC01091 |
| ENSG00000166592 | 47.69751903 | 51.08108033 | 46.50928087 | 1.758425104 | 0 | 3.000184182 | 4.94115196 | 8.99E-10 | RRAD |
| ENSG00000135253 | 12.17808997 | 5.212355136 | 11.89772301 | 0.879212552 | 0 | 0 | 4.847441106 | 0.004943397 | KCP |
| ENSG00000074317 | 13.1929308 | 19.80694952 | 18.38739011 | 1.758425104 | 0 | 0 | 4.781150748 | 0.000572388 | SNCB |
| ENSG00000115607 | 49.72720069 | 53.16602238 | 51.91733679 | 2.637637656 | 0.964219012 | 2.000122788 | 4.77368359 | 1.30E-10 | IL18RAP |
| ENSG00000261594 | 89.30599308 | 63.59073266 | 97.34500648 | 3.516850207 | 1.928438024 | 4.000245576 | 4.726672573 | 3.14E-16 | TPBGL |
| ENSG00000110324 | 17.25229412 | 7.29729719 | 14.06094538 | 0.879212552 | 0 | 1.000061394 | 4.354896239 | 0.003217505 | IL10RA |
| ENSG00000171724 | 91.33567474 | 43.78378314 | 76.794394 | 2.637637656 | 2.892657035 | 5.00030697 | 4.345207292 | 2.72E-13 | VAT1L |
| ENSG00000261239 | 64.94981315 | 40.65637006 | 43.26444732 | 5.275275311 | 0 | 2.000122788 | 4.313820405 | 1.07E-09 | ANKRD26P1 |
| ENSG00000175899 | 26.38586159 | 20.84942054 | 8.652889465 | 0.879212552 | 0.964219012 | 1.000061394 | 4.302506072 | 0.000318242 | A2M |
| ENSG00000110848 | 38.56395156 | 38.571428 | 52.99894797 | 0.879212552 | 4.821095059 | 1.000061394 | 4.292655722 | 7.64E-09 | CD69 |
| ENSG00000107562 | 47.69751903 | 47.95366725 | 88.69211701 | 3.516850207 | 0 | 6.000368364 | 4.281967078 | 2.83E-11 | CXCL12 |
| ENSG00000162981 | 32.47490658 | 23.97683362 | 17.30577893 | 0 | 0 | 4.000245576 | 4.267315558 | 3.73E-05 | FAM84A |
| ENSG00000112299 | 14.20777163 | 8.339768217 | 11.89772301 | 1.758425104 | 0 | 0 | 4.211324293 | 0.006085323 | VNN1 |
| ENSG00000268894 | 36.5342699 | 37.52895698 | 25.95866839 | 4.396062759 | 0 | 1.000061394 | 4.159226366 | 7.51E-07 | PLCE1-AS1 |
| ENSG00000164440 | 19.28197578 | 10.42471027 | 19.4690013 | 1.758425104 | 0.964219012 | 0 | 4.129718607 | 0.000742908 | TXLNB |
| ENSG00000196208 | 7.103885813 | 17.72200746 | 7.571278282 | 1.758425104 | 0 | 0 | 4.125043731 | 0.009394121 | GREB1 |
| ENSG00000049249 | 35.51942907 | 23.97683362 | 19.4690013 | 1.758425104 | 1.928438024 | 1.000061394 | 4.067092153 | 1.00E-05 | TNFRSF9 |
| ENSG00000173110 | 146.1370796 | 101.1196896 | 187.1187347 | 5.275275311 | 9.642190118 | 14.00085951 | 3.92718638 | 6.30E-24 | HSPA6 |
| ENSG00000267640 | 23.3413391 | 26.06177568 | 7.571278282 | 1.758425104 | 0 | 2.000122788 | 3.918606337 | 0.000453654 | AC016582.3 |
| ENSG00000105246 | 76.11306229 | 75.05791395 | 73.54956045 | 6.154487863 | 3.856876047 | 5.00030697 | 3.896272971 | 4.01E-15 | EBI3 |
| ENSG00000162643 | 9.133567474 | 18.76447849 | 14.06094538 | 0 | 0 | 3.000184182 | 3.863129364 | 0.003058912 | WDR63 |
| ENSG00000168453 | 20.29681661 | 8.339768217 | 11.89772301 | 0.879212552 | 0 | 2.000122788 | 3.833205273 | 0.003176896 | HR |
| ENSG00000170323 | 539.8953218 | 463.8996071 | 468.3376423 | 41.32298994 | 34.71188443 | 34.00208739 | 3.737316218 | 3.32E-88 | FABP4 |
| ENSG00000080709 | 19.28197578 | 20.84942054 | 20.55061248 | 1.758425104 | 1.928438024 | 1.000061394 | 3.68570734 | 0.000145561 | KCNN2 |
| ENSG00000107105 | 16.23745329 | 10.42471027 | 9.734500648 | 0 | 0.964219012 | 2.000122788 | 3.664944733 | 0.005641466 | ELAVL2 |
| ENSG00000143869 | 1138.651412 | 938.2239244 | 883.6763366 | 86.16283008 | 76.17330193 | 73.00448176 | 3.650096433 | 5.09E-146 | GDF7 |
| ENSG00000007908 | 981.3510831 | 757.8764367 | 949.6546188 | 79.12912967 | 85.81549205 | 67.00411339 | 3.533155639 | 1.38E-128 | SELE |
| ENSG00000206337 | 40.59363322 | 25.01930465 | 27.04027958 | 4.396062759 | 2.892657035 | 1.000061394 | 3.458453072 | 4.73E-06 | HCP5 |
| ENSG00000181634 | 2041.859751 | 1737.799202 | 1818.188399 | 193.4267614 | 150.4181658 | 173.0106211 | 3.4342239 | 5.10E-250 | TNFSF15 |
| ENSG00000064300 | 67.99433564 | 69.84555882 | 63.8150598 | 7.912912967 | 4.821095059 | 6.000368364 | 3.419564745 | 9.58E-13 | NGFR |
| ENSG00000102109 | 12.17808997 | 29.18918876 | 8.652889465 | 0.879212552 | 0 | 4.000245576 | 3.387194711 | 0.002692348 | PCSK1N |
| ENSG00000154928 | 151.2112837 | 148.0308859 | 142.7726762 | 15.82582593 | 15.42750419 | 13.00079812 | 3.316758888 | 1.22E-26 | EPHB1 |
| ENSG00000171246 | 27.40070242 | 9.382239244 | 29.20350194 | 1.758425104 | 0.964219012 | 4.000245576 | 3.30925515 | 0.000361463 | NPTX1 |
| ENSG00000004468 | 17.25229412 | 12.50965233 | 7.571278282 | 0.879212552 | 2.892657035 | 0 | 3.306634156 | 0.008218277 | CD38 |
| ENSG00000113645 | 12.17808997 | 15.63706541 | 8.652889465 | 1.758425104 | 1.928438024 | 0 | 3.282945308 | 0.008018466 | WWC1 |
| ENSG00000186827 | 18.26713495 | 21.89189157 | 23.79544603 | 2.637637656 | 1.928438024 | 2.000122788 | 3.27588472 | 0.000180361 | TNFRSF4 |
| ENSG00000105825 | 1109.221028 | 1084.169868 | 1087.019239 | 116.0560568 | 104.1356533 | 123.0075515 | 3.257233846 | 5.82E-181 | TFPI2 |
| ENSG00000160223 | 32.47490658 | 57.33590649 | 35.69316904 | 2.637637656 | 3.856876047 | 7.000429757 | 3.235179789 | 2.31E-07 | ICOSLG |
| ENSG00000157404 | 375.4911073 | 453.4748968 | 334.2178556 | 39.56456483 | 36.64032245 | 48.00294691 | 3.230405395 | 5.77E-58 | KIT |
| ENSG00000163735 | 347.075564 | 265.8301119 | 338.5443003 | 37.80613973 | 28.92657035 | 37.00227158 | 3.195425936 | 5.81E-49 | CXCL5 |
| ENSG00000019582 | 53.78656402 | 63.59073266 | 43.26444732 | 5.275275311 | 2.892657035 | 10.00061394 | 3.155650159 | 4.46E-09 | CD74 |
| ENSG00000056558 | 519.5985052 | 527.4903397 | 587.3148724 | 63.30330373 | 57.85314071 | 68.00417479 | 3.111473758 | 1.08E-86 | TRAF1 |
| ENSG00000007129 | 45.66783737 | 34.4015439 | 25.95866839 | 3.516850207 | 3.856876047 | 5.00030697 | 3.107196203 | 2.10E-06 | CEACAM21 |
| ENSG00000069122 | 905.2380208 | 694.2857041 | 726.842715 | 95.83416815 | 69.42376885 | 105.0064464 | 3.106043861 | 8.06E-92 | ADGRF5 |
| ENSG00000141668 | 69.00917647 | 71.93050087 | 85.44728347 | 12.30897573 | 8.677971106 | 5.00030697 | 3.105880214 | 1.19E-12 | CBLN2 |
| ENSG00000107719 | 587.5928408 | 513.9382164 | 522.4182014 | 65.94094139 | 66.53111182 | 58.00356085 | 3.090130572 | 4.61E-85 | PALD1 |
| ENSG00000102760 | 594.6967267 | 581.6988331 | 612.1919296 | 70.33700415 | 63.63845478 | 81.00497291 | 3.058381823 | 1.40E-94 | RGCC |
| ENSG00000205710 | 27.40070242 | 21.89189157 | 28.12189076 | 2.637637656 | 4.821095059 | 2.000122788 | 3.033319267 | 6.88E-05 | C17orf107 |
| ENSG00000237988 | 55.81624568 | 74.01544293 | 60.57022625 | 12.30897573 | 2.892657035 | 8.000491151 | 3.021345845 | 5.60E-10 | OR2I1P |
| ENSG00000174059 | 937.7129274 | 872.5482497 | 879.3498919 | 124.8481824 | 86.77971106 | 122.0074901 | 3.009093784 | 3.32E-122 | CD34 |
| ENSG00000162692 | 1244.194858 | 1078.957513 | 1263.321862 | 146.8284962 | 127.2769096 | 175.0107439 | 2.999300029 | 8.13E-146 | VCAM1 |
| ENSG00000163083 | 25.37102076 | 30.23165979 | 19.4690013 | 1.758425104 | 3.856876047 | 4.000245576 | 2.981089304 | 0.000114807 | INHBB |
| ENSG00000137285 | 14.20777163 | 3.127413081 | 42.18283614 | 0.879212552 | 2.892657035 | 4.000245576 | 2.953065456 | 0.005748285 | TUBB2B |
| ENSG00000115008 | 51.75688235 | 25.01930465 | 29.20350194 | 4.396062759 | 5.785314071 | 4.000245576 | 2.903537904 | 8.16E-06 | IL1A |
| ENSG00000120337 | 667.7562665 | 486.8339697 | 634.9057645 | 99.35101836 | 69.42376885 | 70.00429757 | 2.899674803 | 1.06E-65 | TNFSF18 |
| ENSG00000142235 | 55.81624568 | 37.52895698 | 87.61050583 | 6.154487863 | 9.642190118 | 9.000552545 | 2.873084265 | 1.37E-08 | LMTK3 |
| ENSG00000185607 | 8.118726644 | 33.35907287 | 25.95866839 | 1.758425104 | 4.821095059 | 3.000184182 | 2.824074691 | 0.001231733 | ACTBP7 |
| ENSG00000135477 | 20.29681661 | 11.4671813 | 15.14255656 | 0.879212552 | 5.785314071 | 0 | 2.821086118 | 0.008374621 | KRT87P |
| ENSG00000108932 | 62.92013149 | 108.4169868 | 85.44728347 | 11.42976317 | 17.35594221 | 8.000491151 | 2.801109597 | 8.05E-12 | SLC16A6 |
| ENSG00000121966 | 181.6565087 | 161.5830092 | 169.8129557 | 27.25558911 | 26.99813233 | 20.00122788 | 2.784032409 | 9.76E-26 | CXCR4 |
| ENSG00000108691 | 3892.929426 | 3586.100333 | 3582.296238 | 538.9572943 | 514.8929523 | 559.0343192 | 2.77824421 | 0 | CCL2 |
| ENSG00000137726 | 29.43038408 | 21.89189157 | 30.28511313 | 5.275275311 | 4.821095059 | 2.000122788 | 2.740610247 | 0.00012197 | FXYD6 |
| ENSG00000110876 | 58.86076817 | 65.67567471 | 49.75411442 | 14.94661338 | 3.856876047 | 7.000429757 | 2.736780041 | 3.59E-08 | SELPLG |
| ENSG00000136002 | 13.1929308 | 17.72200746 | 19.4690013 | 1.758425104 | 2.892657035 | 3.000184182 | 2.728329109 | 0.003167921 | ARHGEF4 |
| ENSG00000274615 | 127.8699446 | 119.8841681 | 67.05989335 | 20.22188869 | 19.28438024 | 8.000491151 | 2.719464136 | 2.90E-12 | AC233968.1 |
| ENSG00000232480 | 32.47490658 | 42.74131211 | 62.73344862 | 5.275275311 | 4.821095059 | 11.00067533 | 2.718825596 | 1.20E-06 | TGFB2-AS1 |
| ENSG00000166670 | 70.0240173 | 59.42084855 | 70.3047269 | 8.792125519 | 7.713752095 | 14.00085951 | 2.71835041 | 3.83E-10 | MMP10 |

| | | | | | | | | | |
|---|---|---|---|---|---|---|---|---|---|
| ENSG00000189056 | 364.3278581 | 277.2972932 | 297.4430754 | 36.04771463 | 36.64032245 | 71.00435897 | 2.717360675 | 2.44E-32 | RELN |
| ENSG00000140323 | 87.27631142 | 95.9073345 | 108.1611183 | 12.30897573 | 16.3917232 | 16.0009823 | 2.709021228 | 1.49E-14 | DISP2 |
| ENSG00000162490 | 66.97949481 | 61.5057906 | 81.12083873 | 12.30897573 | 7.713752095 | 13.00079812 | 2.664244286 | 2.57E-10 | DRAXIN |
| ENSG00000124257 | 27.40070242 | 31.27413081 | 31.36672431 | 4.396062759 | 3.856876047 | 6.000368364 | 2.663630996 | 4.83E-05 | NEURL2 |
| ENSG00000198576 | 5.074204152 | 25.01930465 | 29.20350194 | 5.275275311 | 0 | 4.000245576 | 2.661955491 | 0.007968275 | ARC |
| ENSG00000182985 | 208.0423702 | 183.4749008 | 160.0784551 | 29.01401421 | 36.64032245 | 22.00135067 | 2.652204881 | 3.32E-24 | CADM1 |
| ENSG00000048052 | 88.29115225 | 70.88802985 | 100.58984 | 6.154487863 | 15.42750419 | 21.00128927 | 2.625589709 | 9.01E-11 | HDAC9 |
| ENSG00000271978 | 47.69751903 | 14.59459438 | 11.89772301 | 3.516850207 | 0.964219012 | 8.000491151 | 2.584623286 | 0.003215674 | AL359643.2 |
| ENSG00000186472 | 20.29681661 | 13.55212335 | 11.89772301 | 0.879212552 | 3.856876047 | 3.000184182 | 2.583889431 | 0.008619268 | PCLO |
| ENSG00000110042 | 42.62331488 | 44.82625417 | 46.50928087 | 7.912912967 | 6.749533083 | 8.000491151 | 2.562469825 | 6.77E-07 | DTX4 |
| ENSG00000167642 | 189.7752353 | 134.4787625 | 171.9761781 | 18.46346359 | 35.67610344 | 31.00190321 | 2.5518225 | 2.28E-19 | SPINT2 |
| ENSG00000135678 | 224.2798235 | 124.0540522 | 155.7520104 | 34.28928952 | 21.21281826 | 31.00190321 | 2.540386035 | 1.19E-17 | CPM |
| ENSG00000184524 | 146.1370796 | 158.4555961 | 191.4451794 | 28.13480166 | 29.89078937 | 30.00184182 | 2.494816789 | 1.79E-21 | CEND1 |
| ENSG00000105499 | 835.2140035 | 797.4903358 | 879.3498919 | 150.3453464 | 148.4897278 | 147.0090249 | 2.49370082 | 3.75E-105 | PLA2G4C |
| ENSG00000109072 | 28.41554325 | 26.06177568 | 34.61155786 | 7.033700415 | 3.856876047 | 5.00030697 | 2.477581862 | 0.000136476 | VTN |
| ENSG00000137809 | 32.47490658 | 12.50965233 | 17.30577893 | 2.637637656 | 6.749533083 | 2.000122788 | 2.456021918 | 0.003993506 | ITGA11 |
| ENSG00000023445 | 716.4776263 | 678.6486387 | 706.2921026 | 145.9492836 | 133.0622236 | 108.0066305 | 2.436548076 | 6.68E-80 | BIRC3 |
| ENSG00000172575 | 20.29681661 | 9.382239244 | 20.55061248 | 5.275275311 | 1.928438024 | 2.000122788 | 2.427978079 | 0.009392124 | RASGRP1 |
| ENSG00000145911 | 397.8176055 | 333.5907287 | 359.0949128 | 70.33700415 | 74.24486391 | 58.00356085 | 2.426407813 | 2.58E-43 | N4BP3 |
| ENSG00000146216 | 20.29681661 | 23.97683362 | 16.22416775 | 4.396062759 | 5.785314071 | 1.000061394 | 2.425373413 | 0.00343312 | TTBK1 |
| ENSG00000013293 | 31.46006574 | 33.35907287 | 29.20350194 | 4.396062759 | 5.785314071 | 8.000491151 | 2.380672204 | 0.000119613 | SLC7A14 |
| ENSG00000272636 | 75.09822146 | 84.4401532 | 65.97828217 | 12.30897573 | 25.06969431 | 6.000368364 | 2.376798013 | 4.25E-08 | DOC2B |
| ENSG00000105605 | 197.8939619 | 176.1776036 | 187.1187347 | 29.89322676 | 35.67610344 | 43.00263994 | 2.375873612 | 4.78E-23 | CACNG7 |
| ENSG00000108551 | 78.14274395 | 81.31274012 | 115.7323966 | 23.7387389 | 14.46328518 | 15.00092091 | 2.360858958 | 1.01E-10 | RASD1 |
| ENSG00000162772 | 453.6338512 | 526.4478687 | 427.2364173 | 95.83416815 | 103.1714343 | 77.00472733 | 2.348112313 | 3.91E-49 | ATF3 |
| ENSG00000106069 | 13.1929308 | 15.63706541 | 23.79544603 | 4.396062759 | 0.964219012 | 5.00030697 | 2.340603669 | 0.009095381 | CHN2 |
| ENSG00000163734 | 275.0218651 | 237.6833942 | 232.5464044 | 49.2359029 | 44.35407454 | 55.00337667 | 2.327993124 | 2.43E-29 | CXCL3 |
| ENSG00000110900 | 281.11091 | 404.4787585 | 328.8097997 | 82.64597988 | 58.81735972 | 62.00380642 | 2.312708786 | 1.97E-31 | TSPAN11 |
| ENSG00000198889 | 64.94981315 | 57.33590649 | 62.73344862 | 17.58425104 | 5.785314071 | 14.00085951 | 2.299785366 | 2.42E-07 | DCAF12L1 |
| ENSG00000137501 | 20.29681661 | 14.59459438 | 16.22416775 | 3.516850207 | 1.928438024 | 5.00030697 | 2.295428939 | 0.008810091 | SYTL2 |
| ENSG00000028277 | 199.9236436 | 203.2818503 | 158.9968439 | 45.7190527 | 36.64032245 | 32.00196461 | 2.292347651 | 7.03E-21 | POU2F2 |
| ENSG00000114654 | 42.62331488 | 42.74131211 | 42.18283614 | 6.154487863 | 15.42750419 | 5.00030697 | 2.265447167 | 3.21E-05 | EFCC1 |
| ENSG00000182752 | 140.0480346 | 116.756755 | 84.36567228 | 29.01401421 | 21.21281826 | 21.00128927 | 2.255674183 | 4.34E-12 | PAPPA |
| ENSG00000185101 | 51.75688235 | 39.61389903 | 61.65183744 | 12.30897573 | 8.677971106 | 11.00067533 | 2.254188968 | 2.05E-06 | ANO9 |
| ENSG00000259081 | 26.38586159 | 17.72200746 | 19.4690013 | 2.637637656 | 4.821095059 | 6.000368364 | 2.253704306 | 0.003528538 | AF111169.3 |
| ENSG00000227507 | 107.573128 | 102.1621607 | 77.87600518 | 17.58425104 | 15.42750419 | 28.00171903 | 2.243982133 | 1.23E-10 | LTB |
| ENSG00000198691 | 83.2169481 | 53.16602238 | 62.73344862 | 20.22188869 | 5.785314071 | 16.0009823 | 2.237889149 | 6.80E-07 | ABCA4 |
| ENSG00000267551 | 26.38586159 | 22.9343626 | 17.30577893 | 3.516850207 | 8.677971106 | 2.000122788 | 2.232532075 | 0.004203493 | AC005264.1 |
| ENSG00000160013 | 67.99433564 | 75.05791395 | 86.52889465 | 14.06740083 | 14.46328518 | 21.00128927 | 2.217669012 | 3.94E-09 | PTGIR |
| ENSG00000188338 | 32.47490658 | 39.61389903 | 37.85639141 | 5.275275311 | 11.57062814 | 7.000429757 | 2.211174281 | 9.30E-05 | SLC38A3 |
| ENSG00000081041 | 1336.545374 | 1052.895737 | 1470.991209 | 272.5558911 | 273.8381994 | 292.017927 | 2.203426059 | 3.69E-82 | CXCL2 |
| ENSG00000144339 | 43.63815571 | 35.44401492 | 31.36672431 | 0.879212552 | 15.42750419 | 8.000491151 | 2.200736121 | 0.000859186 | TMEFF2 |
| ENSG00000065609 | 27.40070242 | 22.9343626 | 18.38739011 | 6.154487863 | 6.749533083 | 2.000122788 | 2.195616565 | 0.003282623 | SNAP91 |
| ENSG00000118407 | 105.5434464 | 120.9266391 | 105.9978959 | 25.497164 | 21.21281826 | 26.00159624 | 2.192452406 | 4.66E-13 | FILIP1 |
| ENSG00000163751 | 52.77172318 | 56.29343547 | 67.05989335 | 15.82582593 | 12.53484715 | 10.00061394 | 2.190676721 | 3.82E-07 | CPA3 |
| ENSG00000100156 | 91.33567474 | 112.5868709 | 75.71278282 | 21.10110124 | 14.46328518 | 26.00159624 | 2.185556614 | 5.28E-10 | SLC16A8 |
| ENSG00000236609 | 27.40070242 | 19.80694952 | 33.52994668 | 7.033700415 | 1.928438024 | 9.000552545 | 2.167991779 | 0.001997201 | ZNF853 |
| ENSG00000128298 | 55.81624568 | 26.06177568 | 55.16217034 | 7.033700415 | 7.713752095 | 16.0009823 | 2.166437283 | 7.62E-05 | BAIAP2L2 |
| ENSG00000008517 | 2167.700014 | 2272.586839 | 2281.117985 | 503.7887922 | 536.1057706 | 509.0312495 | 2.117738052 | 1.05E-205 | IL32 |
| ENSG00000174672 | 38.56395156 | 27.10424671 | 37.85639141 | 4.396062759 | 8.677971106 | 11.00067533 | 2.117478469 | 0.000316511 | BRSK2 |
| ENSG00000118946 | 167.448737 | 132.3938204 | 154.6703992 | 32.53086442 | 42.42563652 | 31.00190321 | 2.101671587 | 9.78E-16 | PCDH17 |
| ENSG00000124749 | 11.16324914 | 26.06177568 | 34.61155786 | 8.792125519 | 4.821095059 | 3.000184182 | 2.094709238 | 0.006969495 | COL21A1 |
| ENSG00000011201 | 31.46006574 | 31.27413081 | 25.95866839 | 7.033700415 | 5.785314071 | 8.000491151 | 2.092613243 | 0.000664449 | ANOS1 |
| ENSG00000128342 | 27.40070242 | 36.48648595 | 29.20350194 | 5.275275311 | 8.677971106 | 8.000491151 | 2.091248957 | 0.000545343 | LIF |
| ENSG00000137872 | 349.1052457 | 333.5907287 | 324.4833549 | 81.76676732 | 74.24486391 | 82.0050343 | 2.08103112 | 1.85E-35 | SEMA6D |
| ENSG00000131386 | 377.5207889 | 492.0463248 | 343.9523562 | 96.71338071 | 110.8851864 | 80.00491151 | 2.076030059 | 9.14E-31 | GALNT15 |
| ENSG00000124875 | 581.5037959 | 467.0270202 | 528.9078685 | 131.0026702 | 131.1337856 | 119.0073059 | 2.048284854 | 2.67E-47 | CXCL6 |
| ENSG00000146411 | 157.3003287 | 141.7760597 | 160.0784551 | 36.04771463 | 38.56876047 | 37.00227158 | 2.040950837 | 2.63E-16 | SLC2A12 |
| ENSG00000158050 | 19.28197578 | 19.80694952 | 22.71383485 | 6.154487863 | 1.928438024 | 7.000429757 | 2.032673456 | 0.008990135 | DUSP2 |
| ENSG00000204381 | 128.8847855 | 112.5868709 | 110.3243407 | 35.16850207 | 26.99813233 | 24.00147345 | 2.023770656 | 5.14E-12 | LAYN |
| ENSG00000113361 | 92.35051557 | 77.14285601 | 86.52889465 | 20.22188869 | 22.17703727 | 21.00128927 | 2.014792234 | 2.78E-09 | CDH6 |
| ENSG00000253522 | 154.2558062 | 136.5637046 | 173.0577893 | 38.68535228 | 44.35407454 | 32.00196461 | 2.00959507 | 3.81E-15 | MIR3142HG |
| ENSG00000214900 | 25.37102076 | 22.9343626 | 22.71383485 | 9.671338071 | 2.892657035 | 5.00030697 | 2.000900782 | 0.005555529 | LINC01588 |
| ENSG00000182308 | 24.35617993 | 27.10424671 | 16.22416775 | 6.154487863 | 7.713752095 | 3.000184182 | 1.999776451 | 0.006775566 | DCAF4L1 |
| ENSG00000035664 | 52.77172318 | 64.63320368 | 42.18283614 | 15.82582593 | 7.713752095 | 17.0010437 | 1.976214409 | 2.16E-05 | DAPK2 |
| ENSG00000228594 | 223.2649827 | 241.8532783 | 197.9348465 | 61.54487863 | 60.74579774 | 46.00282412 | 1.975090527 | 1.91E-20 | FNDC10 |
| ENSG00000176697 | 909.2973841 | 845.444003 | 868.53378 | 242.6626643 | 223.6988107 | 203.012463 | 1.968163353 | 4.17E-76 | BDNF |
| ENSG00000165092 | 6501.07036 | 6719.768241 | 6350.139256 | 1684.571249 | 1672.919986 | 1661.101975 | 1.963268223 | 0 | ALDH1A1 |
| ENSG00000115009 | 63.93497232 | 43.78378314 | 46.50928087 | 13.18818828 | 16.3917232 | 10.00061394 | 1.961176635 | 2.00E-05 | CCL20 |
| ENSG00000130755 | 93.3653564 | 88.61003731 | 71.38633808 | 7.033700415 | 31.81922739 | 27.00165764 | 1.95650077 | 3.32E-06 | GMFG |
| ENSG00000164056 | 213.1165744 | 187.6447849 | 153.588788 | 42.20220249 | 44.35407454 | 58.00356085 | 1.944013225 | 5.62E-16 | SPRY1 |
| ENSG00000092969 | 1081.820325 | 1018.494194 | 966.9603977 | 236.5081765 | 256.4822571 | 306.0187865 | 1.943877718 | 1.16E-74 | TGFB2 |
| ENSG00000021300 | 52.77172318 | 51.08108033 | 36.77478023 | 14.06740083 | 16.3917232 | 6.000368364 | 1.941449792 | 0.000105105 | PLEKHB1 |
| ENSG00000170214 | 36.5342699 | 43.78378314 | 27.04027958 | 9.671338071 | 11.57062814 | 7.000429757 | 1.924753067 | 0.000565066 | ADRA1B |
| ENSG00000065320 | 22.32649827 | 37.52895698 | 41.10122496 | 7.033700415 | 8.677971106 | 11.00067533 | 1.922758332 | 0.001035138 | NTN1 |
| ENSG00000110328 | 21.31165744 | 20.84942054 | 29.20350194 | 7.033700415 | 6.749533083 | 5.00030697 | 1.920132789 | 0.005483917 | GALNT18 |
| ENSG00000243742 | 62.92013149 | 72.9729719 | 85.44728347 | 20.22188869 | 10.60640913 | 28.00171903 | 1.913428798 | 1.55E-06 | RPLP0P2 |
| ENSG00000082497 | 51.75688235 | 46.91119622 | 67.05989335 | 10.55055062 | 23.14125628 | 11.00067533 | 1.893448136 | 3.87E-05 | SERTAD4 |
| ENSG00000118473 | 587.5928408 | 519.1505715 | 558.1113705 | 156.4998342 | 133.0622236 | 160.009823 | 1.888558255 | 2.45E-46 | SGIP1 |
| ENSG00000257591 | 38.56395156 | 23.97683362 | 33.52994668 | 1.758425104 | 12.53484715 | 12.00073673 | 1.886376605 | 0.004582506 | ZNF625 |
| ENSG00000140873 | 533.8062768 | 454.5173678 | 465.0928087 | 125.7273949 | 134.9906617 | 133.0081654 | 1.885514837 | 3.96E-41 | ADAMTS18 |
| ENSG00000072041 | 173.537782 | 161.5830092 | 184.9555123 | 39.56456483 | 53.03204565 | 49.0030083 | 1.880325122 | 8.27E-16 | SLC6A15 |
| ENSG00000087085 | 47.69751903 | 29.18918876 | 52.99894797 | 6.154487863 | 12.53484715 | 17.0010437 | 1.875388899 | 0.000454387 | ACHE |
| ENSG00000184368 | 76.11306229 | 70.88802985 | 61.65183744 | 26.37637656 | 20.24859925 | 10.00061394 | 1.871145481 | 3.69E-06 | MAP7D2 |
| ENSG00000179256 | 27.40070242 | 17.72200746 | 20.55061248 | 6.154487863 | 4.821095059 | 7.000429757 | 1.871029054 | 0.009782465 | SMCO3 |
| ENSG00000137474 | 163.3893737 | 219.9613867 | 190.3635682 | 50.11511546 | 63.63845478 | 43.00263994 | 1.870925648 | 1.54E-15 | MYO7A |
| ENSG00000128645 | 134.9738305 | 135.5212335 | 168.7313446 | 27.25558911 | 50.13938861 | 45.00276273 | 1.849524209 | 1.60E-11 | HOXD1 |
| ENSG00000135914 | 105.5434464 | 99.03474758 | 101.6714512 | 25.497164 | 38.56876047 | 21.00128927 | 1.848084096 | 5.31E-09 | HTR2B |
| ENSG00000108679 | 35.51942907 | 20.84942054 | 18.38739011 | 5.275275311 | 9.642190118 | 6.000368364 | 1.843273104 | 0.008207129 | LGALS3BP |
| ENSG00000270607 | 426.2331488 | 394.0540483 | 514.8469232 | 108.1431439 | 134.9906617 | 130.0079812 | 1.841199926 | 1.11E-31 | AC009549.1 |
| ENSG00000147408 | 300.3928858 | 325.2509605 | 296.3614642 | 81.76676732 | 100.2787772 | 76.00466594 | 1.837253804 | 1.05E-25 | CSGALNACT1 |
| ENSG00000100767 | 903.2083391 | 746.4092554 | 867.4521688 | 246.1795145 | 217.9134967 | 241.0147959 | 1.835248813 | 2.73E-59 | PAPLN |
| ENSG00000155966 | 58.86076817 | 44.82625417 | 43.26444732 | 16.70503849 | 14.46328518 | 10.00061394 | 1.830065128 | 7.81E-05 | AFF2 |

| | | | | | | | | | |
|---|---|---|---|---|---|---|---|---|---|
| ENSG00000112149 | 104.5286055 | 114.671813 | 89.7737282 | 22.85952635 | 34.71188443 | 30.00184182 | 1.824003839 | 3.98E-09 | CD83 |
| ENSG00000270547 | 452.6190104 | 382.586867 | 370.9926358 | 121.3313322 | 87.74393008 | 132.008104 | 1.822704107 | 2.88E-27 | LINC01235 |
| ENSG00000162595 | 483.0642353 | 437.8378314 | 405.6041937 | 123.0897573 | 115.7062814 | 138.0084724 | 1.817388359 | 5.92E-35 | DIRAS3 |
| ENSG00000074527 | 7517.940872 | 7601.69873 | 7014.248522 | 2003.725406 | 2195.52669 | 2100.128927 | 1.813574691 | 0 | NTN4 |
| ENSG00000188112 | 84.23178893 | 55.25096444 | 71.38633808 | 24.61795145 | 18.32016122 | 17.0010437 | 1.809506266 | 2.62E-06 | C6orf132 |
| ENSG00000171004 | 29.43038408 | 35.44401492 | 23.79544603 | 11.42976317 | 7.713752095 | 6.000368364 | 1.809247817 | 0.00327557 | HS6ST2 |
| ENSG00000144218 | 122.7957405 | 151.1582989 | 123.3036749 | 35.16850207 | 29.89078937 | 49.0030083 | 1.803932424 | 8.35E-11 | AFF3 |
| ENSG00000175264 | 200.9384844 | 193.899611 | 246.6073497 | 73.85385436 | 49.1751696 | 61.00374503 | 1.796977891 | 3.64E-16 | CHST1 |
| ENSG00000254585 | 22.32649827 | 43.78378314 | 31.36672431 | 12.30897573 | 4.821095059 | 11.00067533 | 1.787704786 | 0.003580026 | MAGEL2 |
| ENSG00000170891 | 276.0367059 | 272.0849381 | 261.7499063 | 87.04204264 | 55.92470269 | 92.00564824 | 1.784618398 | 1.71E-19 | CYTL1 |
| ENSG00000198046 | 37.54911073 | 26.06177568 | 31.36672431 | 6.154487863 | 8.677971106 | 13.00079812 | 1.781329941 | 0.002671326 | ZNF667 |
| ENSG00000149633 | 48.71235986 | 22.9343626 | 28.12189076 | 7.033700415 | 11.57062814 | 11.00067533 | 1.760710596 | 0.003314827 | KIAA1755 |
| ENSG00000143369 | 297.3483633 | 283.5521194 | 317.9936878 | 91.43810539 | 86.77971106 | 87.00534127 | 1.759857269 | 5.24E-25 | ECM1 |
| ENSG00000125740 | 23.3413391 | 31.27413081 | 44.34605851 | 9.671338071 | 11.57062814 | 8.000491151 | 1.755491765 | 0.002642018 | FOSB |
| ENSG00000013016 | 144.1073979 | 120.9266391 | 144.9358985 | 34.28928952 | 40.4971985 | 47.00288551 | 1.755377543 | 2.25E-11 | EHD3 |
| ENSG00000100311 | 2192.056194 | 2078.687228 | 2179.446534 | 664.6846892 | 642.1698619 | 603.0370205 | 1.754731717 | 7.02E-145 | PDGFB |
| ENSG00000135063 | 405.9363322 | 332.5482577 | 349.3604141 | 120.4521196 | 82.92283502 | 119.0073059 | 1.753696488 | 5.23E-24 | FAM189A2 |
| ENSG00000081842 | 35.51942907 | 17.72200746 | 25.95866839 | 9.671338071 | 6.749533083 | 7.000429757 | 1.75314371 | 0.008182844 | PCDHA6 |
| ENSG00000129910 | 23.3413391 | 25.01930465 | 33.52994668 | 11.42976317 | 6.749533083 | 6.000368364 | 1.748227379 | 0.00654233 | CDH15 |
| ENSG00000267534 | 19.28197578 | 43.78378314 | 38.93800259 | 7.033700415 | 16.3917232 | 7.000429757 | 1.746991977 | 0.004932987 | S1PR2 |
| ENSG00000174175 | 55.81624568 | 72.9729719 | 59.4886507 | 24.61795145 | 12.53484715 | 19.00116648 | 1.739381753 | 2.09E-05 | SELP |
| ENSG00000188372 | 25.37102076 | 26.06177568 | 23.79544603 | 6.154487863 | 10.60640913 | 6.000368364 | 1.727103486 | 0.009027873 | ZP3 |
| ENSG00000168386 | 1648.101509 | 1618.957505 | 1443.950929 | 465.1034399 | 510.0718573 | 453.0278114 | 1.722213142 | 8.36E-94 | FILIP1L |
| ENSG00000182107 | 32.47490658 | 34.4015439 | 25.95866839 | 13.18818828 | 4.821095059 | 10.00061394 | 1.722039754 | 0.004560227 | TMEM30B |
| ENSG00000147676 | 51.75688235 | 56.29343547 | 60.57022625 | 12.30897573 | 18.32016122 | 21.00128927 | 1.71442744 | 4.65E-05 | MAL2 |
| ENSG00000128872 | 308.5116125 | 236.6409232 | 249.8521833 | 74.73306691 | 77.13752095 | 91.00558685 | 1.713617789 | 6.02E-19 | TMOD2 |
| ENSG00000163131 | 230.3688685 | 177.2200746 | 228.2199596 | 61.54487863 | 70.38798786 | 62.00380642 | 1.713547887 | 6.71E-16 | CTSS |
| ENSG00000159784 | 106.5582872 | 90.69497936 | 89.7737282 | 31.65165187 | 31.81922739 | 24.00147345 | 1.711728508 | 4.31E-08 | FAM131B |
| ENSG00000176907 | 780.4125986 | 709.9227695 | 758.2094394 | 222.4407756 | 242.018972 | 223.0136908 | 1.710033641 | 1.63E-55 | TCIM |
| ENSG00000172738 | 234.4282318 | 250.1930465 | 264.9947399 | 76.49149201 | 77.13752095 | 76.00466594 | 1.706359656 | 3.40E-20 | TMEM217 |
| ENSG00000196562 | 3609.788834 | 3616.331993 | 3342.178556 | 1107.807815 | 1136.814215 | 1011.062069 | 1.698223517 | 4.66E-174 | SULF2 |
| ENSG00000277117 | 113.662173 | 165.7528933 | 102.7530624 | 59.78645353 | 27.96235134 | 30.00184182 | 1.690100366 | 3.02E-07 | FP565260.3 |
| ENSG00000164099 | 66.97949481 | 77.14285601 | 87.61050583 | 32.53086442 | 19.28438024 | 20.00122788 | 1.681342267 | 3.89E-06 | PRSS12 |
| ENSG00000123095 | 26.38586159 | 41.69884109 | 35.69316904 | 7.033700415 | 6.749533083 | 19.00116648 | 1.673212392 | 0.005351634 | BHLHE41 |
| ENSG00000078401 | 12725.08917 | 12474.20831 | 12927.41686 | 3869.414441 | 4189.531606 | 3936.241646 | 1.668551767 | 0 | EDN1 |
| ENSG00000158406 | 55.81624568 | 41.69884109 | 28.12189076 | 12.30897573 | 17.35594221 | 10.00061394 | 1.664774553 | 0.001399332 | HIST1H4H |
| ENSG00000273264 | 28.41554325 | 37.52895698 | 27.04027958 | 7.912912967 | 10.60640913 | 11.00067533 | 1.660461517 | 0.004333402 | AL360219.1 |
| ENSG00000165072 | 239.502436 | 221.0038578 | 208.7509583 | 65.06172884 | 76.17330193 | 71.00435897 | 1.658931105 | 4.32E-17 | MAMDC2 |
| ENSG00000184922 | 58.86076817 | 56.29343547 | 62.73344862 | 23.7387389 | 12.53484715 | 20.00122788 | 1.65555865 | 5.63E-05 | FMNL1 |
| ENSG00000064692 | 183.6861903 | 183.4749008 | 182.7922899 | 62.42409118 | 55.92470269 | 56.00343806 | 1.655530482 | 1.30E-14 | SNCAIP |
| ENSG00000165449 | 66.97949481 | 93.82239244 | 57.3253927 | 18.46346359 | 24.1054753 | 27.00165764 | 1.654084729 | 1.33E-05 | SLC16A9 |
| ENSG00000148200 | 32.47490658 | 32.31660184 | 24.87705721 | 7.033700415 | 8.677971106 | 13.00079812 | 1.651910388 | 0.005894215 | NR6A1 |
| ENSG00000268355 | 42.62331488 | 43.78378314 | 35.69316904 | 8.792125519 | 21.21281826 | 9.000552545 | 1.650500819 | 0.001861765 | AC243960.3 |
| ENSG00000196611 | 27.40070242 | 47.95366725 | 32.44833549 | 7.033700415 | 10.60640913 | 17.0010437 | 1.648069243 | 0.003961425 | MMP1 |
| ENSG00000272720 | 61.90529066 | 60.46331957 | 69.22311572 | 18.46346359 | 26.03391332 | 17.0010437 | 1.639611367 | 2.36E-05 | AL022322.1 |
| ENSG00000228495 | 1608.522716 | 1632.509628 | 1601.866162 | 608.4150859 | 464.7535637 | 479.0294077 | 1.638981178 | 1.13E-72 | LINC01013 |
| ENSG00000154654 | 66.97949481 | 66.71814574 | 76.794394 | 29.89322676 | 18.32016122 | 19.00116648 | 1.638929567 | 1.23E-05 | NCAM2 |
| ENSG00000163545 | 237.4727543 | 272.0849381 | 290.9534083 | 81.76676732 | 84.85127304 | 91.00558685 | 1.636376972 | 3.21E-19 | NUAK2 |
| ENSG00000127561 | 90.32083391 | 77.14285601 | 82.20244992 | 21.9803138 | 32.7834464 | 26.00159624 | 1.631789084 | 1.16E-06 | SYNGR3 |
| ENSG00000140945 | 1620.700806 | 1651.274107 | 1602.947773 | 545.9909947 | 547.6763987 | 484.0297147 | 1.626413461 | 3.05E-100 | CDH13 |
| ENSG00000085465 | 45.66783737 | 38.571428 | 35.69316904 | 7.912912967 | 16.3917232 | 15.00092091 | 1.618859192 | 0.001650179 | OVGP1 |
| ENSG00000167971 | 62.92013149 | 51.08108033 | 46.50928087 | 20.22188869 | 24.1054753 | 8.000491151 | 1.611474205 | 0.000549567 | CASKIN1 |
| ENSG00000255836 | 66.97949481 | 56.29343547 | 40.01961377 | 19.34267614 | 16.3917232 | 18.00110509 | 1.60367591 | 0.000206633 | AC131206.1 |
| ENSG00000235505 | 639.3497232 | 622.3552032 | 698.7208243 | 207.4941622 | 219.8419347 | 223.0136908 | 1.59256209 | 1.62E-43 | CASP17P |
| ENSG00000175874 | 34.50458824 | 40.65637006 | 35.69316904 | 14.06740083 | 7.713752095 | 15.00092091 | 1.590413484 | 0.002699883 | CREG2 |
| ENSG00000130540 | 37.54911073 | 51.08108033 | 63.8150598 | 15.82582593 | 26.03391332 | 9.000552545 | 1.580272419 | 0.001244646 | SULT4A1 |
| ENSG00000152689 | 535.8359585 | 457.6447809 | 432.6444732 | 170.5672351 | 149.4539468 | 157.0096388 | 1.579235445 | 5.17E-29 | RASGRP3 |
| ENSG00000154678 | 176.5823045 | 138.6486466 | 146.0175097 | 49.2359029 | 53.03204565 | 53.00325388 | 1.572460238 | 6.21E-11 | PDE1C |
| ENSG00000176463 | 67.99433564 | 72.9729719 | 110.3243407 | 31.65165187 | 30.85500838 | 22.00135067 | 1.567022898 | 1.04E-05 | SLCO3A1 |
| ENSG00000058085 | 1207.660588 | 1005.984541 | 1173.548134 | 412.3506868 | 372.1885386 | 370.0227158 | 1.551506437 | 1.04E-56 | LAMC2 |
| ENSG00000187372 | 66.97949481 | 86.52509525 | 72.46794927 | 25.497164 | 26.03391332 | 26.00159624 | 1.543408997 | 8.77E-06 | PCDHB13 |
| ENSG00000189134 | 187.7455536 | 209.5366765 | 150.3439545 | 63.30330373 | 71.35220687 | 54.00331527 | 1.537013131 | 2.37E-11 | NKAPL |
| ENSG00000130203 | 178.6119862 | 180.3474877 | 163.3232886 | 58.90724098 | 65.5668928 | 56.00343806 | 1.533208813 | 2.91E-12 | APOE |
| ENSG00000184292 | 1512.112837 | 1433.397662 | 1515.337268 | 543.3533571 | 512.9645143 | 504.0309425 | 1.514455113 | 4.03E-86 | TACSTD2 |
| ENSG00000043591 | 234.4282318 | 209.5366765 | 268.2395734 | 84.40440498 | 78.10173996 | 87.00534127 | 1.512729572 | 4.65E-15 | ADRB1 |
| ENSG00000185133 | 49.72720069 | 45.86872519 | 57.3253927 | 17.58425104 | 23.14125628 | 13.00079812 | 1.506830171 | 0.000622222 | INPP5J |
| ENSG00000284237 | 36.5342699 | 34.4015439 | 32.44833549 | 16.70503849 | 14.46328518 | 5.00030697 | 1.504826486 | 0.009255942 | AL356275.1 |
| ENSG00000242615 | 43.63815571 | 27.10424671 | 33.52994668 | 13.18818828 | 9.642190118 | 14.00085951 | 1.50188218 | 0.006158791 | AC022415.1 |
| ENSG00000142871 | 23765.54257 | 23713.08845 | 25279.41657 | 8289.215939 | 8731.003152 | 8700.534127 | 1.500318984 | 0 | CYR61 |
| ENSG00000257800 | 32.47490658 | 29.18918876 | 56.24378152 | 13.18818828 | 13.49906617 | 15.00092091 | 1.499351523 | 0.00452422 | FNBP1P1 |
| ENSG00000147113 | 3911.196561 | 3702.857088 | 3722.905692 | 1317.060403 | 1293.981914 | 1406.08632 | 1.497398579 | 7.66E-168 | CXorf36 |
| ENSG00000204389 | 6953.68937 | 7974.903358 | 8000.677921 | 2391.458141 | 2830.947019 | 2912.178779 | 1.495285924 | 1.43E-42 | HSPA1A |
| ENSG00000250474 | 42.62331488 | 45.86872519 | 30.28511313 | 6.154487863 | 16.3917232 | 20.00122788 | 1.494117176 | 0.006839327 | WBP1LP2 |
| ENSG00000186642 | 1300.011104 | 1238.45558 | 1230.873526 | 430.8141504 | 490.787477 | 419.0257241 | 1.491722489 | 1.44E-66 | PDE2A |
| ENSG00000196169 | 137.0035121 | 169.9227774 | 154.6703992 | 59.78645353 | 49.1751696 | 55.00337667 | 1.491232206 | 3.78E-10 | KIF19 |
| ENSG00000111981 | 166.4338962 | 203.2818503 | 164.4048998 | 49.2359029 | 69.42376885 | 72.00442036 | 1.490948884 | 1.31E-10 | ULBP1 |
| ENSG00000109743 | 405.9363322 | 393.0115772 | 442.3789739 | 153.8621966 | 138.8475377 | 150.0092091 | 1.486441319 | 8.81E-26 | BST1 |
| ENSG00000204403 | 41.60847405 | 40.65637006 | 54.08055916 | 11.42976317 | 10.60640913 | 27.00165764 | 1.483683287 | 0.003546429 | CASP12 |
| ENSG00000116774 | 27.40070242 | 43.78378314 | 33.52994668 | 14.06740083 | 13.49906617 | 10.00061394 | 1.474677893 | 0.006876865 | OLFML3 |
| ENSG00000125378 | 3753.896232 | 3781.042415 | 3786.720752 | 1308.268277 | 1372.083654 | 1399.08589 | 1.473351287 | 7.51E-181 | BMP4 |
| ENSG00000068976 | 52.77172318 | 39.61389903 | 52.99894797 | 20.22188869 | 18.32016122 | 14.00085951 | 1.463820071 | 0.001083951 | PYGM |
| ENSG00000182013 | 241.5321177 | 223.0887998 | 192.5267906 | 73.85385436 | 75.20908292 | 91.00558685 | 1.455802074 | 4.98E-13 | PNMA8A |
| ENSG00000213626 | 229.3540277 | 247.0656334 | 211.9957919 | 70.33700415 | 89.6723681 | 92.00564824 | 1.453586107 | 1.10E-13 | LBH |
| ENSG00000165152 | 74.08338062 | 115.714284 | 103.8346736 | 44.83984015 | 25.06969431 | 37.00227158 | 1.452971233 | 1.19E-05 | TMEM246 |
| ENSG00000120324 | 48.71235986 | 36.48648595 | 33.52994668 | 14.94661338 | 14.46328518 | 14.00085951 | 1.451992191 | 0.003604284 | PCDHB10 |
| ENSG00000100228 | 181.6565087 | 129.2664074 | 137.3646203 | 53.63196566 | 58.81735972 | 52.00319248 | 1.447632586 | 6.55E-09 | RAB36 |
| ENSG00000179909 | 288.2147959 | 183.4749008 | 240.1176826 | 106.3847188 | 74.24486391 | 80.00491151 | 1.446328173 | 6.48E-11 | ZNF154 |
| ENSG00000207870 | 52.77172318 | 46.91119622 | 33.52994668 | 16.70503849 | 19.28438024 | 13.00079812 | 1.442770891 | 0.002504249 | MIR221 |
| ENSG00000243710 | 51.75688235 | 62.54826163 | 73.54956045 | 35.16850207 | 14.46328518 | 19.00116648 | 1.442177331 | 0.000764728 | CFAP57 |
| ENSG00000088899 | 276.0367059 | 264.7876409 | 319.075299 | 111.6599941 | 88.70814909 | 116.0071217 | 1.441720972 | 5.92E-16 | LZTS3 |
| ENSG00000160145 | 511.4797786 | 465.9845491 | 426.1548061 | 162.6543221 | 180.3089552 | 175.0107439 | 1.439848323 | 4.38E-26 | KALRN |
| ENSG00000178343 | 224.2798235 | 190.772198 | 219.5670702 | 66.82015394 | 86.77971106 | 81.00497291 | 1.438498256 | 1.40E-12 | SHISA3 |
| ENSG00000197822 | 282.1257509 | 265.8301119 | 224.9751261 | 93.1965305 | 96.42190118 | 96.00589382 | 1.437400068 | 3.13E-15 | OCLN |

| | | | | | | | | | |
|---|---|---|---|---|---|---|---|---|---|
| ENSG00000157240 | 321.7045433 | 364.8648595 | 329.8914108 | 118.6936945 | 128.2411286 | 129.0079198 | 1.436105209 | 2.26E-20 | FZD1 |
| ENSG00000184838 | 54.80140485 | 61.5057906 | 51.91733679 | 25.497164 | 10.60640913 | 26.00159624 | 1.435670582 | 0.001100015 | PRR16 |
| ENSG00000175746 | 544.969526 | 501.4285641 | 512.6837008 | 147.7077087 | 232.3767818 | 199.0122174 | 1.432223272 | 6.04E-22 | C15orf54 |
| ENSG00000117643 | 40.59363322 | 46.91119622 | 37.85639141 | 12.30897573 | 15.42750419 | 19.00116648 | 1.429317294 | 0.0030648 | MAN1C1 |
| ENSG00000137266 | 489.1532803 | 462.857136 | 555.9481481 | 153.8621966 | 213.0924016 | 194.0119104 | 1.429012555 | 3.15E-23 | SLC22A23 |
| ENSG00000164932 | 1698.84355 | 1682.548238 | 1823.596455 | 662.9262641 | 675.9175273 | 598.0367136 | 1.425126007 | 2.41E-80 | CTHRC1 |
| ENSG00000166257 | 198.9088028 | 194.9420821 | 168.7313446 | 58.02802842 | 71.35220687 | 81.00497291 | 1.423639024 | 6.56E-11 | SCN3B |
| ENSG00000233822 | 66.97949481 | 70.88802985 | 57.3253927 | 25.497164 | 17.35594221 | 30.00184182 | 1.423142372 | 0.000212724 | HIST1H2BN |
| ENSG00000108387 | 63.93497232 | 50.0386093 | 36.77478023 | 21.9803138 | 24.1054753 | 10.00061394 | 1.422228466 | 0.003098747 | SEPTIN4 |
| ENSG00000183049 | 90.32083391 | 94.86486347 | 59.48861507 | 43.08141504 | 26.03391332 | 22.00135067 | 1.418159588 | 0.000146323 | CAMK1D |
| ENSG00000156345 | 46.6826782 | 50.0386093 | 36.77478023 | 16.70503849 | 18.32016122 | 15.00092091 | 1.416129657 | 0.002238864 | CDK20 |
| ENSG00000168140 | 39.57879239 | 53.16602238 | 34.61155786 | 21.10110124 | 12.53484715 | 14.00085951 | 1.412640085 | 0.004380007 | VASN |
| ENSG00000136367 | 104.5286055 | 70.88802985 | 80.03922755 | 35.16850207 | 35.67610344 | 25.00153485 | 1.411899162 | 2.99E-05 | ZFHX2 |
| ENSG00000148848 | 52.77172318 | 59.42084855 | 31.36672431 | 12.30897573 | 18.32016122 | 24.00147345 | 1.402666321 | 0.003917335 | ADAM12 |
| ENSG00000138311 | 37.54911073 | 34.4015439 | 46.50928087 | 8.792125519 | 21.21281826 | 15.00092091 | 1.402530147 | 0.007588424 | ZNF365 |
| ENSG00000146374 | 110.6176505 | 75.05791395 | 117.895619 | 29.01401421 | 40.4971985 | 46.00282412 | 1.398786548 | 1.30E-05 | RSPO3 |
| ENSG00000204642 | 74.08338062 | 66.71814574 | 74.63117163 | 33.41007697 | 19.28438024 | 29.00178042 | 1.395275982 | 0.000120751 | HLA-F |
| ENSG00000135919 | 166.4338962 | 215.7915026 | 164.4048998 | 81.76676732 | 64.60267379 | 61.00374503 | 1.3947315 | 9.93E-10 | SERPINE2 |
| ENSG00000204388 | 2753.263173 | 3099.266364 | 3093.407984 | 945.1534933 | 1212.023298 | 1252.076865 | 1.392320235 | 2.78E-24 | HSPA1B |
| ENSG00000198342 | 96.4098789 | 71.93050087 | 49.75411442 | 24.61795145 | 26.03391332 | 33.002026 | 1.387371731 | 0.000311401 | ZNF442 |
| ENSG00000277639 | 61.90529066 | 50.0386093 | 57.3253927 | 25.497164 | 26.99813233 | 12.00073673 | 1.386179102 | 0.001311355 | AC007906.2 |
| ENSG00000169604 | 216.1610969 | 189.7297269 | 154.6703992 | 53.63196566 | 81.958616 | 80.00491151 | 1.383935476 | 4.30E-09 | ANTXR1 |
| ENSG00000100344 | 47.69751903 | 60.46331957 | 70.3047269 | 24.61795145 | 22.17703727 | 22.00135067 | 1.372302625 | 0.000547488 | PNPLA3 |
| ENSG00000163739 | 5228.459959 | 4508.687192 | 5355.056968 | 1921.079426 | 1912.0463 | 2029.124568 | 1.36447115 | 4.79E-110 | CXCL1 |
| ENSG00000047617 | 229.3540277 | 241.8532783 | 244.4441274 | 86.16283008 | 85.81549205 | 107.0065692 | 1.361224897 | 2.62E-13 | ANO2 |
| ENSG00000229689 | 37.54911073 | 35.44401492 | 56.24378152 | 24.61795145 | 9.642190118 | 16.0009823 | 1.353425314 | 0.009204518 | AC009237.3 |
| ENSG00000167103 | 56.83108651 | 32.31660184 | 36.77478023 | 14.06740083 | 11.57062814 | 24.00147345 | 1.34963884 | 0.009683084 | PIP5KL1 |
| ENSG00000060656 | 255.7398893 | 271.0424671 | 297.4430754 | 118.6936945 | 83.88705403 | 121.0074287 | 1.347629638 | 3.41E-13 | PTPRU |
| ENSG00000184545 | 740.8338062 | 770.3860891 | 829.5957774 | 305.9659681 | 301.8005507 | 317.0194619 | 1.3397604 | 3.29E-39 | DUSP8 |
| ENSG00000169429 | 4844.850125 | 4116.718086 | 4712.579925 | 1772.492505 | 1794.411581 | 1841.113026 | 1.33851562 | 3.26E-110 | CXCL8 |
| ENSG00000152128 | 120.7660588 | 139.6911176 | 82.20244992 | 44.83984015 | 49.1751696 | 42.00257854 | 1.333810254 | 6.87E-06 | TMEM163 |
| ENSG00000269388 | 43.63815571 | 57.33590649 | 27.04027958 | 17.58425104 | 14.46328518 | 19.00116648 | 1.327999746 | 0.008692403 | AC018755.3 |
| ENSG00000254837 | 49.72720069 | 56.29343547 | 50.83572561 | 19.34267614 | 15.42750419 | 28.00171903 | 1.324928403 | 0.002092069 | AP001372.2 |
| ENSG00000189337 | 129.8996263 | 174.0926615 | 153.588788 | 54.51117822 | 68.45954984 | 60.00368364 | 1.323631779 | 4.16E-08 | KAZN |
| ENSG00000139926 | 3548.898384 | 3416.177556 | 3563.908848 | 1371.571581 | 1429.936795 | 1412.086688 | 1.321580986 | 2.35E-142 | FRMD6 |
| ENSG00000105696 | 59.875609 | 39.61389903 | 33.52994668 | 13.18818828 | 18.32016122 | 22.00135067 | 1.321474451 | 0.007339815 | TMEM59L |
| ENSG00000165801 | 289.2296367 | 236.6409232 | 218.485459 | 101.988656 | 97.38612019 | 100.0061394 | 1.314548838 | 2.67E-12 | ARHGEF40 |
| ENSG00000075275 | 532.791436 | 559.8069416 | 491.0514771 | 201.3396744 | 206.3428685 | 230.0141206 | 1.314018891 | 1.05E-25 | CELSR1 |
| ENSG00000169860 | 48.71235986 | 51.08108033 | 38.93800259 | 15.82582593 | 15.42750419 | 25.00153485 | 1.308094149 | 0.004690892 | P2RY1 |
| ENSG00000136274 | 261.8289343 | 271.0424671 | 307.177576 | 131.0026702 | 90.63658711 | 117.0071831 | 1.307954644 | 7.30E-13 | NACAD |
| ENSG00000149243 | 142.0777163 | 215.7915026 | 131.9565643 | 58.90724098 | 75.20908292 | 64.00392921 | 1.307541436 | 3.99E-07 | KLHL35 |
| ENSG00000118971 | 130.9144671 | 187.6447849 | 140.6094538 | 70.33700415 | 53.03204565 | 62.00380642 | 1.306372438 | 1.80E-07 | CCND2 |
| ENSG00000136732 | 51.75688235 | 81.31274012 | 72.46794927 | 24.61795145 | 25.06969431 | 34.00208739 | 1.298810862 | 0.000597254 | GYPC |
| ENSG00000164684 | 318.6600208 | 268.957525 | 261.7499063 | 122.2105447 | 91.60080612 | 132.008104 | 1.296899418 | 1.85E-12 | ZNF704 |
| ENSG00000171435 | 272.9921834 | 247.0656334 | 180.6290676 | 91.43810539 | 81.958616 | 114.0069989 | 1.288413584 | 1.88E-09 | KSR2 |
| ENSG00000205502 | 429.2776713 | 366.9498016 | 414.2570831 | 185.5138484 | 149.4539468 | 160.009823 | 1.288056657 | 1.74E-18 | C2CD4B |
| ENSG00000138080 | 104.5286055 | 95.9073345 | 88.69211701 | 32.53086442 | 42.42563652 | 44.00270133 | 1.285757837 | 1.83E-05 | EMILIN1 |
| ENSG00000166016 | 140.0480346 | 88.61003731 | 121.1404525 | 62.42409118 | 43.38985553 | 39.00239436 | 1.267273867 | 2.69E-05 | ABTB2 |
| ENSG00000129946 | 308.5116125 | 386.7567511 | 394.7880818 | 142.4324334 | 173.5594221 | 137.008411 | 1.266459075 | 6.88E-15 | SHC2 |
| ENSG00000234745 | 3764.04464 | 3924.903417 | 3771.578195 | 1555.327004 | 1579.390741 | 1633.100256 | 1.265713053 | 6.69E-139 | HLA-B |
| ENSG00000113555 | 693.1362872 | 598.3783696 | 617.5999856 | 277.8311664 | 243.94741 | 273.0167605 | 1.26401828 | 1.62E-27 | PCDH12 |
| ENSG00000184371 | 1938.345986 | 1896.254798 | 1955.553019 | 778.982321 | 872.6182057 | 761.0467208 | 1.263085788 | 1.27E-72 | CSF1 |
| ENSG00000008300 | 174.5526228 | 159.4980672 | 170.8945669 | 58.02802842 | 73.2806449 | 80.00491151 | 1.260704793 | 1.75E-08 | CELSR3 |
| ENSG00000211448 | 71.03885813 | 42.74131211 | 70.3047269 | 21.10110124 | 27.96235134 | 28.00171903 | 1.259583017 | 0.001913267 | DIO2 |
| ENSG00000158683 | 114.6770138 | 135.5212335 | 127.6301196 | 52.75275311 | 42.42563652 | 64.00392921 | 1.248219977 | 2.49E-06 | PKD1L1 |
| ENSG00000113448 | 1015.855671 | 949.6911057 | 998.327122 | 420.2635998 | 422.3279272 | 406.0249259 | 1.24686746 | 7.24E-45 | PDE4D |
| ENSG00000111012 | 63.93497232 | 65.67567471 | 63.8150598 | 27.25558911 | 15.42750419 | 39.00239436 | 1.24630117 | 0.002230516 | CYP27B1 |
| ENSG00000074590 | 2209.308488 | 2288.223905 | 2105.896974 | 912.6226288 | 961.3263548 | 911.0559299 | 1.245816937 | 1.16E-83 | NUAK1 |
| ENSG00000057657 | 133.9589896 | 107.3745158 | 131.9565643 | 51.87354056 | 50.13938861 | 56.00343806 | 1.240903116 | 1.70E-06 | PRDM1 |
| ENSG00000076826 | 47.69751903 | 47.95366725 | 50.83572561 | 22.85959255 | 25.06969431 | 14.00085951 | 1.237751616 | 0.005356702 | CAMSAP3 |
| ENSG00000088881 | 77.12790312 | 123.0115812 | 89.7737282 | 49.2359029 | 41.46141751 | 32.00196461 | 1.23583332 | 0.000142064 | EBF4 |
| ENSG00000133800 | 1079.790644 | 1035.17373 | 1063.223793 | 415.867537 | 451.2544975 | 489.0300216 | 1.230432409 | 6.14E-43 | LYVE1 |
| ENSG00000160867 | 63.93497232 | 61.5057906 | 74.63117163 | 24.61795145 | 33.74766541 | 27.00165764 | 1.23013812 | 0.000783654 | FGFR4 |
| ENSG00000099282 | 507.4204152 | 568.1467098 | 564.6010376 | 243.5418769 | 247.804286 | 208.0127699 | 1.228056646 | 4.49E-23 | TSPAN15 |
| ENSG00000133805 | 73.06853979 | 95.9073345 | 72.46794927 | 30.77243932 | 34.71188443 | 38.00233297 | 1.224926664 | 0.000235938 | AMPD3 |
| ENSG00000122641 | 1727.259093 | 1577.258664 | 1589.968439 | 716.5582298 | 693.2734695 | 686.0421162 | 1.223353623 | 3.73E-63 | INHBA |
| ENSG00000164442 | 4605.347689 | 4635.868658 | 4829.393933 | 1971.194541 | 2133.816673 | 1926.118245 | 1.222109671 | 9.73E-128 | CITED2 |
| ENSG00000004139 | 194.8494395 | 199.1119662 | 201.1796801 | 95.83416815 | 89.6723681 | 69.00423618 | 1.222014786 | 2.69E-09 | SARM1 |
| ENSG00000261468 | 154.2558062 | 125.0965233 | 126.5485084 | 65.94094139 | 44.35407454 | 64.00392921 | 1.218696326 | 2.82E-06 | AC096921.2 |
| ENSG00000133401 | 66.97949481 | 68.80308779 | 68.14150454 | 30.77243932 | 27.96235134 | 29.00178042 | 1.215716173 | 0.000567374 | PDZD2 |
| ENSG00000091879 | 2977.542997 | 2790.69494 | 2907.37086 | 1193.970645 | 1251.556277 | 1295.079505 | 1.2144987 | 8.38E-98 | ANGPT2 |
| ENSG00000128165 | 100.4692422 | 114.671813 | 86.52889465 | 35.16850207 | 53.03204565 | 43.00263994 | 1.204774426 | 6.70E-05 | ADM2 |
| ENSG00000282057 | 65.96465398 | 56.29343547 | 56.24378152 | 23.7387389 | 30.85500838 | 23.00141206 | 1.20313019 | 0.001960413 | AC092807.3 |
| ENSG00000117586 | 738.8041246 | 688.0308779 | 723.5978815 | 283.1064417 | 300.8363317 | 353.0216721 | 1.200743788 | 2.36E-27 | TNFSF4 |
| ENSG00000262576 | 153.2409654 | 156.3706541 | 117.895619 | 59.78645353 | 62.67423577 | 64.00392921 | 1.198788431 | 9.47E-07 | PCDHGA4 |
| ENSG00000188042 | 247.6211626 | 274.1698801 | 288.7901859 | 114.2976317 | 127.2769096 | 113.0069375 | 1.192569756 | 3.11E-12 | ARL4C |
| ENSG00000064547 | 99.45440139 | 59.42084855 | 67.05989335 | 36.92692718 | 19.28438024 | 43.00263994 | 1.188434688 | 0.002796487 | LPAR2 |
| ENSG00000211445 | 163.3893737 | 129.2664074 | 120.0588413 | 59.78645353 | 62.67423577 | 59.00362224 | 1.186532612 | 2.10E-06 | GPX3 |
| ENSG00000247746 | 62.92013149 | 72.9729719 | 64.89667099 | 35.16850207 | 31.81922739 | 21.00128927 | 1.185029837 | 0.001475096 | USP51 |
| ENSG00000215105 | 71.03885813 | 67.76061676 | 75.71278282 | 32.53086442 | 32.7834464 | 29.00178042 | 1.184029034 | 0.000572212 | TTC3P1 |
| ENSG00000144810 | 8878.842426 | 8418.996015 | 8559.870903 | 3790.285311 | 3762.382584 | 3841.235814 | 1.182458248 | 2.93E-204 | COL8A1 |
| ENSG00000182492 | 15483.42655 | 15519.26618 | 15032.23222 | 6828.84389 | 6920.199848 | 6605.405507 | 1.17730333 | 1.77E-261 | BGN |
| ENSG00000102032 | 158.3151696 | 124.0540522 | 143.8542874 | 73.85385436 | 65.5668928 | 49.0030083 | 1.17379284 | 3.59E-06 | RENBP |
| ENSG00000172031 | 141.0628754 | 139.6911176 | 164.4048998 | 79.12912967 | 63.63845478 | 54.00331527 | 1.172840624 | 1.49E-06 | EPHX4 |
| ENSG00000158373 | 93.3653564 | 72.9729719 | 74.63117163 | 31.65165187 | 34.71188443 | 41.00251715 | 1.170042354 | 0.000413749 | HIST1H2BD |
| ENSG00000133256 | 62.92013149 | 56.29343547 | 63.8150598 | 26.37637656 | 26.03391332 | 29.00178042 | 1.16963882 | 0.001870134 | PDE6B |
| ENSG00000156466 | 367.3723806 | 332.5482577 | 343.9523562 | 149.4661338 | 161.024575 | 154.0094547 | 1.168966615 | 2.01E-15 | GDF6 |
| ENSG00000143603 | 108.5879689 | 90.69497936 | 78.95761637 | 37.80613973 | 53.03204565 | 33.002026 | 1.168927881 | 0.000277673 | KCNN3 |
| ENSG00000171223 | 1440.059138 | 1492.818511 | 1542.377547 | 694.577916 | 684.5954984 | 610.0374503 | 1.168470278 | 1.35E-51 | JUNB |
| ENSG00000156804 | 174.5526228 | 204.3243213 | 164.4048998 | 86.16283008 | 81.958616 | 74.00454315 | 1.164546202 | 4.54E-08 | FBXO32 |
| ENSG00000151014 | 676.8988339 | 722.4324218 | 750.6381611 | 332.3423446 | 331.6913401 | 295.0181112 | 1.163211627 | 6.18E-28 | NOCT |
| ENSG00000158528 | 43.63815571 | 62.54826163 | 49.75411442 | 23.7387389 | 29.89078937 | 16.0009823 | 1.160558763 | 0.008623524 | PPP1R9A |
| ENSG00000144152 | 126.8551038 | 126.1389943 | 107.0795071 | 42.20220249 | 57.85314071 | 62.00380642 | 1.156847399 | 1.99E-05 | FBLN7 |

| | | | | | | | | | |
|---|---|---|---|---|---|---|---|---|---|
| ENSG00000104951 | 258.7844118 | 246.0231624 | 230.383182 | 100.2302309 | 99.31455822 | 131.0080426 | 1.156235099 | 4.82E-10 | IL4I1 |
| ENSG00000242732 | 508.4352561 | 440.9652445 | 494.2963107 | 208.3733748 | 214.0566206 | 226.013875 | 1.155637804 | 1.75E-19 | RTL5 |
| ENSG00000213949 | 251.680526 | 265.8301119 | 188.2003459 | 112.5392066 | 100.2787772 | 104.006385 | 1.155230969 | 4.85E-09 | ITGA1 |
| ENSG00000234465 | 60.89044983 | 57.33590649 | 103.8346736 | 39.56456483 | 24.1054753 | 36.00221018 | 1.15174544 | 0.00329924 | PINLYP |
| ENSG00000197646 | 543.9546851 | 494.1312669 | 574.3355382 | 228.5952635 | 280.5877324 | 218.0133839 | 1.148836032 | 1.89E-18 | PDCD1LG2 |
| ENSG00000206538 | 343.0162007 | 272.0849381 | 316.9120766 | 135.398733 | 145.5970708 | 140.0085951 | 1.147264726 | 1.83E-12 | VGLL3 |
| ENSG00000163762 | 1767.852727 | 1567.876425 | 1557.520104 | 693.6987034 | 769.4467714 | 754.046291 | 1.143227029 | 1.76E-48 | TM4SF18 |
| ENSG00000137801 | 282359.1643 | 266306.5212 | 273726.5869 | 124533.4243 | 122799.0765 | 125189.6854 | 1.14250335 | 0 | THBS1 |
| ENSG00000118523 | 56612.89573 | 55921.27331 | 56927.35979 | 25265.05189 | 25948.09783 | 25692.57727 | 1.139838094 | 0 | CTGF |
| ENSG00000279117 | 2132.180585 | 2173.552092 | 2013.960023 | 941.6366431 | 945.8988506 | 982.0602888 | 1.139555319 | 4.64E-71 | AP001972.5 |
| ENSG00000243243 | 72.05369896 | 56.29343547 | 107.0795071 | 53.63196566 | 23.14125628 | 30.00184182 | 1.132663927 | 0.006103011 | AC073130.2 |
| ENSG00000167992 | 1220.853519 | 1242.625464 | 1273.056363 | 563.5752458 | 612.2790725 | 530.0325388 | 1.130782032 | 2.86E-43 | VWCE |
| ENSG00000128590 | 1483.697294 | 1641.891868 | 1519.663712 | 693.6987034 | 729.9137919 | 699.0429144 | 1.130055606 | 3.11E-52 | DNAJB9 |
| ENSG00000175155 | 1730.303616 | 1559.536657 | 1527.234991 | 763.156495 | 686.5239364 | 753.0462296 | 1.128817765 | 1.27E-47 | YPEL2 |
| ENSG00000105290 | 307.4967716 | 274.1698801 | 307.177576 | 133.6403079 | 126.3126905 | 147.0090249 | 1.12779326 | 3.35E-12 | APLP1 |
| ENSG00000132561 | 240.5172768 | 258.5328147 | 234.7096267 | 91.43810539 | 138.8475377 | 107.0065692 | 1.123821982 | 6.50E-09 | MATN2 |
| ENSG00000099860 | 816.9468685 | 782.8957414 | 841.4935005 | 350.8058082 | 423.2921462 | 348.0213651 | 1.121809523 | 6.28E-27 | GADD45B |
| ENSG00000111339 | 1119.369436 | 1077.915042 | 1062.142182 | 519.6146182 | 500.4296671 | 478.0293463 | 1.120781717 | 1.98E-40 | ART4 |
| ENSG00000100031 | 50.74204152 | 68.80308779 | 75.71278282 | 29.01401421 | 35.67610344 | 25.00153485 | 1.120452632 | 0.00340617 | GGT1 |
| ENSG00000164692 | 181.6565087 | 173.0501905 | 189.281957 | 95.83416815 | 84.85127304 | 69.00423618 | 1.119734795 | 1.65E-07 | COL1A2 |
| ENSG00000138028 | 52.77172318 | 65.67567471 | 58.40700389 | 36.92692718 | 26.03391332 | 18.00110509 | 1.119238449 | 0.007066101 | CGREF1 |
| ENSG00000115266 | 112.6473322 | 77.14285601 | 137.3646203 | 46.59826525 | 53.03204565 | 51.00313109 | 1.11922023 | 0.000235299 | APC2 |
| ENSG00000090339 | 3516.423478 | 3433.899563 | 3681.804467 | 1594.891569 | 1625.673254 | 1681.103203 | 1.11738864 | 4.23E-100 | ICAM1 |
| ENSG00000100292 | 28657.07537 | 30341.11924 | 30932.99823 | 13233.02812 | 14042.88569 | 14243.87443 | 1.115143898 | 7.50E-191 | HMOX1 |
| ENSG00000237238 | 56.83108651 | 57.33590649 | 55.16217034 | 31.65165187 | 14.46328518 | 32.00196461 | 1.114359254 | 0.009668982 | BMS1P10 |
| ENSG00000101335 | 1074.716439 | 1063.320448 | 1070.795071 | 531.0443813 | 522.6067044 | 430.0263994 | 1.111219702 | 8.94E-33 | MYL9 |
| ENSG00000138449 | 2429.528948 | 2474.826218 | 2443.359663 | 1043.625299 | 1260.234248 | 1099.067472 | 1.111211802 | 1.41E-58 | SLC40A1 |
| ENSG00000090530 | 198.9088028 | 205.3667923 | 226.0567373 | 96.71338071 | 107.9925293 | 87.00534127 | 1.110491551 | 1.12E-08 | P3H2 |
| ENSG00000177606 | 8750.972481 | 8618.107981 | 9180.715722 | 4102.405767 | 4183.746292 | 4112.252452 | 1.09849999 | 4.92E-172 | JUN |
| ENSG00000130475 | 60.89044983 | 78.18532704 | 57.3253927 | 28.13480166 | 34.71188443 | 29.00178042 | 1.097784724 | 0.003142953 | FCHO1 |
| ENSG00000187566 | 135.9886713 | 149.0733569 | 150.3439545 | 49.2359029 | 81.958616 | 73.00448176 | 1.096607275 | 1.49E-05 | NHLRC1 |
| ENSG00000223547 | 423.1886263 | 341.9304969 | 383.97197 | 152.1037715 | 206.3428685 | 180.0110509 | 1.095905111 | 2.67E-12 | ZNF844 |
| ENSG00000162654 | 447.5448062 | 447.2200706 | 471.5824758 | 200.4604618 | 227.5556868 | 212.0130155 | 1.094860925 | 1.27E-17 | GBP4 |
| ENSG00000179044 | 127.8699446 | 167.8378354 | 162.2416775 | 72.09542925 | 76.17330193 | 66.004052 | 1.094376712 | 3.20E-06 | EXOC3L1 |
| ENSG00000160323 | 72.05369896 | 94.86486347 | 86.52889465 | 28.13480166 | 42.42563652 | 49.0030083 | 1.089286862 | 0.001287223 | ADAMTS13 |
| ENSG00000183117 | 207.0275294 | 200.1544372 | 180.6290676 | 82.64597988 | 80.99439699 | 114.0069989 | 1.085560062 | 2.88E-07 | CSMD1 |
| ENSG00000234964 | 212.1017336 | 168.8803064 | 176.3026228 | 85.28361753 | 93.52924415 | 84.00515709 | 1.085135579 | 2.22E-07 | FABP5P7 |
| ENSG00000105929 | 79.15758478 | 60.46331957 | 91.93695056 | 37.80613973 | 48.21095059 | 23.00141206 | 1.083576545 | 0.004002499 | ATP6V0A4 |
| ENSG00000167889 | 262.8437751 | 248.1081045 | 276.8924629 | 108.1431439 | 122.4558145 | 142.0087179 | 1.082958893 | 7.81E-10 | MGAT5B |
| ENSG00000188659 | 56.83108651 | 50.0386093 | 68.14150454 | 35.16850207 | 23.14125628 | 24.00147345 | 1.082209641 | 0.007690788 | SAXO2 |
| ENSG00000119630 | 2242.798235 | 2071.389931 | 2199.997146 | 1004.939947 | 1011.465743 | 1069.06563 | 1.078641899 | 3.85E-64 | PGF |
| ENSG00000090661 | 97.42471973 | 114.671813 | 109.2427295 | 62.42409118 | 48.21095059 | 41.00251715 | 1.078232719 | 0.000178504 | CERS4 |
| ENSG00000198108 | 88.29115225 | 66.71814574 | 111.4059519 | 37.80613973 | 40.4971985 | 48.00294691 | 1.078141701 | 0.00111691 | CHSY3 |
| ENSG00000198855 | 459.7228962 | 438.8803024 | 431.5628621 | 188.1514861 | 208.2713066 | 235.0144276 | 1.077444718 | 1.02E-15 | FICD |
| ENSG00000175274 | 2003.295799 | 2078.687228 | 2037.755469 | 984.7180581 | 1007.608867 | 908.0557457 | 1.076605461 | 2.11E-61 | TP53I11 |
| ENSG00000168672 | 487.1235986 | 518.1081005 | 520.2549791 | 252.3340024 | 259.3749142 | 211.0129541 | 1.076090316 | 1.45E-17 | FAM84B |
| ENSG00000026103 | 402.8918097 | 364.8648595 | 322.3201326 | 173.2048727 | 194.7722404 | 149.0091477 | 1.076065727 | 8.53E-12 | FAS |
| ENSG00000166922 | 67.99433564 | 69.84555882 | 54.08055916 | 23.7387389 | 36.64032245 | 31.00190321 | 1.07505819 | 0.004833884 | SCG5 |
| ENSG00000164659 | 2026.637138 | 1975.482596 | 1957.716241 | 988.2349083 | 948.7915076 | 899.0551931 | 1.070573633 | 5.57E-61 | KIAA1324L |
| ENSG00000179598 | 96.4098789 | 91.73745039 | 100.58984 | 38.68535228 | 48.21095059 | 51.00313109 | 1.069344051 | 0.00027407 | PLD6 |
| ENSG00000179546 | 155.2706471 | 151.1582989 | 176.3026228 | 87.92125519 | 76.17330193 | 66.004052 | 1.065253778 | 2.62E-06 | HTR1D |
| ENSG00000105464 | 327.7935882 | 265.8301119 | 287.7085747 | 135.398733 | 161.024575 | 125.0076742 | 1.064651768 | 5.38E-10 | GRIN2D |
| ENSG00000010626 | 58.86076817 | 93.82239244 | 72.46794927 | 36.04771463 | 38.56876047 | 33.002026 | 1.063901825 | 0.002622088 | LRRC23 |
| ENSG00000130513 | 6488.89227 | 6895.945844 | 6917.985127 | 3071.968656 | 3333.305124 | 3315.203521 | 1.062978308 | 8.98E-117 | GDF15 |
| ENSG00000230257 | 118.7363772 | 97.99227655 | 113.5691742 | 62.42409118 | 43.38985553 | 52.00319248 | 1.062582096 | 0.000155882 | NFE4 |
| ENSG00000165246 | 911.3270658 | 817.2972853 | 743.0668828 | 405.3169864 | 390.5086998 | 388.0238208 | 1.062145748 | 1.49E-24 | NLGN4Y |
| ENSG00000155465 | 327.7935882 | 344.015439 | 341.7891339 | 133.6403079 | 167.7741081 | 186.0114193 | 1.059525556 | 3.15E-11 | SLC7A7 |
| ENSG00000124212 | 150.1964429 | 199.1119662 | 160.0784551 | 78.24991712 | 77.13752095 | 90.00552545 | 1.054856925 | 2.48E-06 | PTGIS |
| ENSG00000067445 | 128.8847855 | 103.2046317 | 112.487563 | 45.7190527 | 61.71001676 | 59.00362224 | 1.053537905 | 0.000118195 | TRO |
| ENSG00000121653 | 311.556135 | 297.1042427 | 300.6879089 | 145.9492836 | 152.3466039 | 143.0087793 | 1.043125852 | 1.43E-11 | MAPK8IP1 |
| ENSG00000050165 | 5804.88955 | 5557.413046 | 5743.355382 | 2761.606625 | 2814.555295 | 2725.167298 | 1.043028529 | 6.50E-132 | DKK3 |
| ENSG00000095015 | 801.7242561 | 767.258676 | 721.4346591 | 362.2355714 | 334.5839971 | 418.0256627 | 1.040082545 | 5.87E-22 | MAP3K1 |
| ENSG00000241717 | 394.7730831 | 403.4362875 | 400.1961377 | 175.8425104 | 184.1658313 | 224.0137522 | 1.039344657 | 2.62E-13 | VWFP1 |
| ENSG00000159261 | 90.32083391 | 117.7992261 | 98.42661766 | 41.32298994 | 37.60454146 | 71.00435897 | 1.036253189 | 0.001350748 | CLDN14 |
| ENSG00000174080 | 920.4606332 | 1079.999984 | 1063.223793 | 416.7467496 | 559.2470269 | 520.0319248 | 1.035646118 | 3.12E-21 | CTSF |
| ENSG00000232531 | 62.92013149 | 67.76061676 | 56.24378152 | 32.53086442 | 23.14125628 | 36.00221018 | 1.028426233 | 0.007399481 | AC027612.3 |
| ENSG00000157570 | 3079.02708 | 3201.428524 | 2980.920421 | 1497.298976 | 1589.99715 | 1457.089451 | 1.027189765 | 9.20E-74 | TSPAN18 |
| ENSG00000105810 | 1654.190554 | 1401.08106 | 1300.096642 | 748.2098816 | 671.0964322 | 723.0443878 | 1.023675086 | 8.14E-29 | CDK6 |
| ENSG00000180573 | 207.0275294 | 208.4942054 | 164.4048998 | 94.9549556 | 107.9925293 | 83.0050957 | 1.020154479 | 1.35E-06 | HIST1H2AC |
| ENSG00000063180 | 174.5526228 | 158.4555961 | 154.6703992 | 85.28361753 | 78.10173996 | 77.00472733 | 1.019674072 | 2.70E-06 | CA11 |
| ENSG00000167191 | 445.5151246 | 415.9459398 | 429.3996397 | 218.0447129 | 187.0584883 | 233.0143048 | 1.016749558 | 6.15E-14 | GPRC5B |
| ENSG00000099953 | 220.2204602 | 244.9806914 | 209.8325695 | 94.07574305 | 123.4200335 | 117.0071831 | 1.015816604 | 1.18E-07 | MMP11 |
| ENSG00000123700 | 689.0769239 | 702.6254723 | 700.8840467 | 342.8928952 | 343.2619682 | 349.0214265 | 1.01552954 | 4.73E-24 | KCNJ2 |
| ENSG00000237172 | 328.8084291 | 341.9304969 | 348.278801 | 142.4324334 | 164.881451 | 198.012156 | 1.014920975 | 1.64E-10 | B3GNT9 |
| ENSG00000163874 | 476.9751903 | 444.0926576 | 495.3779219 | 227.7160509 | 289.2657035 | 184.0112965 | 1.014005401 | 3.38E-11 | ZC3H12A |
| ENSG00000254486 | 152.2261246 | 125.0965233 | 161.1600663 | 80.00834222 | 48.21095059 | 89.00546406 | 1.012938161 | 0.000167302 | LINC02547 |
| ENSG00000134070 | 676.8988339 | 614.015435 | 620.8448191 | 285.7440794 | 321.0849309 | 342.0209967 | 1.012764772 | 3.31E-19 | IRAK2 |
| ENSG00000091656 | 95.39503806 | 77.14285601 | 54.08055916 | 35.16850207 | 45.31829356 | 32.00196461 | 1.01228821 | 0.006038279 | ZFHX4 |
| ENSG00000196632 | 104.5286055 | 88.61003731 | 112.487563 | 58.90724098 | 48.21095059 | 44.00270133 | 1.012222203 | 0.000505041 | WNK3 |
| ENSG00000118503 | 900.1638166 | 853.7837712 | 922.6143392 | 441.364701 | 483.0737249 | 404.0248031 | 1.010079611 | 1.54E-25 | TNFAIP3 |
| ENSG00000120129 | 2789.797443 | 2744.826214 | 2714.84407 | 1310.026702 | 1403.902881 | 1385.085031 | 1.009710854 | 3.77E-73 | DUSP1 |
| ENSG00000159167 | 75.09822146 | 67.76061676 | 133.0381755 | 44.83984015 | 49.1751696 | 43.00263994 | 1.0077856 | 0.004179495 | STC1 |
| ENSG00000073756 | 510.4649377 | 487.8764407 | 546.2136475 | 296.29463 | 246.840067 | 224.0137522 | 1.006615047 | 2.28E-14 | PTGS2 |
| ENSG00000109472 | 310.5412941 | 380.5019249 | 316.9120766 | 162.6543221 | 189.9511453 | 149.0091477 | 1.006555774 | 4.61E-10 | CPE |
| ENSG00000276180 | 218.1907786 | 178.2625456 | 165.486511 | 103.7470811 | 86.77971106 | 89.00546406 | 1.006362679 | 2.40E-06 | HIST1H4I |
| ENSG00000204103 | 530.7617543 | 573.3590649 | 576.4987606 | 257.6092777 | 341.3335302 | 238.0146118 | 1.005764261 | 1.75E-13 | MAFB |
| ENSG00000165194 | 116.7066955 | 92.77992141 | 80.03922755 | 53.63196566 | 33.74766541 | 57.00349945 | 1.004403348 | 0.001930727 | PCDH19 |
| ENSG00000253953 | 375.4911073 | 351.3127361 | 306.0959648 | 203.0980995 | 163.917232 | 147.0090249 | 1.004211567 | 1.10E-09 | PCDHGB4 |
| ENSG00000163513 | 17991.09824 | 17370.69473 | 17674.60834 | 8661.122849 | 9085.835748 | 8712.534864 | 1.00328897 | 1.20E-197 | TGFBR2 |
| ENSG00000145632 | 6986.164277 | 6984.555882 | 7164.592477 | 3515.091782 | 3584.966286 | 3449.211748 | 1.002377768 | 6.90E-143 | PLK2 |
| ENSG00000158042 | 1022.959557 | 999.729715 | 963.7155641 | 1921.958638 | 2036.430553 | 2019.123954 | -1.000500653 | 1.18E-54 | MRPL17 |
| ENSG00000100749 | 311.556135 | 320.0386053 | 279.0556852 | 619.8448491 | 630.5992337 | 574.0352401 | -1.002249083 | 8.65E-19 | VRK1 |
| ENSG00000172244 | 64.94981315 | 44.82625417 | 56.24378152 | 99.35101836 | 121.4915955 | 113.0069375 | -1.006047799 | 0.000340611 | C5orf34 |

| | | | | | | | | | |
|---|---|---|---|---|---|---|---|---|---|
| ENSG00000183763 | 69.00917647 | 92.77992141 | 67.05989335 | 163.5335346 | 159.0961369 | 137.008411 | -1.00676159 | 2.44E-05 | TRAIP |
| ENSG00000134955 | 42.62331488 | 44.82625417 | 47.59089206 | 81.76676732 | 87.74393008 | 102.0062622 | -1.006835546 | 0.000970838 | SLC37A2 |
| ENSG00000149548 | 118.7363772 | 120.9266391 | 133.0381755 | 221.5615631 | 260.3391332 | 268.0164536 | -1.00806391 | 1.75E-08 | CCDC15 |
| ENSG00000205403 | 70.0240173 | 66.71814574 | 83.2840611 | 144.1908585 | 153.3108229 | 145.0089021 | -1.009066755 | 1.39E-05 | CFI |
| ENSG00000111684 | 147.1519204 | 123.0115812 | 122.2220637 | 269.0390409 | 263.2317902 | 258.0158396 | -1.009319056 | 3.58E-09 | LPCAT3 |
| ENSG00000134363 | 45.66783737 | 64.63320368 | 85.44728347 | 130.1234577 | 130.1695666 | 133.0081654 | -1.009712293 | 0.000219131 | FST |
| ENSG00000123473 | 380.5653114 | 418.0308819 | 398.0329154 | 764.9149201 | 824.4072551 | 834.0512025 | -1.017650599 | 2.45E-25 | STIL |
| ENSG00000074370 | 27.40070242 | 44.82625417 | 36.77478023 | 75.61227946 | 80.03017798 | 65.00399061 | -1.019376602 | 0.004232934 | ATP2A3 |
| ENSG00000119969 | 318.6600208 | 282.5096484 | 259.5866839 | 599.6229604 | 580.4598451 | 568.0348718 | -1.021352482 | 3.37E-18 | HELLS |
| ENSG00000186767 | 113.662173 | 105.2895737 | 83.2840611 | 201.3396744 | 218.8777157 | 194.0119104 | -1.021388995 | 4.09E-07 | SPIN4 |
| ENSG00000136824 | 760.115782 | 665.0965153 | 688.9863236 | 1424.324334 | 1441.507423 | 1428.087671 | -1.021680367 | 6.06E-41 | SMC2 |
| ENSG00000138658 | 123.8105813 | 112.5868709 | 98.42661766 | 247.9379396 | 203.4502115 | 229.0140592 | -1.022494867 | 8.87E-08 | ZGRF1 |
| ENSG00000006634 | 326.7787474 | 318.9961343 | 270.4027958 | 620.7240616 | 625.7781387 | 617.0378801 | -1.023305978 | 1.57E-19 | DBF4 |
| ENSG00000159259 | 240.5172768 | 259.5752858 | 250.9337945 | 487.9629663 | 549.6048367 | 489.0300216 | -1.023359196 | 9.49E-17 | CHAF1B |
| ENSG00000119640 | 72.05369896 | 82.35521114 | 57.3253927 | 130.1234577 | 150.4181658 | 151.0092705 | -1.024858755 | 2.82E-05 | ACYP1 |
| ENSG00000021645 | 115.6918547 | 108.4169868 | 134.1197867 | 252.3340024 | 242.983191 | 233.0143048 | -1.025253642 | 1.07E-08 | NRXN3 |
| ENSG00000188229 | 5013.313703 | 5001.775988 | 5092.22545 | 10200.62403 | 10583.26787 | 10054.61725 | -1.029544799 | 1.07E-172 | TUBB4B |
| ENSG00000115602 | 13621.19363 | 13126.79517 | 14436.26446 | 27355.81934 | 29130.02057 | 27594.69404 | -1.029726422 | 4.83E-155 | IL1RL1 |
| ENSG00000117650 | 197.8939619 | 260.6177568 | 199.0164577 | 420.2635998 | 457.0398116 | 469.0287938 | -1.033145112 | 4.03E-13 | NEK2 |
| ENSG00000138778 | 790.5610069 | 775.5984442 | 723.5978815 | 1524.554565 | 1541.7862 | 1633.100256 | -1.036473627 | 7.88E-46 | CENPE |
| ENSG00000197635 | 256.7547301 | 234.5559811 | 228.2199596 | 502.9095797 | 492.715915 | 481.0295305 | -1.036814647 | 3.18E-17 | DPP4 |
| ENSG00000211772 | 660.6613806 | 697.4131172 | 736.5772157 | 1447.18386 | 1383.654282 | 1467.090065 | -1.03750413 | 1.07E-42 | TRBC2 |
| ENSG00000152760 | 44.65299654 | 44.82625417 | 35.69316904 | 88.80046774 | 68.45954984 | 100.0061394 | -1.037731203 | 0.001540189 | TCTEX1D1 |
| ENSG00000120334 | 160.3448512 | 146.9884148 | 125.4668972 | 279.5895915 | 341.3335302 | 270.0165764 | -1.040170805 | 2.16E-09 | CENPL |
| ENSG00000128951 | 532.791436 | 552.5096444 | 536.4791468 | 1114.841516 | 1209.130641 | 1011.062069 | -1.040434381 | 1.40E-30 | DUT |
| ENSG00000146147 | 40.59363322 | 33.35907287 | 42.18283614 | 81.76676732 | 79.06595897 | 78.00478873 | -1.040974299 | 0.001385549 | MLIP |
| ENSG00000156802 | 663.7059031 | 580.6563621 | 571.0907047 | 1113.962303 | 1357.620369 | 1277.0784 | -1.044756198 | 1.70E-28 | ATAD2 |
| ENSG00000000460 | 94.38019723 | 101.1196896 | 105.9978959 | 189.9099112 | 199.5933354 | 233.0143048 | -1.045371301 | 1.17E-07 | C1orf112 |
| ENSG00000132967 | 370.4169031 | 358.6100333 | 441.2973627 | 873.9372766 | 776.1963045 | 773.0474575 | -1.051640957 | 2.22E-22 | HMGB1P5 |
| ENSG00000249565 | 77.12790312 | 57.33590649 | 68.14150454 | 132.7610953 | 137.8833187 | 150.0092091 | -1.052525647 | 1.51E-05 | SERBP1P5 |
| ENSG00000125319 | 62.92013149 | 84.4401532 | 82.20244992 | 155.6206217 | 165.84567 | 155.0095161 | -1.054819174 | 3.61E-06 | C17orf53 |
| ENSG00000146263 | 208.0423702 | 240.8108073 | 247.6889609 | 443.1231261 | 498.5012291 | 506.0310653 | -1.055472245 | 7.19E-16 | MMS22L |
| ENSG00000121957 | 253.7102076 | 203.2818503 | 262.8315175 | 472.1371404 | 543.8195227 | 481.0295305 | -1.056363288 | 4.78E-15 | GPSM2 |
| ENSG00000131470 | 65.96465398 | 90.69497936 | 81.12083873 | 160.895897 | 173.5594221 | 160.009823 | -1.057122978 | 2.38E-06 | PSMC3IP |
| ENSG00000232759 | 61.90529066 | 59.42084855 | 64.89667099 | 126.6066075 | 139.8117567 | 121.0074287 | -1.057295651 | 2.09E-05 | AC002480.1 |
| ENSG00000123975 | 310.5412941 | 321.0810764 | 348.278801 | 718.3166549 | 725.0926969 | 594.036468 | -1.057452551 | 1.03E-19 | CKS2 |
| ENSG00000138376 | 185.715872 | 189.7297269 | 168.7313446 | 398.283286 | 370.2601005 | 367.0225316 | -1.061274465 | 5.96E-14 | BARD1 |
| ENSG00000133863 | 93.3653564 | 69.84555882 | 68.14150454 | 162.6543221 | 136.9190997 | 184.0112965 | -1.061566083 | 9.20E-06 | TEX15 |
| ENSG00000119403 | 785.4868028 | 846.486474 | 837.1670557 | 1770.734079 | 1744.272192 | 1643.10087 | -1.063509156 | 9.03E-52 | PHF19 |
| ENSG00000139832 | 48.71235986 | 29.18918876 | 44.34605851 | 86.16283008 | 76.17330193 | 94.00577103 | -1.067613022 | 0.001282301 | RAB20 |
| ENSG00000077152 | 163.3893737 | 199.1119662 | 178.4658452 | 396.5248609 | 390.5086998 | 347.0213037 | -1.06898446 | 3.31E-13 | UBE2T |
| ENSG00000205208 | 350.1200865 | 282.5096484 | 277.9740741 | 621.6032742 | 658.5615851 | 643.0394763 | -1.07719539 | 4.14E-20 | C4orf46 |
| ENSG00000115896 | 365.342699 | 325.2509605 | 309.3407984 | 674.3560273 | 716.4147258 | 723.0443878 | -1.078573654 | 8.48E-24 | PLCL1 |
| ENSG00000101868 | 280.0960692 | 262.7026988 | 270.4027958 | 560.0583955 | 642.1698619 | 516.0316793 | -1.079250137 | 2.36E-18 | POLA1 |
| ENSG00000185480 | 173.537782 | 170.9652484 | 124.3852861 | 320.9125814 | 334.5839971 | 339.0208125 | -1.082645234 | 3.38E-11 | PARPBP |
| ENSG00000119514 | 31.46006574 | 28.14671773 | 19.4690013 | 61.54487663 | 48.21095059 | 58.00356085 | -1.083087392 | 0.008881001 | GALNT12 |
| ENSG00000168496 | 813.902346 | 788.1080965 | 814.4532209 | 1732.048727 | 1706.667651 | 1686.10351 | -1.084795117 | 1.61E-58 | FEN1 |
| ENSG00000173207 | 289.2296367 | 301.2741268 | 335.2994668 | 650.6172884 | 674.9533083 | 642.0394149 | -1.08855814 | 1.79E-23 | CKS1B |
| ENSG00000164032 | 1964.731848 | 2022.393793 | 2168.630422 | 4406.61331 | 4439.26433 | 4251.260986 | -1.089764868 | 1.98E-102 | H2AFZ |
| ENSG00000058804 | 565.2663426 | 623.3976742 | 578.661983 | 1254.636312 | 1285.303943 | 1243.076313 | -1.09805069 | 1.21E-43 | NDC1 |
| ENSG00000279348 | 23.3413391 | 54.20849341 | 30.28511313 | 72.97464181 | 61.71001676 | 97.00595521 | -1.102769902 | 0.005567779 | AC012513.3 |
| ENSG00000197594 | 80.17242561 | 79.22779806 | 63.8150598 | 151.2245589 | 182.2373932 | 147.0090249 | -1.104456228 | 1.74E-06 | ENPP1 |
| ENSG00000223865 | 37.54911073 | 29.18918876 | 35.69316904 | 81.76676732 | 66.53111182 | 72.00442036 | -1.105821745 | 0.001395035 | HLA-DPB1 |
| ENSG00000138346 | 126.8551038 | 116.756755 | 160.0784551 | 291.0193547 | 279.6235134 | 300.0184182 | -1.110285315 | 9.05E-11 | DNA2 |
| ENSG00000151470 | 69.00917647 | 77.14285601 | 67.05989335 | 162.6543221 | 159.0961369 | 141.0086565 | -1.118590333 | 9.36E-07 | C4orf33 |
| ENSG00000231607 | 64.94981315 | 47.95366725 | 40.01961377 | 122.2105447 | 112.8136244 | 98.0060166 | -1.12139833 | 0.000113576 | DLEU2 |
| ENSG00000253669 | 27.40070242 | 50.0386093 | 22.71383485 | 70.33700415 | 65.5668928 | 83.0050957 | -1.126221163 | 0.003766789 | GASAL1 |
| ENSG00000164087 | 162.3745329 | 157.4131251 | 151.4255656 | 341.1344701 | 385.6876047 | 302.018541 | -1.126544343 | 7.89E-13 | POC1A |
| ENSG00000143248 | 13552.18445 | 13213.32027 | 12995.55836 | 28488.24511 | 29891.75359 | 28568.75384 | -1.12872214 | 5.49E-266 | RGS5 |
| ENSG00000100629 | 114.6770138 | 109.4594578 | 88.69217701 | 213.6486501 | 259.3749142 | 212.0130155 | -1.129049482 | 7.85E-09 | CEP128 |
| ENSG00000181938 | 234.4282318 | 223.0887998 | 206.587736 | 455.4321019 | 516.8213903 | 483.0296533 | -1.130539213 | 1.04E-18 | GINS3 |
| ENSG00000112118 | 1119.369436 | 1237.413109 | 1158.405577 | 2555.870888 | 2612.069303 | 2563.157353 | -1.137148796 | 2.99E-81 | MCM3 |
| ENSG00000169851 | 469.8713045 | 454.5173678 | 417.5019167 | 967.1338071 | 1024.96481 | 961.0589996 | -1.137248546 | 4.26E-36 | PCDH7 |
| ENSG00000171345 | 696.1808097 | 762.0463208 | 726.842715 | 1647.644322 | 1607.353093 | 1553.095345 | -1.138300863 | 6.35E-56 | KRT19 |
| ENSG00000198554 | 310.5412941 | 306.486482 | 345.0339674 | 684.0273654 | 685.5597174 | 755.0463524 | -1.143143668 | 1.94E-26 | WDHD1 |
| ENSG00000229855 | 214.1314152 | 202.2393793 | 231.4647932 | 457.190527 | 454.1471546 | 520.0319248 | -1.143536295 | 2.50E-18 | AC008568.1 |
| ENSG00000133119 | 292.2741592 | 266.8725829 | 282.3005188 | 600.5021729 | 662.4184611 | 597.0366522 | -1.144017756 | 4.34E-24 | RFC3 |
| ENSG00000114346 | 599.7709308 | 568.1467098 | 562.4378152 | 1155.285293 | 1336.40755 | 1348.082759 | -1.148827493 | 6.50E-40 | ECT2 |
| ENSG00000132423 | 52.77172318 | 50.0386093 | 47.59089206 | 139.7947957 | 99.31455822 | 94.00577103 | -1.149377868 | 7.83E-05 | COQ3 |
| ENSG00000184344 | 28.41554325 | 33.35907287 | 37.85639141 | 78.24991712 | 78.10173996 | 66.004052 | -1.160980867 | 0.000767358 | GDF3 |
| ENSG00000166394 | 33.48974741 | 61.5057906 | 59.48861507 | 131.0026702 | 78.10173996 | 136.0083496 | -1.162206081 | 0.000411473 | CYB5R2 |
| ENSG00000175305 | 92.35051557 | 80.27026909 | 99.50822885 | 183.7554233 | 256.4822571 | 169.0103756 | -1.163047433 | 2.30E-07 | CCNE2 |
| ENSG00000142856 | 93.3653564 | 93.82239244 | 90.85533938 | 192.5475489 | 218.8777157 | 212.0130155 | -1.163955702 | 1.86E-09 | ITGB3BP |
| ENSG00000167767 | 314.6006574 | 288.7644745 | 341.7891339 | 676.1144524 | 744.3770771 | 708.0434669 | -1.171419711 | 3.41E-27 | KRT80 |
| ENSG00000140525 | 480.0197128 | 526.4478687 | 533.2343133 | 1120.116791 | 1190.81048 | 1169.07177 | -1.176687949 | 2.48E-44 | FANCI |
| ENSG00000147642 | 28.41554325 | 45.86872519 | 40.01961377 | 81.76676732 | 101.2429962 | 77.00472733 | -1.187239769 | 0.000340818 | SYBU |
| ENSG00000137135 | 38.56395156 | 44.82625417 | 62.73344862 | 119.5729071 | 120.5273765 | 92.00564824 | -1.188594463 | 6.11E-05 | ARHGEF39 |
| ENSG00000140534 | 144.1073979 | 156.3706541 | 165.486511 | 332.3423446 | 368.3316625 | 362.0222246 | -1.189627163 | 1.46E-15 | TICRR |
| ENSG00000163507 | 350.1200865 | 300.2316558 | 346.1155786 | 744.6930314 | 776.1963045 | 756.0464138 | -1.192071874 | 1.34E-30 | CIP2A |
| ENSG00000101945 | 131.929308 | 151.1582989 | 142.7726762 | 307.7243932 | 334.5839971 | 331.0203214 | -1.1923977 | 1.40E-14 | SUV39H1 |
| ENSG00000147536 | 216.1610969 | 213.7065606 | 194.690013 | 423.78045 | 549.6048367 | 458.0281184 | -1.195257564 | 1.85E-17 | GINS4 |
| ENSG00000138794 | 268.9328201 | 274.1698801 | 271.484407 | 608.4150859 | 627.7065767 | 630.0386782 | -1.195722614 | 9.00E-28 | CASP6 |
| ENSG00000144395 | 41.60847405 | 50.0386093 | 38.93800259 | 99.35101836 | 107.9925293 | 92.00564824 | -1.196561046 | 4.37E-05 | CCDC150 |
| ENSG00000197467 | 1383.228052 | 1287.451719 | 1287.117308 | 3111.533221 | 3026.683478 | 2938.180375 | -1.197359295 | 9.01E-99 | COL13A1 |
| ENSG00000181019 | 1072.686758 | 1190.501913 | 986.429399 | 2438.056406 | 2532.039125 | 2487.152687 | -1.197861398 | 1.94E-69 | NQO1 |
| ENSG00000159055 | 113.662173 | 106.3320448 | 124.3852861 | 283.1064417 | 253.5896001 | 254.0155941 | -1.200640267 | 5.06E-12 | MIS18A |
| ENSG00000173597 | 3049.596696 | 2965.830072 | 3084.755094 | 6898.301682 | 6916.342972 | 7192.441545 | -1.206831164 | 1.29E-187 | SULT1B1 |
| ENSG00000082153 | 3734.614256 | 3821.698785 | 3889.473814 | 8494.951676 | 9202.506249 | 8729.535908 | -1.20719533 | 1.42E-181 | BZW1 |
| ENSG00000105851 | 124.8254221 | 135.5212335 | 110.3243407 | 271.6766785 | 292.1583606 | 299.0183568 | -1.217493281 | 3.58E-13 | PIK3CG |
| ENSG00000092470 | 223.2649827 | 174.0926615 | 229.3015708 | 452.7944642 | 547.6763987 | 459.0281798 | -1.21968179 | 1.94E-17 | WDR76 |
| ENSG00000228716 | 484.0790761 | 496.2162089 | 525.663035 | 1207.158834 | 1242.878306 | 1059.065016 | -1.221351941 | 1.36E-41 | DHFR |
| ENSG00000080839 | 237.4727543 | 241.8532783 | 213.0774031 | 518.7354056 | 526.4635805 | 575.0353015 | -1.225253903 | 1.47E-23 | RBL1 |
| ENSG00000167670 | 412.0253772 | 453.4748968 | 432.6444732 | 1039.229236 | 1036.535438 | 958.0588154 | -1.225327414 | 8.71E-42 | CHAF1A |

| | | | | | | | | | |
|---|---|---|---|---|---|---|---|---|---|
| ENSG00000154920 | 42.62331488 | 40.65637006 | 49.75411442 | 110.7807815 | 109.9209673 | 90.00552545 | -1.226036064 | 2.22E-05 | EME1 |
| ENSG00000101000 | 2488.389716 | 2571.776024 | 2489.868944 | 5652.457496 | 6228.854816 | 5782.35498 | -1.226069564 | 3.02E-142 | PROCR |
| ENSG00000164867 | 1620.700806 | 1608.532795 | 1731.659504 | 3959.973334 | 3915.693407 | 3758.230718 | -1.230200092 | 4.26E-123 | NOS3 |
| ENSG00000169684 | 27.40070242 | 52.12355136 | 47.59089206 | 100.2302309 | 107.9925293 | 92.00564824 | -1.242747645 | 8.25E-05 | CHRNA5 |
| ENSG00000149503 | 771.2790312 | 784.9806834 | 785.2497189 | 1849.863209 | 1907.225205 | 1798.110386 | -1.24658637 | 4.62E-77 | INCENP |
| ENSG00000111206 | 917.4161108 | 879.8455469 | 866.3705577 | 2082.854535 | 2221.560603 | 2026.124384 | -1.248666752 | 1.43E-78 | FOXM1 |
| ENSG00000235387 | 263.8586159 | 253.3204596 | 230.383182 | 535.4404441 | 610.3506345 | 634.0389238 | -1.249695146 | 1.16E-24 | SPAAR |
| ENSG00000140379 | 19.28197578 | 12.50965233 | 31.36672431 | 51.87354056 | 52.06782664 | 46.00282412 | -1.251754175 | 0.007177204 | BCL2A1 |
| ENSG00000164104 | 1120.384277 | 1098.764463 | 1124.87563 | 2668.410095 | 2724.882927 | 2590.15901 | -1.255566234 | 1.87E-105 | HMGB2 |
| ENSG00000146918 | 599.7709308 | 503.5135061 | 548.3768698 | 1367.175518 | 1231.307678 | 1402.086074 | -1.275926698 | 2.22E-47 | NCAPG2 |
| ENSG00000088305 | 105.5434464 | 166.7953643 | 140.6094538 | 369.2692718 | 330.7271211 | 299.0183568 | -1.276657869 | 1.09E-12 | DNMT3B |
| ENSG00000171757 | 18.26713495 | 20.84942054 | 20.55061248 | 56.26960332 | 54.96048367 | 33.002026 | -1.276699805 | 0.005771555 | LRRC34 |
| ENSG00000166508 | 1404.539709 | 1471.96909 | 1570.499438 | 3515.970995 | 3738.277109 | 3518.215984 | -1.276864896 | 1.08E-112 | MCM7 |
| ENSG00000172927 | 33.48974741 | 28.14671773 | 49.75411442 | 96.71338071 | 81.958616 | 91.00558685 | -1.279108159 | 9.57E-05 | MYEOV |
| ENSG00000125885 | 216.1610969 | 240.8108073 | 194.690013 | 508.184855 | 547.6763987 | 527.0323546 | -1.279362752 | 4.07E-24 | MCM8 |
| ENSG00000167513 | 333.8826332 | 399.2664034 | 402.3593601 | 890.642315 | 973.8612019 | 891.054702 | -1.279799969 | 9.25E-37 | CDT1 |
| ENSG00000075702 | 187.7455536 | 202.2393793 | 191.4451794 | 480.0500533 | 454.1471546 | 478.0293463 | -1.280498348 | 1.65E-23 | WDR62 |
| ENSG00000171476 | 52.77172318 | 29.18918876 | 27.04027958 | 83.52519243 | 76.17330193 | 106.0065078 | -1.280742396 | 0.000285948 | HOPX |
| ENSG00000164109 | 281.11091 | 298.1467138 | 299.6062977 | 675.2352398 | 742.4486391 | 727.0446334 | -1.286909215 | 8.77E-34 | MAD2L1 |
| ENSG00000149781 | 1033.107965 | 1134.208478 | 1117.304352 | 2751.935287 | 2688.242605 | 2596.159379 | -1.291440119 | 1.82E-100 | FERMT3 |
| ENSG00000101842 | 36.5342699 | 18.76447849 | 31.36672431 | 67.69936649 | 60.74579774 | 84.00515709 | -1.291867107 | 0.00071447 | VSIG1 |
| ENSG00000113810 | 1404.539709 | 1205.096507 | 1268.729918 | 3185.387075 | 3274.487764 | 3058.187743 | -1.294928108 | 5.32E-96 | SMC4 |
| ENSG00000186638 | 54.80140485 | 78.18532704 | 61.65183744 | 152.1037715 | 136.9190997 | 189.0116035 | -1.295763956 | 1.47E-07 | KIF24 |
| ENSG00000010292 | 1067.612554 | 1123.783767 | 1117.304352 | 2646.429781 | 2800.09201 | 2678.164413 | -1.296140851 | 2.64E-108 | NCAPD2 |
| ENSG00000140465 | 184.7010311 | 143.8610017 | 135.2013979 | 379.8198224 | 417.5068321 | 348.0213651 | -1.302862208 | 2.68E-16 | CYP1A1 |
| ENSG00000203760 | 133.9589896 | 167.8378354 | 135.2013979 | 375.4237597 | 361.5821294 | 344.0211195 | -1.307122939 | 1.89E-17 | CENPW |
| ENSG00000228649 | 22.32649827 | 34.4015439 | 55.16217034 | 90.55889284 | 86.77971106 | 99.006078 | -1.308899275 | 0.000182116 | SNHG26 |
| ENSG00000186193 | 276.0367059 | 310.6563661 | 279.0556852 | 771.069408 | 719.3073828 | 655.040213 | -1.310101468 | 1.64E-31 | SAPCD2 |
| ENSG00000185347 | 105.5434464 | 157.4131251 | 184.9555123 | 390.370373 | 368.3316625 | 355.0217948 | -1.316991132 | 1.20E-13 | TEDC1 |
| ENSG00000285108 | 19.28197578 | 21.89189157 | 25.95866839 | 77.37070456 | 39.53297948 | 50.0030697 | -1.319203104 | 0.003017558 | AC103718.1 |
| ENSG00000094804 | 329.8232699 | 312.7413081 | 373.1558582 | 835.2519243 | 841.7631973 | 857.0526146 | -1.319647282 | 2.21E-39 | CDC6 |
| ENSG00000162073 | 133.9589896 | 151.1582989 | 146.0175097 | 392.1287981 | 363.5105675 | 325.019953 | -1.327268954 | 6.24E-18 | PAQR4 |
| ENSG00000112029 | 199.9236436 | 169.9227774 | 233.6280156 | 494.9966667 | 494.6443531 | 525.0322318 | -1.328459527 | 1.31E-22 | FBXO5 |
| ENSG00000076003 | 696.1808097 | 727.6447769 | 723.5978815 | 1721.498177 | 2010.39664 | 1669.102466 | -1.330701512 | 3.06E-60 | MCM6 |
| ENSG00000137310 | 511.4797786 | 459.729723 | 498.6227554 | 1184.299307 | 1316.158951 | 1211.074348 | -1.336016676 | 1.67E-54 | TCF19 |
| ENSG00000129173 | 394.7730831 | 433.6679473 | 432.6444732 | 1025.161835 | 1091.495921 | 1074.065937 | -1.33940081 | 2.36E-51 | E2F8 |
| ENSG00000120802 | 1203.601225 | 1301.003842 | 1188.69069 | 3022.732753 | 3313.056525 | 3011.184857 | -1.339407745 | 3.36E-105 | TMPO |
| ENSG00000169247 | 13.1929308 | 20.84942054 | 16.22416775 | 51.87354056 | 40.4971985 | 35.00214879 | -1.344973964 | 0.006131412 | SH3TC2 |
| ENSG00000180385 | 37.54911073 | 42.74131211 | 40.01961377 | 88.80046774 | 103.1714343 | 114.0069989 | -1.345357915 | 4.56E-06 | EMC3-AS1 |
| ENSG00000136928 | 720.5369896 | 766.2162049 | 701.9656578 | 1754.908254 | 1887.940825 | 1927.118306 | -1.346906086 | 2.72E-78 | GABBR2 |
| ENSG00000128408 | 19.28197578 | 19.80694952 | 21.63222366 | 50.11511546 | 43.38985553 | 61.00374503 | -1.347248075 | 0.001726148 | RIBC2 |
| ENSG00000177602 | 75.09822146 | 94.86486347 | 96.2633953 | 232.9913262 | 221.7703727 | 224.0137522 | -1.352063683 | 2.17E-12 | HASPIN |
| ENSG00000187741 | 296.3335225 | 344.015439 | 325.5649661 | 866.9035761 | 757.8761433 | 840.0515709 | -1.352328727 | 1.14E-37 | FANCA |
| ENSG00000100479 | 96.4098789 | 84.4401532 | 87.61050583 | 215.4070752 | 216.9492777 | 255.0156555 | -1.354854209 | 1.79E-12 | POLE2 |
| ENSG00000121621 | 226.3095052 | 189.7297269 | 219.5670702 | 542.4741445 | 583.3525021 | 500.030697 | -1.355167544 | 4.56E-26 | KIF18A |
| ENSG00000139618 | 181.6565087 | 175.1351326 | 169.8129557 | 439.6062759 | 431.0058983 | 479.0294077 | -1.356901641 | 4.01E-24 | BRCA2 |
| ENSG00000166845 | 179.626827 | 188.6872559 | 169.8129557 | 429.9349379 | 498.5012291 | 451.0276887 | -1.357203048 | 1.13E-23 | C18orf54 |
| ENSG00000167900 | 496.2571661 | 516.0231584 | 607.8654849 | 1258.153162 | 1543.714638 | 1350.082882 | -1.358146981 | 2.64E-44 | TK1 |
| ENSG00000147697 | 60.89044983 | 52.12355136 | 64.89667099 | 166.1711723 | 149.4539468 | 142.0087179 | -1.364694738 | 7.86E-09 | GSDMC |
| ENSG00000175832 | 187.7455536 | 208.4942054 | 197.9348465 | 503.7887922 | 563.1039029 | 463.0284254 | -1.364962349 | 5.45E-25 | ETV4 |
| ENSG00000213707 | 16.23745329 | 20.84942054 | 19.4690013 | 48.35669035 | 43.38985553 | 54.00331527 | -1.366316806 | 0.001952212 | HMGB1P10 |
| ENSG00000164167 | 66.97949481 | 59.42084855 | 68.14150454 | 184.6346359 | 168.7383271 | 148.0090863 | -1.367401333 | 1.66E-09 | LSM6 |
| ENSG00000129514 | 36.5342699 | 35.44401492 | 22.71383485 | 77.37070456 | 95.45768217 | 74.00454315 | -1.3791582 | 5.89E-05 | FOXA1 |
| ENSG00000012048 | 279.0812284 | 281.4671773 | 245.5257386 | 683.1481528 | 709.6651927 | 717.0440194 | -1.387008013 | 3.13E-37 | BRCA1 |
| ENSG00000087586 | 338.9568374 | 413.8609978 | 359.0949128 | 925.8108171 | 1022.072153 | 971.0596135 | -1.392258951 | 1.85E-44 | AURKA |
| ENSG00000100297 | 885.956045 | 950.7335767 | 975.6132872 | 2403.767117 | 2570.607886 | 2424.148819 | -1.395856265 | 1.13E-105 | MCM5 |
| ENSG00000135476 | 312.5709758 | 335.6756707 | 341.7891339 | 938.1197928 | 861.0475776 | 814.0499747 | -1.401588726 | 3.61E-43 | ESPL1 |
| ENSG00000101695 | 24.35617993 | 26.06177568 | 10.81611183 | 54.51117822 | 52.06782664 | 56.00343806 | -1.404274576 | 0.001507575 | RNF125 |
| ENSG00000073111 | 512.4946194 | 498.301151 | 523.4998126 | 1363.658668 | 1399.081786 | 1298.079689 | -1.404549888 | 9.65E-70 | MCM2 |
| ENSG00000097046 | 105.5434464 | 139.6911176 | 104.9162848 | 301.5699053 | 324.941807 | 302.018541 | -1.406616045 | 1.65E-16 | CDC7 |
| ENSG00000167325 | 1447.163024 | 1294.749016 | 1300.096642 | 3290.892582 | 3773.953212 | 3665.225009 | -1.407749156 | 6.61E-105 | RRM1 |
| ENSG00000118276 | 232.3985502 | 214.7490316 | 206.587736 | 597.8645353 | 564.0681219 | 575.0353015 | -1.409345236 | 1.59E-32 | B4GALT6 |
| ENSG00000212232 | 21.31165744 | 17.72200746 | 9.734500648 | 55.39039077 | 44.35407454 | 30.00184182 | -1.410513505 | 0.006398276 | SNORD17 |
| ENSG00000214826 | 102.4989239 | 102.1621607 | 148.1807321 | 336.7384074 | 247.804286 | 353.0216721 | -1.412025392 | 7.43E-13 | DDX12P |
| ENSG00000206432 | 14.20777163 | 23.97683362 | 38.93800259 | 62.42409118 | 61.71001676 | 81.00497291 | -1.415731239 | 0.000630811 | TMEM200C |
| ENSG00000129810 | 74.08338062 | 80.27026909 | 71.38633808 | 184.6346359 | 219.8419347 | 199.0122174 | -1.417647803 | 4.79E-12 | SGO1 |
| ENSG00000171320 | 138.0183529 | 115.714284 | 124.3852861 | 296.29463 | 341.3335302 | 376.0230841 | -1.420728901 | 8.39E-18 | ESCO2 |
| ENSG00000013810 | 1018.900194 | 947.6061637 | 1080.529572 | 2729.954974 | 2797.199353 | 2645.162387 | -1.423788201 | 8.35E-114 | TACC3 |
| ENSG00000226167 | 16.23745329 | 9.382239244 | 17.30577893 | 33.41007697 | 29.89078937 | 52.00319248 | -1.42404709 | 0.008519324 | AP4B1-AS1 |
| ENSG00000101412 | 274.0070242 | 240.8108073 | 244.4441274 | 654.1341386 | 743.4128581 | 644.0395377 | -1.4262411 | 9.66E-35 | E2F1 |
| ENSG00000089685 | 470.8861453 | 483.7065566 | 554.8665369 | 1311.785127 | 1334.479112 | 1414.086811 | -1.428188828 | 4.73E-63 | BIRC5 |
| ENSG00000111788 | 38.56395156 | 47.95366725 | 23.79544603 | 128.3650326 | 83.88705403 | 85.00521848 | -1.430074807 | 2.98E-05 | AC009533.1 |
| ENSG00000123219 | 147.1519204 | 136.5637046 | 188.2003459 | 431.693363 | 422.3279272 | 416.0255399 | -1.430182693 | 5.21E-22 | CENPK |
| ENSG00000117593 | 377.5207889 | 410.7335847 | 400.1961377 | 1032.195536 | 1097.281235 | 1093.067104 | -1.439059113 | 9.17E-59 | DARS2 |
| ENSG00000085840 | 135.9886713 | 170.9652484 | 118.9772301 | 368.3900592 | 349.0472823 | 439.0269519 | -1.439519849 | 5.10E-18 | ORC1 |
| ENSG00000104738 | 1340.604737 | 1318.725849 | 1263.321862 | 3391.122813 | 3733.456014 | 3549.217887 | -1.443618713 | 3.22E-142 | MCM4 |
| ENSG00000137807 | 470.8861453 | 448.2625417 | 471.5824758 | 1174.627969 | 1298.803009 | 1327.08147 | -1.449706493 | 8.93E-64 | KIF23 |
| ENSG00000127423 | 38.56395156 | 29.18918876 | 35.69316904 | 101.1094435 | 95.45768217 | 87.00534127 | -1.455376368 | 2.23E-06 | AUNIP |
| ENSG00000113368 | 1129.517844 | 1160.270253 | 1174.629745 | 3069.331019 | 3251.346508 | 3203.196645 | -1.458895659 | 1.85E-150 | LMNB1 |
| ENSG00000116574 | 36.5342699 | 17.72200746 | 20.55061248 | 72.09542925 | 67.49533083 | 67.00411339 | -1.462280855 | 0.000173411 | RHOU |
| ENSG00000176208 | 96.4098789 | 105.2895737 | 135.2013979 | 325.3086442 | 298.9078937 | 305.0187251 | -1.466389694 | 2.04E-17 | ATAD5 |
| ENSG00000163584 | 423.1886263 | 429.4980632 | 365.5845799 | 1075.276951 | 1202.381108 | 1121.068823 | -1.478974496 | 1.90E-56 | RPL22L1 |
| ENSG00000111665 | 194.8494395 | 142.8185307 | 165.486511 | 465.9826525 | 464.7535637 | 473.0290393 | -1.479228534 | 7.26E-26 | CDCA3 |
| ENSG00000136982 | 101.484083 | 83.39768217 | 95.18178411 | 218.9239254 | 283.4803895 | 280.0171903 | -1.480385121 | 9.94E-15 | DSCC1 |
| ENSG00000011426 | 1314.218875 | 1186.332029 | 1237.363193 | 3260.120142 | 3661.139588 | 3517.215922 | -1.481033976 | 9.94E-129 | ANLN |
| ENSG00000100162 | 56.83108651 | 70.88802985 | 74.63117163 | 198.7020367 | 197.6648974 | 169.0103756 | -1.484884334 | 9.85E-12 | CENPM |
| ENSG00000163535 | 279.0812284 | 291.8918876 | 294.1982418 | 755.2435821 | 847.5485114 | 823.0505272 | -1.487149759 | 1.51E-46 | SGO2 |
| ENSG00000166803 | 97.42471973 | 97.99227655 | 122.2220637 | 309.4828183 | 288.3014845 | 292.017927 | -1.488159553 | 3.47E-18 | PCLAF |
| ENSG00000122952 | 392.7434014 | 432.6254763 | 577.5803718 | 1312.66434 | 1425.115699 | 1203.073857 | -1.49064544 | 1.21E-14 | ZWINT |
| ENSG00000215784 | 23.3413391 | 20.84942054 | 19.4690013 | 75.61227946 | 48.21095059 | 55.00337667 | -1.491511116 | 0.000282495 | FAM72D |
| ENSG00000138182 | 344.0310415 | 342.9729679 | 296.3614642 | 851.9569628 | 959.3979168 | 958.0588154 | -1.492218144 | 4.94E-48 | KIF20B |
| ENSG00000068489 | 590.6373633 | 548.3397603 | 526.7446462 | 1503.453464 | 1697.025461 | 1493.091661 | -1.493808419 | 5.47E-73 | PRR11 |
| ENSG00000256663 | 42.62331488 | 59.42084855 | 52.99894797 | 139.7947957 | 149.4539468 | 148.0090863 | -1.496601542 | 1.26E-09 | AC112777.1 |

| | | | | | | | | | |
|---|---|---|---|---|---|---|---|---|---|
| ENSG00000161800 | 315.6154983 | 373.2046277 | 377.4823029 | 951.3079811 | 1029.785905 | 1032.063359 | -1.499169635 | 1.39E-52 | RACGAP1 |
| ENSG00000111247 | 159.3300104 | 142.8185307 | 148.1807321 | 416.7467496 | 443.5407454 | 418.0256627 | -1.504605138 | 3.61E-27 | RAD51AP1 |
| ENSG00000140451 | 150.1964429 | 167.8378354 | 173.0577893 | 505.5472173 | 470.5388778 | 425.0260924 | -1.514333451 | 1.55E-27 | PIF1 |
| ENSG00000242193 | 145.1222388 | 127.1814653 | 111.4059519 | 389.4911605 | 336.5124351 | 375.0230227 | -1.519508551 | 8.88E-22 | CRYZL2P |
| ENSG00000090889 | 415.0698997 | 353.3976782 | 338.5443003 | 1030.437111 | 1111.744521 | 1048.064341 | -1.52566455 | 9.67E-56 | KIF4A |
| ENSG00000146410 | 38.56395156 | 37.52895698 | 34.61155786 | 105.5055062 | 108.9567483 | 105.0064464 | -1.528137272 | 8.44E-08 | MTFR2 |
| ENSG00000100558 | 51.75688235 | 57.33590649 | 86.52889465 | 183.7554233 | 194.7722404 | 185.0113579 | -1.529745524 | 3.74E-11 | PLEK2 |
| ENSG00000249992 | 40.59363322 | 62.54826163 | 83.2840611 | 181.9969982 | 181.2731742 | 174.0106825 | -1.531211377 | 6.96E-10 | TMEM158 |
| ENSG00000123080 | 131.929308 | 159.4980672 | 139.5278426 | 424.6596626 | 448.3618405 | 384.0235753 | -1.545043275 | 9.55E-26 | CDKN2C |
| ENSG00000178401 | 14.20777163 | 15.63706541 | 14.06094538 | 38.68535228 | 43.38985553 | 47.00288551 | -1.554293765 | 0.00109958 | DNAJC22 |
| ENSG00000165480 | 190.7900761 | 187.6447849 | 177.384234 | 524.0106809 | 615.1717295 | 497.0305128 | -1.557194884 | 1.11E-31 | SKA3 |
| ENSG00000122966 | 294.3038408 | 296.0617717 | 250.9337945 | 824.7013737 | 844.6558544 | 820.050343 | -1.564040667 | 4.53E-51 | CIT |
| ENSG00000101447 | 327.7935882 | 371.1196857 | 348.278801 | 932.8445175 | 1101.138111 | 1073.065876 | -1.568532934 | 4.59E-55 | FAM83D |
| ENSG00000101003 | 237.4727543 | 211.6216185 | 223.8935149 | 686.665003 | 672.0606512 | 638.0391693 | -1.569076315 | 2.46E-43 | GINS1 |
| ENSG00000265415 | 72.05369896 | 70.88802985 | 71.38633808 | 215.4070752 | 208.2713066 | 219.0134453 | -1.584213193 | 1.11E-15 | AC099850.3 |
| ENSG00000198056 | 55.81624568 | 33.35907287 | 61.65183744 | 137.1571581 | 157.1676989 | 158.0097002 | -1.584729919 | 1.64E-09 | PRIM1 |
| ENSG00000102384 | 129.8996263 | 120.9266391 | 123.3036749 | 320.9125814 | 382.7949477 | 420.0257854 | -1.58492411 | 8.56E-23 | CENPI |
| ENSG00000143476 | 202.9681661 | 201.1969082 | 186.0371235 | 587.3139847 | 588.1735972 | 600.0368364 | -1.58821609 | 8.93E-41 | DTL |
| ENSG00000181544 | 26.38586159 | 28.14671773 | 47.59089206 | 108.1431439 | 94.49346316 | 107.0065692 | -1.60480527 | 3.82E-07 | FANCB |
| ENSG00000079616 | 420.1441038 | 413.8609978 | 385.0535812 | 1246.723399 | 1268.91222 | 1202.073796 | -1.608203952 | 7.80E-80 | KIF22 |
| ENSG00000153044 | 58.86076817 | 75.05791395 | 61.65183744 | 169.6880225 | 228.5199058 | 201.0123402 | -1.614318551 | 3.27E-13 | CENPH |
| ENSG00000276043 | 655.5871765 | 651.544392 | 639.2322092 | 1968.556904 | 2015.217735 | 1999.122726 | -1.619782113 | 8.90E-129 | UHRF1 |
| ENSG00000104147 | 50.74204152 | 43.78378314 | 57.3253927 | 150.3453464 | 178.3805172 | 138.0084724 | -1.621084166 | 5.09E-11 | OIP5 |
| ENSG00000088325 | 1097.042938 | 1123.783767 | 1145.426243 | 3458.822179 | 3508.792984 | 3437.211011 | -1.628303281 | 4.48E-203 | TPX2 |
| ENSG00000076382 | 298.3632042 | 334.6331997 | 334.2178556 | 1012.85286 | 964.2190118 | 1023.062806 | -1.633844628 | 7.46E-66 | SPAG5 |
| ENSG00000185432 | 268.9328201 | 251.2355175 | 218.485459 | 794.8081469 | 831.1567882 | 669.0410725 | -1.634931736 | 1.99E-41 | METTL7A |
| ENSG00000084734 | 39.57879239 | 69.84555882 | 48.67250324 | 189.9099112 | 155.2392609 | 146.0089635 | -1.637562954 | 2.29E-10 | GCKR |
| ENSG00000187951 | 38.56395156 | 51.08108033 | 47.59089206 | 135.398733 | 168.7383271 | 127.007797 | -1.652650465 | 2.23E-10 | AC091057.1 |
| ENSG00000148773 | 1444.118502 | 1243.667935 | 1254.686972 | 4350.343707 | 3753.704613 | 4318.265099 | -1.655417695 | 3.05E-128 | MKI67 |
| ENSG00000144354 | 298.3632042 | 257.4903437 | 290.9534083 | 867.7827887 | 943.9704126 | 860.0527988 | -1.657543606 | 6.27E-58 | CDCA7 |
| ENSG00000284946 | 18.26713495 | 30.23165979 | 40.01961377 | 74.73306691 | 105.0998723 | 99.006078 | -1.658379274 | 2.10E-06 | AC068831.7 |
| ENSG00000109674 | 45.66783737 | 72.9729719 | 42.18283614 | 158.2582593 | 188.0227073 | 162.0099458 | -1.659063031 | 4.55E-11 | NEIL3 |
| ENSG00000118193 | 187.7455536 | 213.7065606 | 190.3635682 | 595.2268976 | 603.6011014 | 672.0412567 | -1.659877242 | 3.92E-42 | KIF14 |
| ENSG00000196584 | 92.35051557 | 80.27026909 | 87.61050583 | 287.5025045 | 265.1602282 | 278.0170675 | -1.674512544 | 3.55E-21 | XRCC2 |
| ENSG00000160298 | 76.11306229 | 85.48262422 | 90.85533938 | 285.7440794 | 229.4841248 | 293.0179884 | -1.680137456 | 9.37E-19 | C21orf58 |
| ENSG00000171241 | 368.3872215 | 327.3359025 | 373.1558582 | 1091.981989 | 1213.951736 | 1134.069621 | -1.68616796 | 2.41E-73 | SHCBP1 |
| ENSG00000108342 | 23.3413391 | 23.97683362 | 19.4690013 | 83.52519243 | 71.35220687 | 60.00368364 | -1.686584598 | 5.46E-06 | CSF3 |
| ENSG00000136492 | 77.12790312 | 101.1196896 | 97.34500648 | 274.3143162 | 314.3353979 | 300.0184182 | -1.689868714 | 2.56E-21 | BRIP1 |
| ENSG00000189184 | 20.29681661 | 14.59459438 | 9.734500648 | 43.96062759 | 60.74579774 | 40.00245576 | -1.692212011 | 0.000437776 | PCDH18 |
| ENSG00000165304 | 330.8381107 | 361.7374464 | 336.3810779 | 1099.01569 | 1164.776566 | 1063.065262 | -1.693090055 | 1.24E-75 | MELK |
| ENSG00000101057 | 700.240173 | 795.4053937 | 726.842715 | 2385.303653 | 2431.760348 | 2385.146425 | -1.696383843 | 5.41E-145 | MYBL2 |
| ENSG00000146670 | 380.5653114 | 394.0540483 | 336.3810779 | 1146.493168 | 1298.803009 | 1162.07134 | -1.698100091 | 2.58E-72 | CDCA5 |
| ENSG00000117724 | 641.3794049 | 600.4633116 | 630.5793197 | 2161.104453 | 1896.618796 | 2030.12463 | -1.701363228 | 2.84E-116 | CENPF |
| ENSG00000164619 | 8.118726644 | 12.50965233 | 10.81611183 | 35.16850207 | 50.13938861 | 17.0010437 | -1.705215518 | 0.006009485 | BMPER |
| ENSG00000071539 | 304.4522491 | 380.5019249 | 375.3190805 | 1127.150492 | 1221.665488 | 1110.068147 | -1.706914953 | 3.39E-67 | TRIP13 |
| ENSG00000072571 | 235.4430727 | 239.7683362 | 293.1166306 | 842.2856247 | 840.7989783 | 823.0505272 | -1.707298138 | 1.21E-55 | HMMR |
| ENSG00000142945 | 270.9625017 | 260.6177568 | 292.0350194 | 829.976649 | 942.0419745 | 920.0564824 | -1.708594126 | 1.76E-60 | KIF2C |
| ENSG00000168078 | 163.3893737 | 183.4749008 | 218.485459 | 612.8111487 | 631.5634527 | 607.0372661 | -1.7133407 | 1.04E-41 | PBK |
| ENSG00000198826 | 457.6932145 | 379.4594539 | 398.0329154 | 1383.001344 | 1353.763493 | 1336.082022 | -1.720659035 | 6.12E-87 | ARHGAP11A |
| ENSG00000129195 | 134.9738305 | 120.9266391 | 150.3439545 | 464.2242274 | 475.3599728 | 405.0248645 | -1.728220596 | 9.19E-32 | PIMREG |
| ENSG00000142731 | 207.0275294 | 172.0077195 | 193.6084018 | 643.583588 | 646.0267379 | 617.0378801 | -1.7351466 | 2.90E-47 | PLK4 |
| ENSG00000184661 | 185.715872 | 232.4710391 | 168.7313446 | 596.9853227 | 672.0606512 | 687.0421776 | -1.735367095 | 6.30E-40 | CDCA2 |
| ENSG00000174371 | 90.32083391 | 80.27026909 | 120.0588413 | 297.1738425 | 347.1188443 | 323.0198302 | -1.736285923 | 5.45E-22 | EXO1 |
| ENSG00000133110 | 156.2854879 | 177.2200746 | 167.6497334 | 544.2325696 | 634.4561098 | 492.0302058 | -1.737674111 | 4.12E-36 | POSTN |
| ENSG00000051341 | 97.42471973 | 95.9073345 | 80.03922755 | 307.7243932 | 295.0510176 | 313.0192163 | -1.742689837 | 3.18E-24 | POLQ |
| ENSG00000134057 | 617.0232249 | 691.158291 | 697.6392131 | 2130.332013 | 2445.259414 | 2141.131444 | -1.743990121 | 4.43E-113 | CCNB1 |
| ENSG00000189057 | 282.1257509 | 295.0193007 | 260.6682951 | 907.3473535 | 978.682297 | 935.0574033 | -1.75070045 | 4.57E-69 | FAM111B |
| ENSG00000131153 | 169.4784187 | 157.4131251 | 125.4668972 | 466.861865 | 559.2470269 | 499.0306356 | -1.751023871 | 1.45E-33 | GINS2 |
| ENSG00000156509 | 16.23745329 | 9.382239244 | 15.14255656 | 47.4774778 | 30.85500838 | 59.00362224 | -1.751550532 | 0.000516022 | FBXO43 |
| ENSG00000100526 | 104.5286055 | 128.2239363 | 120.0588413 | 375.4237597 | 387.6160428 | 426.0261538 | -1.75301127 | 4.33E-30 | CDKN3 |
| ENSG00000134690 | 250.6656851 | 302.3165979 | 270.4027958 | 920.5355418 | 979.646516 | 891.054702 | -1.761680721 | 1.11E-64 | CDCA8 |
| ENSG00000166250 | 163.3893737 | 169.9227774 | 164.4048998 | 560.0583955 | 552.4974938 | 578.0354857 | -1.764001038 | 1.09E-45 | CLMP |
| ENSG00000143228 | 155.2706471 | 145.9459438 | 178.4658452 | 473.0163529 | 589.1378162 | 577.0354243 | -1.772767759 | 3.15E-36 | NUF2 |
| ENSG00000169607 | 131.929308 | 168.8803064 | 188.2003459 | 561.8168206 | 593.9589113 | 513.0314951 | -1.773242279 | 3.34E-36 | CKAP2L |
| ENSG00000166851 | 585.5631592 | 618.1853191 | 597.0493731 | 2033.618632 | 2126.102921 | 2002.122911 | -1.774824768 | 2.51E-144 | PLK1 |
| ENSG00000138160 | 434.3518754 | 375.2895698 | 447.7870298 | 1437.512522 | 1481.040402 | 1423.087364 | -1.788076753 | 1.20E-98 | KIF11 |
| ENSG00000085999 | 56.83108651 | 47.95366725 | 77.87600518 | 230.3536886 | 210.1997446 | 189.0116035 | -1.788730646 | 1.66E-15 | RAD54L |
| ENSG00000241472 | 11.16324914 | 10.42471027 | 4.326444732 | 32.53086442 | 32.7834464 | 25.00153485 | -1.796466828 | 0.003883894 | PTPRG-AS1 |
| ENSG00000110675 | 832.169481 | 803.7451619 | 820.942888 | 2816.997016 | 3006.434879 | 2750.168833 | -1.803030589 | 2.79E-182 | ELMOD1 |
| ENSG00000139549 | 39.57879239 | 19.80694952 | 30.28511313 | 94.07574305 | 91.60080612 | 128.0078584 | -1.803526909 | 4.95E-08 | DHH |
| ENSG00000117399 | 760.115782 | 781.8532704 | 946.4097852 | 2974.376063 | 3009.327536 | 2724.167237 | -1.808314332 | 6.25E-126 | CDC20 |
| ENSG00000213347 | 64.94981315 | 79.22779806 | 81.12083873 | 314.7580936 | 242.018972 | 233.0143048 | -1.812707295 | 3.38E-19 | MXD3 |
| ENSG00000112742 | 158.3151696 | 148.0308859 | 143.8542874 | 495.8758793 | 544.7837417 | 543.0333369 | -1.813363443 | 5.08E-43 | TTK |
| ENSG00000240583 | 137.0035121 | 144.9034728 | 140.6094538 | 472.1371404 | 598.7800063 | 415.0254785 | -1.814539629 | 1.15E-30 | AQP1 |
| ENSG00000093009 | 171.5081004 | 176.1776036 | 181.7106788 | 606.6566608 | 615.1717295 | 645.0395991 | -1.818289933 | 6.95E-52 | CDC45 |
| ENSG00000144554 | 130.9144671 | 135.5212335 | 105.9978959 | 444.0023387 | 451.2544975 | 420.0257854 | -1.819021796 | 6.55E-35 | FANCD2 |
| ENSG00000112984 | 469.8713045 | 504.5559771 | 423.9915838 | 1550.930942 | 1840.694094 | 1548.095038 | -1.819900348 | 1.95E-87 | KIF20A |
| ENSG00000284906 | 13.1929308 | 14.59459438 | 12.9793342 | 65.06172884 | 31.81922739 | 47.00288551 | -1.822069447 | 0.000217439 | AC091057.6 |
| ENSG00000137804 | 44.65299654 | 43.78378314 | 34.61155786 | 132.7610953 | 147.5255088 | 157.0096388 | -1.826442738 | 1.03E-12 | NUSAP1 |
| ENSG00000170312 | 287.199955 | 283.5521194 | 255.2602392 | 892.4007402 | 1081.853731 | 958.0588154 | -1.826558192 | 5.26E-65 | CDK1 |
| ENSG00000186871 | 85.24662976 | 97.99227655 | 73.54956045 | 288.381717 | 308.5500838 | 317.0194619 | -1.829806944 | 4.87E-25 | ERCC6L |
| ENSG00000007968 | 34.50458824 | 37.52895698 | 49.75411442 | 147.7077087 | 140.7759757 | 144.0088407 | -1.831416329 | 1.35E-12 | E2F2 |
| ENSG00000138587 | 26.38586159 | 13.55212335 | 23.79544603 | 51.87354056 | 82.92283502 | 93.00570964 | -1.834401808 | 4.80E-06 | MNS1 |
| ENSG00000165071 | 38.56395156 | 31.27413081 | 36.77478023 | 88.80046774 | 167.7741081 | 126.0077356 | -1.841251387 | 2.87E-09 | TMEM71 |
| ENSG00000135547 | 37.54911073 | 31.27413081 | 55.16217034 | 147.7077087 | 152.3466039 | 146.0089635 | -1.850384021 | 2.54E-12 | HEY2 |
| ENSG00000146678 | 14.20777163 | 9.382239244 | 9.734500648 | 35.16850207 | 43.38985553 | 42.00257854 | -1.850932195 | 0.000335234 | IGFBP1 |
| ENSG00000154839 | 153.2409654 | 160.5405382 | 177.384234 | 579.4010717 | 621.9212626 | 582.0357313 | -1.861284644 | 1.03E-49 | SKA1 |
| ENSG00000105011 | 166.4338962 | 183.4749008 | 221.7302925 | 673.4768147 | 768.4825524 | 636.0390465 | -1.863863549 | 1.89E-46 | ASF1B |
| ENSG00000075218 | 293.289 | 359.6525044 | 350.4420233 | 1211.554896 | 1166.705004 | 1274.078216 | -1.864800145 | 2.76E-85 | GTSE1 |
| ENSG00000197299 | 82.20210727 | 71.93050087 | 84.36567228 | 276.9519538 | 289.2657035 | 308.0189093 | -1.873743307 | 1.06E-25 | BLM |
| ENSG00000151725 | 159.3300104 | 121.9691102 | 146.0175097 | 496.7550918 | 561.1754649 | 511.0313723 | -1.875644841 | 5.34E-41 | CENPU |
| ENSG00000041353 | 6.089044983 | 16.67953643 | 11.89772301 | 47.4774778 | 48.21095059 | 32.00196461 | -1.885799197 | 0.000389578 | RAB27B |
| ENSG00000024526 | 200.9384844 | 232.4710391 | 204.4245136 | 742.9346063 | 816.693503 | 804.0493607 | -1.889295013 | 1.00E-63 | DEPDC1 |

| | | | | | | | | | |
|---|---|---|---|---|---|---|---|---|---|
| ENSG00000121152 | 179.626827 | 178.2625456 | 139.5278426 | 605.7774482 | 641.2056429 | 606.0372047 | -1.8954937 | 2.56E-49 | NCAPH |
| ENSG00000092853 | 181.6565087 | 241.8532783 | 191.4451794 | 722.7127176 | 795.4806847 | 771.0473347 | -1.896065328 | 2.52E-55 | CLSPN |
| ENSG00000183856 | 243.5617993 | 251.2355175 | 244.4441274 | 995.2686087 | 894.795243 | 872.0535355 | -1.902419363 | 9.33E-74 | IQGAP3 |
| ENSG00000127564 | 158.3151696 | 137.6061756 | 147.0991209 | 554.7831202 | 544.7837417 | 559.0343192 | -1.904016388 | 4.07E-49 | PKMYT1 |
| ENSG00000109805 | 399.8472872 | 407.6061716 | 401.2777489 | 1445.425435 | 1586.140274 | 1536.094301 | -1.917578025 | 6.34E-122 | NCAPG |
| ENSG00000134222 | 56.83108651 | 71.93050087 | 99.50822885 | 291.8985672 | 314.3353979 | 255.0156555 | -1.919705245 | 6.38E-21 | PSRC1 |
| ENSG00000157456 | 228.3391869 | 248.1081045 | 213.0774031 | 809.7547603 | 989.2887061 | 825.05065 | -1.927343967 | 1.91E-61 | CCNB2 |
| ENSG00000196550 | 27.40070242 | 27.10424671 | 23.79544603 | 116.0560568 | 96.42190118 | 86.00527988 | -1.931311972 | 1.54E-09 | FAM72A |
| ENSG00000186185 | 258.7844118 | 256.4478727 | 271.484407 | 1032.195536 | 979.646516 | 995.061087 | -1.935016668 | 2.55E-88 | KIF18B |
| ENSG00000123485 | 183.6861903 | 181.3899587 | 203.3429024 | 727.1087804 | 746.3055151 | 700.0429757 | -1.935936073 | 4.84E-64 | HJURP |
| ENSG00000171848 | 421.1589446 | 433.6679473 | 412.0938608 | 1668.745423 | 1644.957634 | 1560.095775 | -1.943859504 | 1.05E-134 | RRM2 |
| ENSG00000188610 | 38.56395156 | 30.23165979 | 33.52994668 | 140.6740083 | 134.9906617 | 119.0073059 | -1.947401602 | 1.06E-12 | FAM72B |
| ENSG00000091651 | 103.5137647 | 140.7335887 | 128.7117308 | 483.5669035 | 513.9287333 | 452.0277501 | -1.959967126 | 4.77E-40 | ORC6 |
| ENSG00000169679 | 320.6897024 | 348.1853231 | 316.9120766 | 1192.21222 | 1343.157083 | 1306.08018 | -1.961631008 | 1.96E-101 | BUB1 |
| ENSG00000152253 | 34.50458824 | 72.9729719 | 46.50928087 | 232.9913262 | 179.3447362 | 187.0114807 | -1.962726052 | 2.08E-14 | SPC25 |
| ENSG00000163808 | 137.0035121 | 114.671813 | 117.895619 | 448.3984015 | 475.3599728 | 523.032109 | -1.96708437 | 8.36E-42 | KIF15 |
| ENSG00000156970 | 271.9773426 | 227.2586839 | 228.2199596 | 961.8585317 | 1024.96481 | 878.0539039 | -1.976682089 | 9.25E-73 | BUB1B |
| ENSG00000135451 | 142.0777163 | 150.1158279 | 178.4658452 | 640.9459503 | 638.3129858 | 573.0351787 | -1.978749113 | 4.63E-52 | TROAP |
| ENSG00000145386 | 378.5356298 | 478.4942015 | 465.0928087 | 1694.242587 | 1887.940825 | 1652.101423 | -1.986092034 | 4.88E-106 | CCNA2 |
| ENSG00000237649 | 255.7398893 | 265.8301119 | 288.7901859 | 1085.827502 | 1019.179495 | 1126.06913 | -1.996200649 | 3.70E-92 | KIFC1 |
| ENSG00000182010 | 9.133567474 | 9.382239244 | 16.22416775 | 48.35669035 | 33.74766541 | 57.00349945 | -2.005965865 | 7.52E-05 | RTKN2 |
| ENSG00000133101 | 50.74204152 | 78.18532704 | 67.05989335 | 269.9182534 | 269.9813233 | 251.0154099 | -2.01471853 | 5.50E-24 | CCNA1 |
| ENSG00000065328 | 156.2854879 | 136.5637046 | 148.1807321 | 618.086424 | 646.0267379 | 536.0329071 | -2.029377669 | 5.66E-52 | MCM10 |
| ENSG00000261253 | 5.074204152 | 5.212355136 | 6.489667099 | 34.28928952 | 13.49906617 | 21.00128927 | -2.041910033 | 0.007243622 | AC137932.2 |
| ENSG00000132470 | 63.93497232 | 64.63320368 | 46.50928087 | 237.387389 | 284.4446085 | 207.0127085 | -2.055649493 | 3.50E-21 | ITGB4 |
| ENSG00000178999 | 172.5229412 | 187.6447849 | 216.3222366 | 827.3390113 | 730.878011 | 836.0513253 | -2.055940048 | 7.27E-67 | AURKB |
| ENSG00000138180 | 426.2331488 | 394.0540483 | 465.0928087 | 1729.41109 | 1883.11973 | 1732.106334 | -2.056468113 | 4.70E-139 | CEP55 |
| ENSG00000137812 | 319.6748616 | 286.6795325 | 247.6889609 | 1156.164506 | 1156.098595 | 1255.077049 | -2.060475827 | 6.57E-94 | KNL1 |
| ENSG00000175063 | 210.0720519 | 233.5135101 | 235.7912379 | 879.2125519 | 973.8612019 | 993.0609642 | -2.066852508 | 5.31E-84 | UBE2C |
| ENSG00000080986 | 176.5823045 | 134.4787625 | 184.9555123 | 689.3026407 | 712.5578497 | 741.0454929 | -2.111340651 | 2.65E-62 | NDC80 |
| ENSG00000115163 | 64.94981315 | 79.22779806 | 57.3253927 | 283.9856543 | 312.4069598 | 288.0176815 | -2.132589516 | 9.37E-30 | CENPA |
| ENSG00000185862 | 7.103885813 | 6.254826163 | 11.89772301 | 23.7387389 | 46.28251257 | 41.00251715 | -2.137821646 | 0.000281166 | EVI2B |
| ENSG00000158402 | 43.63815571 | 38.571428 | 36.77478023 | 175.8425104 | 158.1319179 | 200.0122788 | -2.164169137 | 6.52E-19 | CDC25C |
| ENSG00000126787 | 447.5448062 | 457.6447809 | 432.6444732 | 1935.146827 | 2144.423082 | 2028.124507 | -2.190154394 | 2.79E-182 | DLGAP5 |
| ENSG00000131747 | 973.2323564 | 888.1853151 | 996.1638996 | 4205.273636 | 4578.111868 | 4267.261968 | -2.191338261 | 1.14E-298 | TOP2A |
| ENSG00000161888 | 72.05369896 | 77.14285601 | 88.69211701 | 351.6850207 | 360.6179104 | 386.0236981 | -2.208205479 | 1.86E-39 | SPC24 |
| ENSG00000066279 | 440.4409204 | 395.0965193 | 362.3397463 | 1868.326673 | 1782.840953 | 1917.117692 | -2.215378063 | 3.03E-160 | ASPM |
| ENSG00000261008 | 4.059363322 | 2.084942054 | 6.489667099 | 15.82582593 | 21.21281826 | 23.00141206 | -2.251918703 | 0.005858025 | LINC01572 |
| ENSG00000239556 | 5.074204152 | 3.127413081 | 3.244833549 | 21.9803138 | 22.17703727 | 11.00067533 | -2.267798401 | 0.008721522 | AC004951.2 |
| ENSG00000065618 | 32.47490658 | 39.61389903 | 36.77478023 | 161.7751095 | 185.1300503 | 182.0111737 | -2.280617733 | 1.41E-20 | COL17A1 |
| ENSG00000121211 | 14.20777163 | 8.339768217 | 10.81611183 | 54.51117822 | 65.5668928 | 46.00282412 | -2.313726794 | 6.28E-07 | MND1 |
| ENSG00000256612 | 5.074204152 | 4.169884109 | 7.571278282 | 22.85952635 | 35.67610344 | 25.00153485 | -2.31567195 | 0.000643805 | CYP2B7P |
| ENSG00000171346 | 15.22261246 | 29.18918876 | 24.87705721 | 123.9689698 | 119.5631575 | 108.0066305 | -2.346801956 | 1.07E-13 | KRT15 |
| ENSG00000123689 | 84.23178893 | 123.0115812 | 94.10017293 | 487.9629663 | 536.1057706 | 522.0320476 | -2.359252159 | 1.43E-53 | G0S2 |
| ENSG00000102900 | 316.6303391 | 345.05791 | 296.3614642 | 1633.576921 | 1660.385138 | 1647.101116 | -2.365988755 | 9.58E-176 | NUP93 |
| ENSG00000163814 | 16.23745329 | 15.63706541 | 28.12189076 | 121.3313322 | 107.9925293 | 91.00558685 | -2.423230072 | 1.15E-12 | CDCP1 |
| ENSG00000035499 | 56.83108651 | 56.29343547 | 71.38633808 | 285.7440794 | 333.6197781 | 379.0232683 | -2.436558572 | 2.56E-36 | DEPDC1B |
| ENSG00000179750 | 5.074204152 | 14.59459438 | 14.06094538 | 61.54487863 | 63.63845478 | 63.00386782 | -2.486398893 | 3.80E-08 | APOBEC3B |
| ENSG00000163072 | 1.01484083 | 4.169884109 | 6.489667099 | 14.94661338 | 38.56876047 | 17.0010437 | -2.604548365 | 0.001975128 | NOSTRIN |
| ENSG00000099937 | 137.0035121 | 126.1389943 | 141.691065 | 1221.226235 | 1147.420624 | 1138.069866 | -3.115538025 | 3.08E-175 | SERPIND1 |
| ENSG00000187513 | 7.103885813 | 17.72200746 | 5.408055916 | 236.5081765 | 296.9794556 | 204.0125244 | -4.605212788 | 1.76E-40 | GJA4 |
| ENSG00000231066 | 0 | 0 | 0 | 1.758425104 | 8.677971106 | 9.000552545 | -5.068353394 | 0.0099406 | NPM1P9 |
| ENSG00000224715 | 0 | 0 | 0 | 7.033700415 | 5.785314071 | 7.000429757 | -5.102010875 | 0.006763631 | Z82186.1 |
